# Explainable Deep Relational Networks for Predicting Compound-Protein Affinities and Contacts

**DOI:** 10.1101/2019.12.28.890103

**Authors:** Mostafa Karimi, Di Wu, Zhangyang Wang, Yang Shen

## Abstract

Predicting compound-protein affinity is beneficial for accelerating drug discovery. Doing so without the often-unavailable structure data is gaining interest. However, recent progress in structure-free affinity prediction, made by machine learning, focuses on accuracy but leaves much to be desired for interpretability. Defining inter-molecular contacts underlying affinities as a vehicle for interpretability, our large-scale interpretability assessment finds previously-used attention mechanisms inadequate. We thus formulate a hierarchical multi-objective learning problem whose predicted contacts form the basis for predicted affinities. And we solve the problem by embedding protein sequences (by hierarchical recurrent neural networks) and compound graphs (by graph neural networks) with joint attentions between protein residues and compound atoms. We further introduce three methodological advances to enhance interpretability: (1) structure-aware regularization of attentions using protein sequence-predicted solvent exposure and residue-residue contact maps; (2) supervision of attentions using known inter-molecular contacts in training data; and (3) an intrinsically explainable architecture where atomic-level contacts or “relations” lead to molecular-level affinity prediction. The first two and all three advances result in DeepAffinity+ and DeepRelations, respectively. Our methods show generalizability in affinity prediction for molecules that are new and dissimilar to training examples. Moreover, they show superior interpretability compared to state-of-the-art interpretable methods: with similar or better affinity prediction, they boost the AUPRC of contact prediction by around 33, 35, 10, and 9-fold for the default test, new-compound, new-protein, and both-new sets, respectively. We further demonstrate their potential utilities in contact-assisted docking, structure-free binding site prediction, and structure-activity relationship studies without docking. Our study represents the first model development and systematic model assessment dedicated to interpretable machine learning for structure-free compound-protein affinity prediction.

## Introduction

Current drug-target interactions are predominantly represented by the interactions between small-molecule compounds as drugs and proteins as targets.^1^ The enormous chemical space to screen compounds is estimated to contain 10^60^ drug-like compounds.^2^ And these compounds act in biological systems of millions or more protein species or “proteoforms” (considering genetic mutations, alternative splicing, and post-translation modifications of proteins).^3,4^ Facing such a combinatorial explosion of compound-protein pairs, drug discovery calls for efficient characterization of compound efficacy and toxicity, and computational prediction of compound-protein interactions (CPI) addresses the need.

Classical physics-driven methods model atomic-level energetics using co-crystallized or docked 3D structures of compound-protein pairs,^5,6^ such as molecular mechanical and quantum mechanical force fields, potentials of mean force, and empirical and statistical scoring. Over the years of development, these methods are increasingly accurate^7–9^ for applications including quantitative structure-activity relationship (QSAR). Moreover, their affinity predictions are intrinsically interpretable toward revealing mechanistic principles, with the consideration of atomic contacts, dynamics, and energetics as well as solvent effects. Recently, thanks to increasingly abundant molecular data and advanced computing power, data-driven machine learning (especially deep learning) methods are also developed using the input structures of compound-protein complexes^10–12^ or proteins alone (see a related task of classifying binding^13,14^), albeit with less focus on interpretability. However, these structure-based methods, physics- or data-driven, are limited by the availability of structure data. Indeed, 3D structures are often not available for compound-protein pairs or even proteins alone and their prediction through docking is still a computationally demanding and challenging task.

To overcome the data limitation of structure-based affinity-prediction methods and broaden the applicability to more chemical-proteomic pairs without structures, our focus of the study is structure-free prediction of compound-protein affinities. Recent developments only use identities of compounds (SMILES^15,16^ or graphs^16,17^) and proteins (amino acid sequences^15,17^ or shorter, predicted structural property sequences^16^) as inputs. Compared to these recent work, our goals are two folds: improved generalizability to “new” molecules unseen in training data as well as improved interpretability to a level that data supports (not yet the level of mechanical principles that can be revealed by physics-driven structure-based methods). In particular, interpretability remains a major gap between the capability of current structure-free machine-learning models and the demand for rational drug discovery. The central question about interpretability is whether and how methods (including machine learning models) could explain *why* they make certain predictions (affinity level for any compound-protein pair in our context). This important topic is rarely addressed in structure-free machine learning models. DeepAffinity^16^ has embedded joint attentions over compound-protein component pairs and uses such joint attentions to assess origins of affinities (binding sites) or specificities. Additionally, attention mechanisms have been used for predictions of CPI,^18^ chemical stability^19^ and protein secondary structures. ^20^ Assessment of interpretability for all these studies was either lacking or limited to a few case studies. We note a recent work proposing *post-hoc* attribution-based test to determine whether a model learns binding mechanisms.^21^

We raise reasonable concerns on how much attention mechanisms can reproduce native contacts in compound-protein interactions. Attention mechanisms were originally developed to boost the performance of seq2seq models for neural machine translations.^22^ And they have gained popularity for interpreting deep learning models in visual question answering, ^23^ natural language processing, ^24^ and healthcare. ^25^ However, they were also found to work differently from human attentions in visual question answering. ^26^

Representing the first effort dedicated to the interpretability of structure-free compound-protein affinity predictors (in particular, deep-learning models), our study is focused on how to define, assess, and enhance interpretability for these methods as follows.

### How to define interpretability for affinity prediction

Interpretable machine learning is increasingly becoming a necessity^27^ for fields beyond drug discovery. Unlike interpretability in a generic case, ^27^ what interpretability actually means and how it should be evaluated is much less ambiguous for compound-protein affinity prediction. So that explanations conform with scientific knowledge, human understanding, and drug-discovery needs, we define interpretability of affinity prediction as to the ability to explain predicted affinity through underlying atomic interactions (or contacts). Specifically, atomic contacts of various types are known to constitute the physical basis of intermolecular interactions, ^28^ modeled in force fields to estimate interaction energies, ^6^ needed to explain mechanisms of actions for drugs, ^29,30^ and relied upon to guide structure-activity research in drug discovery. ^31,32^ Therefore we use the ability to replicate such corresponding contacts *while* predicting affinities as a vehicle for interpretability. The current definition of interpretability (residue-atom pairs in contact) is primitive compared to mechanistic principles in structure-based classical methods. But it is expected to serve as a vehicle to help fill the mechanistic void in structure-free affinity predictors (especially deep-learning models). We emphasize that simultaneous prediction of affinity and contacts does not necessarily make the affinity predictors intrinsically interpretable unless predicted contacts form the basis for predicted affinities.

### How to assess interpretability for affinity prediction

Once interpretability of affinity predictors is defined first through atomic contacts, it can be readily assessed against ground truth known in compound-protein structures, which overcomes the barrier for interpretable machine learning without ground truth.^33^ In our study, we have curated a dataset of compound-protein pairs, all of which are labeled with *K_d_* values and some of which with contact details; and we have split them into training, test, compound-unique, protein-unique, and both-unique (or double-unique) sets. We measure the accuracy of contact prediction over various sets using area under the precision-recall curve (AUPRC) which is suitable for binary classification (contacts/non-contacts) with highly imbalanced classes (far fewer contacts than non-contacts). We have performed large-scale assessments of attention mechanisms in various molecular data representations (protein amino-acid sequences and structure-property annotated sequences^16^ as well as compound SMILES and graphs) and corresponding neural network architectures (convolutional and recurrent neural networks [CNN and RNN] as well as graph convolutional and isomorphism networks [GCN and GIN]). And we have found that current attention mechanisms inadequate for interpretable affinity prediction, as their AUPRCs were merely slightly more than chance (0.004).

### How to enhance interpretability for affinity prediction

We have made three main contributions to enhance interpretability for structure-free deep-learning models.

The first contribution is to incorporate physical constraints into data representations, model architectures, and model training. (1) To respect the sequence nature of protein inputs and to overcome the computational bottlenecks of RNNs, inspired by protein folding principles, we represent protein sequences as hierarchical *k*-mers and model them with hierarchical attention networks (HANs). (2) To respect the structural contexts of proteins, we predict from protein sequences solvent exposure over residues and contact maps over residue pairs; and we introduce novel structure-aware regularizations for structured sparsity of model attentions.

The second contribution is to supervise attentions with native intermolecular contacts available to training data and to accordingly teach models how to pay attention to pairs of compound atoms and protein residues while making affinity predictions. We have formulated a hierarchical multi-objective optimization problem where contact predictions form the basis for affinity prediction. We utilize contact data available to training compound-protein pairs and design hierarchical training strategies accordingly.

The last contribution is to design intrinsic explainability into the architecture of a deep “relational” network. Inspired by physics, we explicitly model and learn various types of atomic interactions (or “relations”) through deep neural networks with joint attentions embedded. This was motivated by relational neural networks first introduced to learn to reason in computer vision^34,35^ and subsequent interaction networks to learn the relations and interactions of complex objects and their dynamics. ^36,37^ Moreover, we combine such deep relational modules in a hierarchy to progressively focus attention from putative protein surfaces, binding-site k-mers and residues, to putative residue-atom binding pairs.

The rest of the paper is organized as follows. The aforementioned contributions in defining, measuring, and enhancing interpretable affinity prediction will be detailed in Methods. In Results, we first show over established affinity-benchmark datasets that the original Deep-Affinity^16^ and its variants (with various molecular representations and neural networks) have comparable or better accuracy in affinity prediction, compared to current non-interpretable structure-free methods. We then describe a dataset newly curated for both affinity and contact prediction. The dataset is designed to be diverse and challenging with the generalizability test in mind. Using this dataset, we incrementally introduce the three contributions to DeepAffinity and compare the resulting DeepAffinity+ (using the first two contributions) and DeepRelations (using all three contributions) to a competing interpretable method. Both methods produce remarkably improved interpretability (now defined as accuracy of contacts predicted by joint attentions) while maintaining accurate and generalizable affinity prediction. Importantly, compared to the competing method and their reduced version without supervising attentions, they show that sufficiently better interpretability (much more accurate contact predictions) can help improve accuracy in affinity prediction. Lastly, we use various focused studies to show the spatial patterns of top-10 predicted contacts, the benefit of these predictions to contact-assisted protein-ligand docking, and the additional utilities of aggregating attentions and decomposing predicted affinities for binding site prediction and QSAR.

## Methods

Toward genome-wide prediction of compound-protein interactions (CPI), we assume that proteins are only available in 1D amino-acid sequences, whereas compounds are available in 1D SMILES or 2D chemical graphs. We start the section with the curation of a dataset of compound-protein pairs with known p*K_d_*/p*K_i_* values, which is also of known intermolecular contacts. We will introduce the state-of-the-art and our newly-adopted neural networks to predict from such molecular data. These neural networks will be first adopted in our previous framework of DeepAffinity^16^ (supervised learning with joint attention) so that the interpretability of attention mechanisms can be systematically assessed in CPI prediction. We will then describe our physics-inspired, intrinsically explainable architecture of deep relational networks where aforementioned neural networks are used as basis models. With carefully designed regularization terms, we will explain multi-stage deep relational networks that increasingly focus attention on putative binding-site k-mers, binding-site residues, and residue-atom interactions, for the prediction and interpretation of compound-protein affinity. We will also explain how the resulting model can be trained strategically.

### Benchmark Set with Compound-Protein Affinities and Contacts

We have previously curated affinity-labeled compound-protein pairs^16^ based on BindingDB.^38^ In this study, we used those p*K_i_*/p*K_d_*-labeled data with amino-acid sequence length no more than 1,000 and curated a subset with known complex-protein co-crystal structures. We further merge the data with the refined set of PDBbind (v. 2019),^39^ leading to 4,446 pairs between 3,672 compounds and 1,287 proteins. More details about procedures are provided in the supplemental Sec. 1.1. Resulting data characteristics, including compound property distributions and protein class statistics, are described in Results.

The compound data are in the format of canonical SMILES as provided in PubChem^40^ and the protein data are in the format of FASTA sequences (UniProt canonical). Compound SMILES were also converted to graphs with RDKit. ^41^ Ionization states of compounds defined in PubChem were validated using the software OpenBabel and the compounds were further sanitized and standardized using “chem.SanitizeMol()” in the software RDKit. More details are provided in the supplemental Sec. 1.2. Atomic-level intermolecular contacts (or “relations”) were derived from compound-protein co-crystal structures in PDB, ^42^ as ground truth for the interpretablity of affinity prediction. Specifically, we cross-referenced aforementioned compound-protein pairs in PDBsum^43^ and used its LigPlot service to collect high-resolution atomic contacts or relations. These direct, first-shell contacts are given in the form of contact types (hydrogen bond or hydrophobic contact), atomic pairs, and atomic distances.

The dataset was randomly split into four folds where fold 1 did not overlap with fold 2 in compounds, did not do so with fold 3 in proteins, and did not do so with fold 4 in either compounds or proteins. Folds 2, 3, and 4 are referred to as new-compound, new-protein, and both-new sets for generalizability tests; and they contain 521, 795 and 205 pairs, respectively. Fold 1 was randomly split into training (2,334) and test (591) sets. More procedural details about data splitting are summarized in the supplemental Algorithm 1. The split of the whole dataset is illustrated in Figure 1 below. And the similarity profiles between training molecules and those in the test and generalization sets are analyzed in Results later.

**Figure 1:**
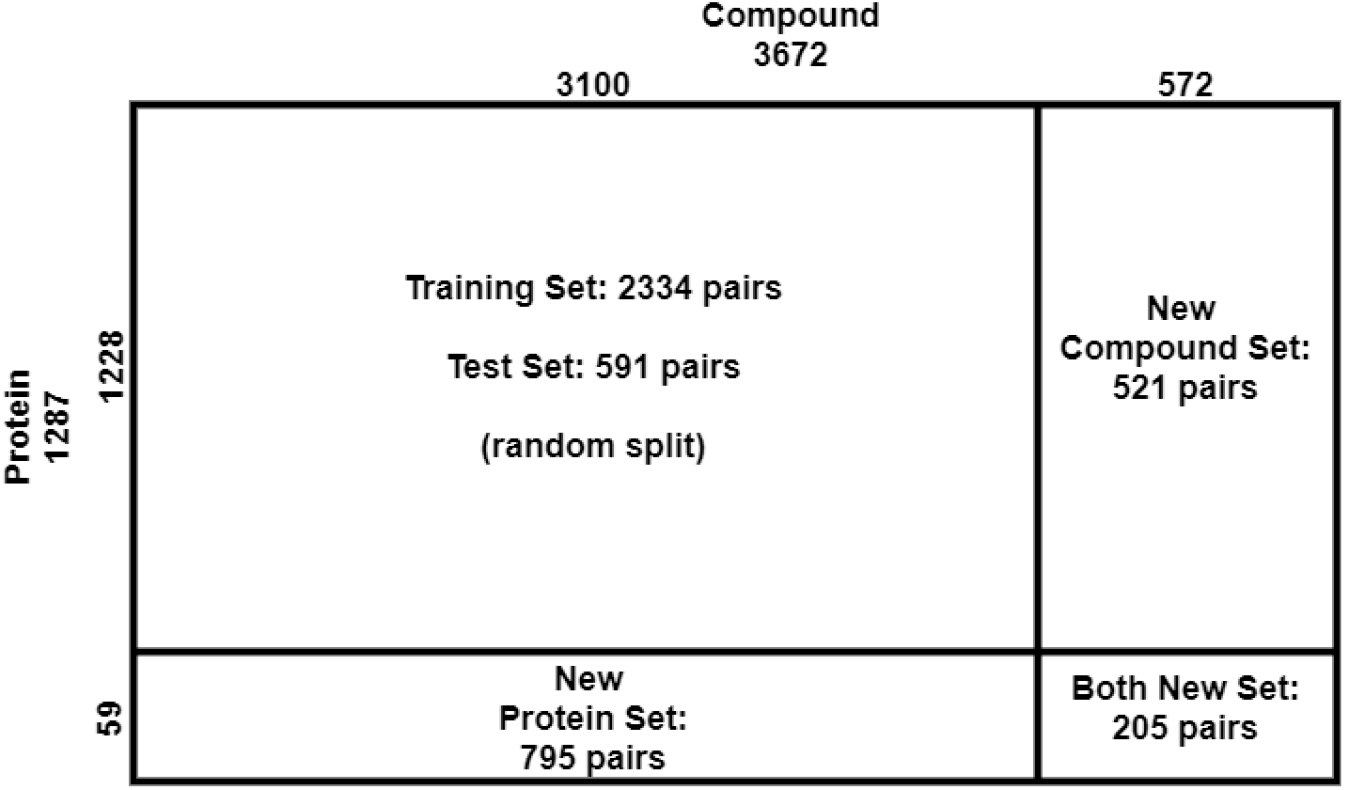
The complete data set consists of training, test, compound-unique, protein-unique, and double unique sets with compound-protein counts provided.

Although monomer structures of proteins are often unavailable, their structural features can be predicted from protein sequences alone with reasonable accuracy. We have predicted the secondary structure and solvent accessibility of each residue using the latest SCRATCH^44,45^ and contact maps for residue pairs using RaptorX-contact^46^ (see details in the supplemental Sec. 1.3). These data provide additional structural information to regularize our machine learning models. If protein structures are available, actual rather than predicted such data can be used instead.

### Data Representation and Corresponding Basis Neural Networks

#### Baseline: CNN and RNN for 1D protein and compound sequences

When molecular data are given in 1D sequences, these inputs are often processed by convolutional neural networks (CNN)^15,47^ and by recurrent neural networks (RNN) that are more suitable for sequence data with long-term interactions. ^16^

Challenges remain in RNN for compound strings or protein sequences. For compounds in SMILES strings, the descriptive power of such strings can be limited. In this study, we overcome the challenge by representing compounds in chemical formulae (2D graphs) and using two types of graph neural networks (GNN). For proteins in amino-acid sequences, the often-large lengths demand deep RNNs that are hard to be trained effectively (gradient vanishing or exploding and non-parallel training).^48^ We previously overcame the second challenge by predicting structure properties from amino-acid sequences and representing proteins as a much shorter structure property sequences where each 4-letter tuple corresponds to a secondary structure. ^16^ This treatment however limits the resolution of interpretability to be at the level of protein secondary structures (multiple neighboring residues) rather than individual residues. In this study, we overcome the second challenge while achieving residue-level interpretability by using biologically-motivated hierarchical RNN (HRNN).

##### Notation summary

Scalars, vectors, and matrices are denoted in normal lower-case, bold-faced lower-case, and upper-case characters, respectively. Subscripts *i, t*, and *j* are for the *i*^th^ protein residue, *t*^th^ protein *k*-mer and *j*^th^ compound atom, respectively. And subscript it represent the *i*^th^ residue in the *t*^th^ *k*-mer (where *i* can be regarded as a global residue index). Therefore, the *j*^th^ atom of compound 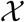 described in *d_g_* features is denoted **X**_*j*_ and its learned representation (embedded through GNN) is denoted **z**_*j*_. The *i*^th^ residue of protein 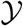 with *d_p_* features is denoted by **y**_*i*_ and its learned representation (embedded through HRNN) is denoted **h**_*it*_ where *t* is the index of the *k*-mer containing residue *i*. These residue representations **h**_*it*_ within the k-mer are then aggregated to obtain the *k*-mer representation **h**_*t*_ and all *k*-mer representations are concatenated to reach the protein representation.

Superscript *r*, (*l*), and [*s*] indicate the *r*^th^ relation about molecular features, the *l*^th^ layer of graph neural networks, and the *s*^th^ stage of DeepRelations, respectively.

#### Proposed: GCN and GIN for 2D compound graphs

Compared to 1D SMILES strings, chemical formulae (2D graphs) of compounds have more descriptive power and are increasingly used as inputs to predictive models.^16–19, 49^ In this study, compounds are represented as 2D graphs in which vertices are atoms and edges are covalent bonds between atoms. Suppose that *n* is the maximum number of atoms in our compound set (compounds with smaller number of atoms are padded to reach size n). Let’s consider a graph 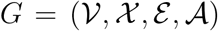, where 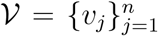 is the set of *n* vertices (each with *d_g_* features), 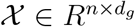 that of vertex features 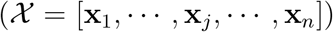, 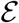 that of edges, and 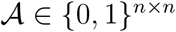 is unweighted symmetric adjacency matrix. Let 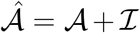 and 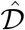 be the degree matrix (the diagonals of 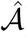).

We used Graph Convolutional Network (GCN)^50^ and Graph Isomorphism Network (GIN)^51^ which are the state of the art for graph embedding and inference. GCN consists of multiple layers and at layer *l* the model can be written as:

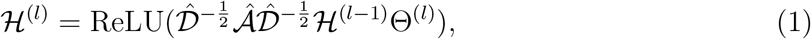

where 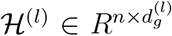 is the output, 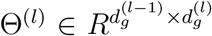 the trainable parameters, and 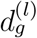 the number of features, all at layer *l*. Initial conditions (when *l* = 0) are 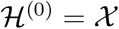 and 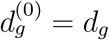.

GIN is the most powerful graph neural network in theory: its discriminative or representational power is equal to that of the Weisfeiler-Lehman graph isomorphism test.^52^ Similar to GCN, GIN consists of multiple layers and at layer *l* the model can be written as a multi-layer perceptron (MLP):

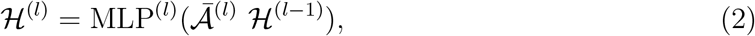

where 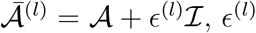 can be either a trainable parameter or a fixed hyper-parameter. Each GIN layer has several nonlinear layers compared to GCN layer with just a ReLU per layer, which might improve predictions but suffer in interpretability.

The final representation for a compound is 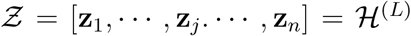 if GCN or GIN has *L* layers. In this study, vertex features are as in,^19^ with few additional features detailed later in physics-inspired relational modules. A summary of these features is provided in the supplemental Table S2.

#### Proposed: HRNN for 1D protein sequences

We aim to keep the use of RNN that respects the sequence nature of protein data and mitigate the difficulty of training RNN for long sequences. To that end, inspired by the hierarchy of protein structures, we model protein sequences using hierarchical attention networks (HANs). Specifically, during protein folding, sequence segments may fold separately into secondary structures and the secondary structures can then collectively pack into a tertiary structure needed for protein functions. We exploit such hierarchical nature by representing a protein sequence of length easily in thousands as tens or hundreds of k-mers (consecutive sequence segments) of length *k* (hyperparameter in this study). Accordingly we process the hierarchical data with hierarchical attention networks (HANs)^53^ which have been proposed for natural language processing. We also refer to it as hierarchical RNN (HRNN). Although the inter-*k*-mer attentions might overcome potential issues brought by *k*-mer definition as they do in natural language processing, ^53^ it would be interesting to examine the potential benefit of using other domain-relevant definition of *k*-mers, such as (predicted or actual) secondary structure elements.

Given 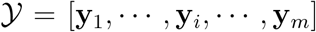, a protein sequence described with *d_p_* features for each residue 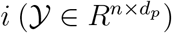, we partition it into *T* consecutive, non-overlapping *k*-mers. We use two types of RNNs in hierarchy for modeling within and across *k*-mers. We first use an embedding layer to represent the *i*^th^ residue in the *t*^th^ *k*-mer as a vector **e**_*it*_. And we use a shared RNN for all *k*-mers for the latent representation of the residue: **h**_*it*_ = RNN(**e**_*it*_) (*t* = 1,…, *T*). We then summarize each *k*-mer as **k**_*t*_ with an intra-*k*-mer attention mechanism:

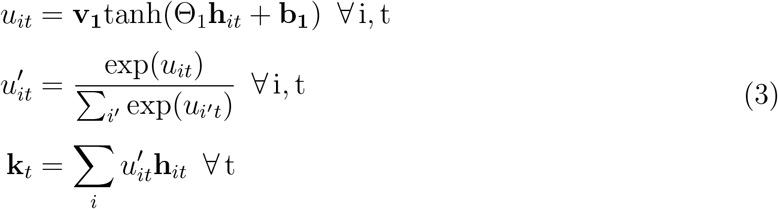

With another RNN for **k**_*t*_ we reach the representation of the *t*^th^ *k*-mer: **h**_*t*_ = RNN(**k**_*t*_) (*t* = 1,…, *T*).

The final representation for a protein sequence is the collection of **h**_*t*_.

#### Joint attention over protein-compound atomic pairs for interpretability

Once the learned representation of protein sequences (**H** = [**h**_1_,…, **h**_*t*_,…, **h**_*T*_] where *t* is the index of protein *k*-mer) and that of compound sequences or graphs (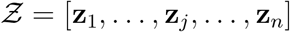 where *j* is the index of compound atom) are defined, they are processed with a joint *k*-mer–atom attention mechanism to interpret any downstream prediction:

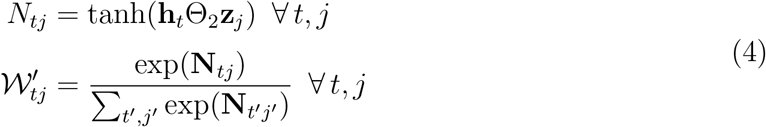

With 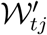, the joint attention between the *t*^th^ *k*-mer and the *j*^th^ atom, we can combine it with the intra-*k*-mer attention over each residue *i* in the *t*^th^ *k*-mer and reach 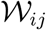, the joint attention between the *i*^th^ protein residue and the *j*^th^ compound atom:

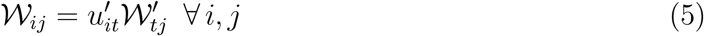

This joint attention mechanism is an extension of our previous work^16^ where a protein sequence was represented as a single, “flat” RNN rather than multiple, hierarchical RNNs.

Given learned representations **h**_*i*_ for protein residue *i* (the *k*-mer index is ignored for simplicity) and **Z**_*j*_ for compound atom *j* as well as the joint attention 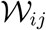 over the pair, we further jointly embed the pair and aggregate over all pairs to reach **f** — the joint embedding of protein 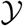, compound 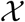, and their residue-atom “interactions” captured by 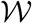:

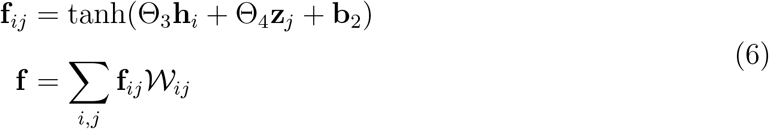

where Θ_3_, Θ_4_ and **b**_2_ are learnable parameters. The joint embedding **f** is fed to a CNN and two multi-layer perceptrons (MLP) to make affinity prediction as before.^16^ In other words, 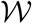 for contact prediction directly forms the basis of **f** for affinity prediction.

In comparison, Gao et al.’s method^18^ also uses joint attention for contact prediction. But the joint attention matrix is marginalized for either the compound or the protein; and the separately processed compound or protein representations were used for affinity prediction. More specifically,

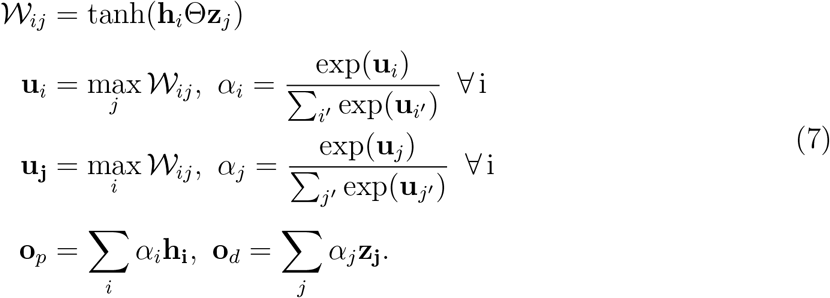

The separate final representations for the compound (**o**_*d*_) and the protein (**o**_*p*_) were fed to downstream layers for affinity prediction, with much of information lost on the joint attention (the basis of contact prediction).

### DeepRelations

#### Overall architecture

We have developed an end-to-end “by-design” interpretable architecture named DeepRelations for joint prediction and interpretation of compound-protein affinity. The overall architecture is shown in Figure 2.

**Figure 2:**
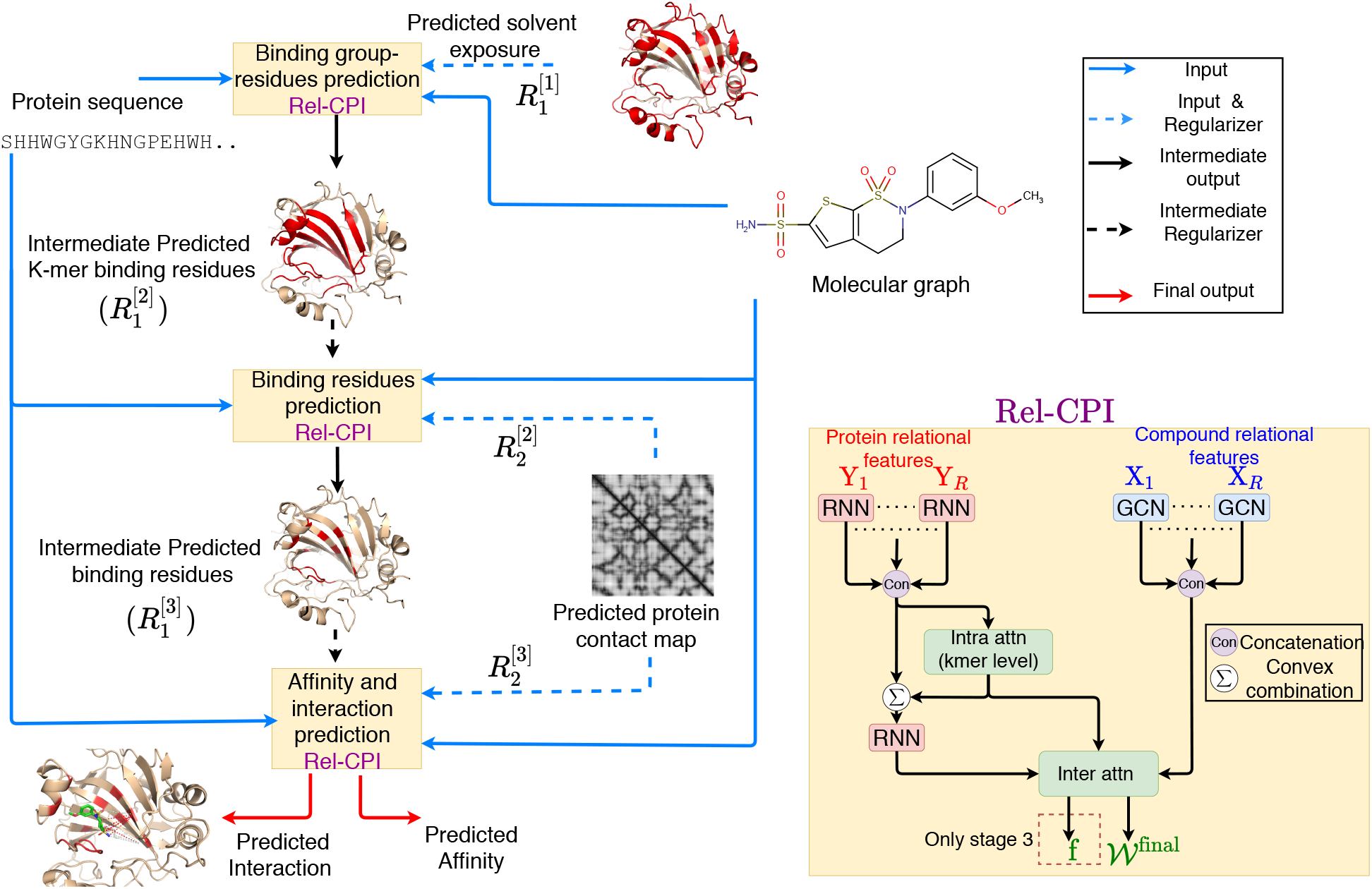
Schematic illustration of DeepRelations, an intrinsically explainable neural network architecture for predicting compound-protein interactions. Three linked relational modules (Rel-CPI in the small yellow boxes) correspond to three stages of attention focusing. Each module embeds relational features with joint attentions over pairs of protein residues and compound atoms (details on the right). In comparison, DeepAffinity+ has a single module with all relational features lumped together. Both methods are structure-free and protein structures are just for illustration.

There are three relational modules (Rel-CPI) corresponding to three stages. Their attentions are trained to progressively focus on putative binding *k*-mers, residues, and pairs; and earlier-stage attentions guide those in the next stage through regularization. In each Rel-CPI module, there are *K* = 10 types of atomic “relational” features for proteins or compounds (9 relation (sub)types are described next and the last is the union of all 9 types of features). All types of relational features are individually fed to aforementioned neural network pairs (for instance, HRNN for protein sequences and GCN for compound graphs, or HRNN-GCN in short), concatenated, and jointly embedded for proteins and compounds with attentions over residue-compound pairs. The embedding output (based on joint attentions for contact prediction) of the last module is fed to CNN and MLP layers for affinity prediction. All three modules are trained end-to-end as a single model. In contrast, DeepAffinity+ only has one module without multi-stage focusing; and its module only uses the last type of relational features (the union of the first 9 types).

#### Physics-inspired relational modules

The relational modules are inspired by physics. Specifically, atomic “relations” or interactions constitute the physical bases and explanations of compound-protein interaction affinities and are often explicitly modelled in force fields. We have considered the following six types of relations with attentions paid on and additional input data defined for.

- *Electrostatic interactions*: The ion feature of a protein residue is its net charge as in the force field CHARMM36 and that of a compound atom is its formal charge. The dipole feature of a protein residue is 1 for polar residues (S, T, C, Y, N, Q and H^54^) or 0 for others; and that of a compound atom is its Gasteiger partial charge. The electrostatics thus include all four combinations (subtypes) of residue-atom relations: ion-ion, ion-dipole, dipole-ion, and dipole-dipole.
- *Hydrogen bond*: Non-covalent interaction (A· · · H–D) between an electronegative atom as a hydrogen “acceptor” (“A”) and a hydrogen atom that is covalently bonded to an electronegative atom called a hydrogen “donor” (“D”). Therefore, if a protein residue or compound atom could provide a hydrogen acceptor/donor, its hydrogen-bond feature is −1/+1; otherwise the feature value is 0. A protein residue is allowed to be both hydrogen-bond donor and acceptor. Specifically, for protein residues, amino acids of hydrogen-bond acceptors are N, D, Q, E, H, S, T, and Y; and those of hydrogenbond donors are Y, W, T, S, K, H, Q, N, and R^55^. For compound atoms, hydrogenbond acceptor or donor is defined as in the base features factory file (atom types “SingleAtomAcceptor” and “SingleAtomDonor” in the file “BaseFeatures.fdef”) of the software RDKit v. 2018.03.4.
- *Halogen bond*: A halogen bond (A· · · X–D) is very similar to hydrogen bond except that a halogen “X” (rather than hydrogen) atom (often found in drug compounds) is involved in such interactions. As standard amino acids do not contain halogen atoms, a protein residue can only be a halogen bond acceptor (“A” in A··· X-D) and assigned a nonzero halogen-bond feature of −1, only if it is amino acid S, T, Y, D, E, H, C, M, F, W^56^, N or Q. On the compound side, only a halogen atom is assigned a nonzero feature value. Specifically, halogen-bond features of iodine, bromine, chlorine and fluorine atoms are assigned at +4, +3, +2 and +1, respectively, for decreasing halogen-bonding strengths ^56^.
- *Hydrophobic interactions*: The interactions between hydrophobic protein residues and compound atoms contribute significantly to the binding energy between them. This feature is only nonzero and set at 1 for hydrophobic residues of proteins or non-polar atoms of compounds (atoms whose absolute values of partial atomic charges are less than 0.2 units^57,58^).
- *Aromatic interactions*: Aromatic rings in histidine, tryptophan, phenylalanine, and tyrosine participate in “stacking” interactions with aromatic moieties of a compound (π-π stacking). Therefore, if a protein residue has an aromatic ring, its aromatic feature is set at 1 and otherwise at 0. Similarly, if a compound atom is part of an aromatic ring, the feature is set at 1 and otherwise at 0.
- *VdW interactions*: Van der Waals are weaker interactions compared to others. But the large amount of these interactions contribute significantly to the overall binding energy between a protein and a compound. We consider the amino-acid type and the atom element as their features and use an embedding layer to derive their continuous representations.

For each (sub)type of atomic relations, corresponding protein and compound features are fed into basis neural network models such as HRNN for protein sequences and GNN for compound graphs. The embeddings over all types are concatenated for protein residues or compound atoms and then jointly embedded with joint attentions over residue-atom pairs.

#### Physical constraints as attention regularization

The joint attention matrices 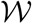 in each Rel-CPI module, for individual relations or overall, are regularized with the following two types of physical constraints. We note that, aiming at the general case where protein structures may not be available, we use sequence-predicted rather than actual structure properties (solvent exposure and residue contacts) when introducing these physical constraints.

##### Focusing regularization

In the first regularization, a constraint input is given as a matrix 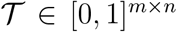 to penalize the attention matrix 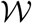 if it is focused on undesired regions of proteins. In addition, an L1 sparsity regularization is on the attention matrix 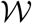 to promote interpretability as a small portion of protein residues interact with compounds. Therefore, this “focusing” penalty can be formalized as:

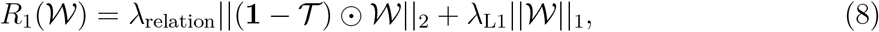

where the 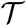 term, a parameter, can be considered as soft thresholding and the matrix norms are element-wise. The L1 regularization term in *R*_1_ (·) is only included in the first module (stage 1) where *R*_1_ (·) is the only regularization term. It is then moved to another term in the second and the last modules where multiple regularization terms are used together.

The first regularization is used for all three Rel-CPI modules or stages with increasingly focusing 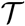. Let 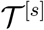 be the constraint matrix and 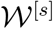 the learned attention matrix in the *s*^th^ stage. In the first stage, 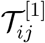, being binary, is one only for any residue *i* predicted to be solvent-exposed (relative solvent-accessible area predicted above 0.25 by SCRATCH^44,45^) in order to focus on potential surfaces. In the second stage, 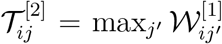 to focus on putative binding residues hierarchically learned for *k*-mers and residues at module/stage 1. In the third and last stage, 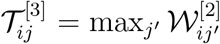 focuses on putative contacts between protein residues and compound atoms based on the learned binding residues at module/stage 2.

##### Structure-aware sparsity regularization over protein contact maps

We further develop a structure aware sparsity constraints based on known or RaptorX-predicted contact maps of the unbound protein. As sequentially distant residues might be close in 3D and form binding sites for compounds, we define overlapping groups of residues where each group consists of a residue and its spatially close neighboring residues. Just in the second stage, we introduce Group Lasso for spatial groups and the Fused Sparse Group Lasso (FSGL) for sequential groups on the overall, joint attention matrix 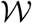:

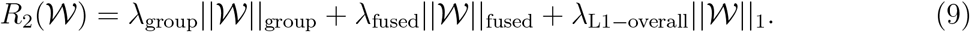

The group Lasso penalty will encourage a structured group-level sparsity so that few clusters of spatially close residues share similar attentions within individual clusters. The fused sparsity will encourage local smoothness of the attention matrix so that sequentially close residues share similar attentions with compound atoms. The L1 term again maintains the sparsity of the attention matrix 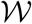. This regularization is only introduced in the second and third stages for 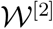 and 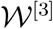, after the first-stage attention matrix 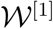 is supposedly focused on protein surfaces. The attention matrix in the last stage, 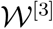, is used for predicting residue-atom contacts.

#### Supervised attentions

It has been shown in visual question answering that attention mechanisms in deep learning can differ from human attentions.^26^ As will be revealed in our results, they do not necessarily focus on actual atomic contacts (relations) in compound-protein interactions either. We have thus curated a relational subset of our compound-protein pairs with affinities, for which known ground-truth atomic contacts or relations are available. We summarize actual contacts of a pair in a matrix 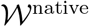 of length *m* × *n*, which is a binary pairwise interaction matrix padded with 0 to reach the maximum number of protein residues or compound atoms and then normalized by the total number of nonzero entries. We have accordingly introduced an additional third regularization term to supervise attention matrix 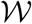 in the second and third stages:

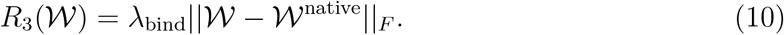

In the case of DeepAffinity+ with a single module, all three regularization terms are included as in the last module of DeepRelations.

#### Training strategy for hierarchical multi-objectives

Accuracy and interpretability are the two objectives we pursue at the same time. In our case, the two objectives are hierarchical: compound-protein affinity originates from atomic-level interactions (or “relations”) and better interpretation in the latter potentially contributes to better prediction of the former.

Challenges remain in solving the hierarchical multi-objective optimization problem. Optimizing for both objectives simultaneously (for instance, through weighted sum of them) does not respect that the two objectives do no perfectly align with each other and are of different sensitivities to model parameters. Therefore, we consider the problem as multi-label machine learning. And we design hierarchical training strategies to solve the corresponding hierarchical multi-objective optimization problem, which is detailed next.

Take DeepAffinity+ as an example. We first “pre-trained” it to minimize mean squared error (MSE) of p*K_i_*/p*K_d_* regression alone, with physical constraints turned on; in other words, attentions were regularized (through *R*_1_(·) and *R*_2_(·)) but not supervised in this stage. We tuned combinations of all hyperparameters except λ_blnd_ in the discrete set of {10^-4^, 10^-3^,10^-2^}, with 200 epochs at the learning rate of 0.001. Over the validation set, we recorded the lowest RMSE for affinity prediction and chose the hyperparameter combination with the highest AUPRC for contact prediction subjective to that the corresponding affinity RMSE (root mean square error) does not deteriorate from the lowest by more than 10%.

With the optimal values of all hyperparameters but Abi⊓a fixed, we then loaded the corresponding optimized model in the first stage and “fine-tuned” the model to minimize MSE additionally regularized by supervised attentions (through *R*_1_(·), *R*_2_(·), and *R*_3_(·)). We used the same learning rate (0.001) and training epochs (200) in fine-tuning; and we tuned λ_bind_ in the set of {10^0^,…, 10^5^} following the same strategy as in pre-training.

The tuned hyperparameters for all DeepAffinity+ variants are summarized as following. For HRNN-GCN_cstr (modeling protein sequences with HRNN and compound graphs with GCN, regularized by physical constraints in *R*_2_(·)), we chose λ_group_ = 10^-4^, λ_fused_ = 10^-3^, and λ_L1-overall_ = 10^-2^; and for its supervised version HRNN-GCN_cstr_sup, the additional λ_bind_ = 10^4^. For HRNN-GIN_cstr (modeling protein sequences with HRNN and compound graph with GIN, regularized by physical constraints in *R*_2_(·)), we chose λ_group_ = 10^-4^,λ_fused_ = 10^-3^, and λ_L1-overall_ = 10^-4^; and for its supervised version HRNN-GIN_cstr_sup, the additional λ_bind_ = 10^3^. *R*_1_(·) was for attentions on individual relations in DeepRelations and not applicable for DeepAffinity+ variants, although a surface-focusing regularization on overall attentions could be introduced.

We did similarly for hyper-parameter tuning for DeepRelations while constraining (and supervising) attentions. The whole DeepRelations model, including the three Rel-CPI modules, is trained end-to-end. ^59^ To save computational resources, we used the same hyperparameters in *R*_2_(·) (λ_L1–overall_, λ_fused_, and λ_group_) as those optimally tuned in HRNN-GCN_cstr_sup. We then tuned the rest of the hyper-parameters (λ_L1_, λ_relation_, and λ_bind_) following the aforementioned process of pre-training and fine-tuning. In the end, we chose λ_relation_ = 10^-4^, λ_L1_ = 10^-5^, λ_group_ =10^-4^, λ_fused_ = 10^-3^, λ_L1–overall_ = 10^-2^ and λ_bind_ = 10^3^ for DeepRelations. λ_bind_ is usually larger because it is multiplied to the attention-supervision term that can be orders of magnitude smaller than other terms.

## Results

We first assess the accuracy of compound-protein affinity predictions made by state-of-the-art non-interpretable methods and our interpretable DeepAffinity framework ^16^ (with new variants), using three established benchmark sets. After establishing that DeepAffinity achieves the state of the art in the accuracy of affinity prediction, we then describe a newly-curated dataset with both affinities and contacts of compound-protein interactions and assess the interpretability of various DeepAffinity versions and a competing interpretable method adapted to affinity prediction. We find that current attention-based interpretable models are not adequate for interpreting affinity (i.e., predicting contacts). Thus we proceed to regularize and supervise attentions in DeepAffinity to make DeepAffinity+ models. And we additionally use a novel, physics-inspired and intrinsically-interpretable deep relational architecture to make DeepRelations models.

Over the curated dataset, we compare our methods with a competing, structure-free interpretable method in accuracy, generalizability, and interpretability. Using a series of case studies, we also analyze the accuracy levels and spatial patterns of their top-predicted contacts, which are shown to benefit protein-ligand docking. We end the section by introducing analytics to aggregate joint attentions and decompose predicted affinity; and by demonstrating their potential utilities toward binding site prediction for proteins and SAR for compounds (scoring and lead optimization).

### DeepAffinity with interpretable attentions achieves the state-of-the-art accuracy in compound-protein affinity prediction

As the starting point of interpretability assessment and improvement, our previous interpretable DeepAffinity framework^16^ is first compared to current methods based on prediction accuracy for established benchmark sets.

For affinity benchmark datasets, we adopt three established ones of increasing difficulty, the Davis,^60^ the Kinase Inhibitor BioActivity (KIBA)^61^ and the refined set of PDBbind (v. 2019)^39^. We filtered and partitioned the first two datasets consistently with earlier studies.^15,61–63^ The Davis dataset^62^ contains all 30,056 K_d_-labeled pairs between 68 kinase inhibitors (including FDA-approved drugs) and 442 kinases, randomly split into 25,046 for training and 5,010 for testing (the widely-used “S1” setting^62^). The filtered KIBA dataset^61,62^ contains 118,254 pairs between 2,111 kinase inhibitors and 229 kinases, including 98,545 for training and 19,709 for testing (S1 split again). Other split settings were not pursued because published performances in such settings are not always available and comparable. The KIBA scores combine *k_i_, k_d_*, and IC_50_ sources for consistency and are further processed.^15,62^ As to the refined PDBbind dataset (v. 2019), we filtered and processed it (see details in the supplemental Sec. S1.1) to reach 3,505 pairs with k_i_ or k_d_ labeled between 1,149 proteins and 2,870 compounds. Compared to Davis and KIBA, the PDBbind dataset contains more diverse protein classes: 2,157 interactions with enzymes including 72 with kinases, 62 with nuclear receptors, 33 with G protein-coupled receptors (GPCRs), and 106 with ion channels. The portion of labeled compound-protein pairs is much lower than that of Davis and KIBA. We randomly split the PDBbind dataset into 2,921 pairs for training and 584 for testing.

For our framework of DeepAffinity, ^16^ we adopt various data representations and corresponding state-of-the-art neural network architectures as detailed in Methods. To model proteins, we have adopted RNN using protein SPS^16^ as input data as well as CNN and newly developed HRNN using protein amino-acid sequences. To model compounds, we have adopted RNN using SMILES as input data as well as GCN and GIN using compound graphs with node features and edge adjacency. ^19^ In the end, we have tested five DeepAffinity variants (including four new) for protein-compound pairs, including RNN-RNN,^16^ RNN-GCN, CNN-GCN, HRNN-GCN, and HRNN-GIN. Names before and after hyphens indicate models to embed proteins and compounds, respectively; and embeddings of a pair of protein and compound are passed through joint attentions in Eq. (6) before being fed to a convolutional neural network (CNN) and multi-layer perceptrons (MLP).^16^ For instance, the first one, RNN-RNN indicates that protein SPS sequences are modeled by RNN and compound SMILES or graphs are modeled by RNN. This is essentially our previous method^16^ except that no unsupervised pretraining or ensemble averaging is used here. We have tuned hyperparameters for DeepAffinity variants including learning rate ({10^-3^,10^-4^}), batch size ({64, 128} (16 for CNN-GCN due to the limit of GPU memory) and dropout rate ({0.1, 0.2}) using random 10% of training data as validation sets. When HRNN was used to model protein sequences, we have also tuned k-mer lengths and group sizes in pairs ({(40,30), (48,25), (30,40), (25,48), (15,80), (80,15)} for Davis and {(40,25), (50,20), (25,40), (20,50)} for KIBA and PDBbind) using the validation sets.

For comparison, we use published current methods that are not structure-based, including DeepDTA,^15^ KronRLS,^64^ and WideDTA,^65^ all of which are non-interpretable. Their results for the Davis and KIBA sets were self reported in individual studies and summarized in a comparison study. ^63^ And their results for the PDBbind set are derived by re-training released source codes with published hyper-parameter grids and individual training sets (except wideDTA whose codes are not available). In addition, we compare to structure-free methods that are interpretable. Except DeepAffinity, the only other interpretable method published so far (Gao et al.) was for predicting binary compound-protein interaction.^18^ As its codes are not publicly available, we have implemented the method, revised its model’s last layer (sigmoid) and retrained the model for affinity prediction using each training set. To ensure fair comparison, all deep-learning models including our DeepAffinity variants here are trained for 100 epochs or until convergence (the validation loss does not improve within 15 epochs), as competing methods previously did. ^63^

We compare aforementioned competing methods and DeepAffinity variants in accuracy using two assessment metrics: RMSE (root mean squared error; see Table 1) and CI (concordance index; see Table 2). Whereas RMSE evaluates the proximity between predictions are to corresponding native values, CI, ^66^ often used for virtual screening, measures the probability of correctly ordering non-equal pairs. We summarize the results in Tables 1 and 2.

**Table 1:** Comparing current methods (non-interpretable except Gao et al.) and interpretable DeepAffinity variants in prediction accuracy (measured by RMSE, the lower the better) for the Davis, KIBA and PDBbind benchmark sets. The best performance in each dataset is bold-faced.

| RMSE | DeepDTA | KronRLS | WideDTA | Gao et al.<sup>b</sup> | DeepAffinity |  |  |  |  |
| --- | --- | --- | --- | --- | --- | --- | --- | --- | --- |
|  |  |  |  |  | RNN-RNN<sup>16</sup> | RNN-GCN | CNN-GCN | HRNN-GCN | HRNN-GIN |
| Davis | 0.5109<sup>a</sup> | 0.6080<sup>a</sup> | 0.5119<sup>a</sup> | 0.7864 | 0.5032 | 0.5095 | 0.8106 | **0.5019** | 0.6604 |
| KIBA | 0.4405<sup>a</sup> | 0.6200<sup>a</sup> | **0.4230<sup>a</sup>** | 0.7368 | 0.4335 | 0.5367 | 0.8244 | 0.4480 | 0.6669 |
| PDBbind | 2.0631 | 1.8005 | - | 1.8071 | 1.4524 | **1.4277** | 1.5580 | 1.4743 | 1.4858 |

<sup>a</sup> Self-reported and published results as summarized in Thafar et al.<sup>63</sup>
<sup>b</sup> Originally a binary classifier, it was implemented and revised by us for affinity prediction.

**Table 2:** Comparing current methods (non-interpretable except Gao et al.) and interpretable DeepAffinity variants in prediction accuracy (measured by concordance index or CI, the larger the better) for the Davis, KIBA and PDBbind benchmark sets. The best performance in each dataset is bold-faced.

| CI | DeepDTA | KronRLS | WideDTA | Gao et al.<sup>b</sup> | DeepAffinity |  |  |  |  |
| --- | --- | --- | --- | --- | --- | --- | --- | --- | --- |
|  |  |  |  |  | RNN-RNN<sup>16</sup> | RNN-GCN | CNN-GCN | HRNN-GCN | HRNN-GIN |
| Davis | 0.8780<sup>a</sup> | 0.8830<sup>a</sup> | 0.8860<sup>a</sup> | 0.7824 | **0.9000** | 0.8808 | 0.7373 | 0.8814 | 0.8224 |
| KIBA | 0.8630<sup>a</sup> | 0.7820<sup>a</sup> | **0.8750<sup>a</sup>** | 0.7335 | 0.8423 | 0.7968 | 0.5761 | 0.8420 | 0.6893 |
| PDBbind | 0.7125 | 0.7197 | - | 0.7610 | **0.8042** | 0.7543 | 0.7119 | 0.7544 | 0.7398 |

<sup>a</sup> Self-reported and published results as summarized in Thafar et al.<sup>63</sup>
<sup>b</sup> Originally a binary classifier, it was implemented and revised by us for affinity prediction.

From both tables we conclude that the original DeepAffinity method^16^ (RNN-RNN; RNN for protein SPS and RNN for compound SMILES) and its variants compared favorably to the state of the art. Specifically, the DeepAffinity variants achieved the best performances in RMSE and CI for both the Davis dataset and the most diverse and sparse dataset of PDBbind. And it closely followed the best performances (WideDTA) for the KIBA dataset. In particular, the newly introduced HRNN models for protein sequences (higher-resolution than SPS) and graph models GCN & GIN for compound graphs achieved the best or close-to-the-best performances, which enables interpreting affinity prediction at the level of protein residues and compound atoms without sacrificing the accuracy. Considering that other methods are not interpretable and the only exception Gao et al. did not perform as well, the performances of interpretable DeepAffinity variants are particularly impressive.

### Our new dataset for both affinity and contact prediction is diverse and challenging

To support systematic assessment and development of explainable affinity prediction, we have constructed a dataset of 4,446 compound-protein pairs (between 1,287 proteins and 3,672 compounds) with both affinity values (p*K_i_* or p*K_d_*) and atomic contacts (available in co-crystal structures). More details are included in Methods and the supplemental Sec. S1.1.

The dataset contains diverse proteins and compounds. Among the 4,446 pairs, there are 2,913 interactions with enzymes including 114 with kinases, 105 with nuclear receptors, 89 with GPCRs, and 111 with ion channels. The enzymes are across all seven enzyme commission classes (see details including EC class breakdowns in the supplemental Sec. 1.1). The 3,672 compounds cover wide ranges of physicochemical properties (logP, molecular weight, and affinity values) as seen in Figure 3.

**Figure 3:**
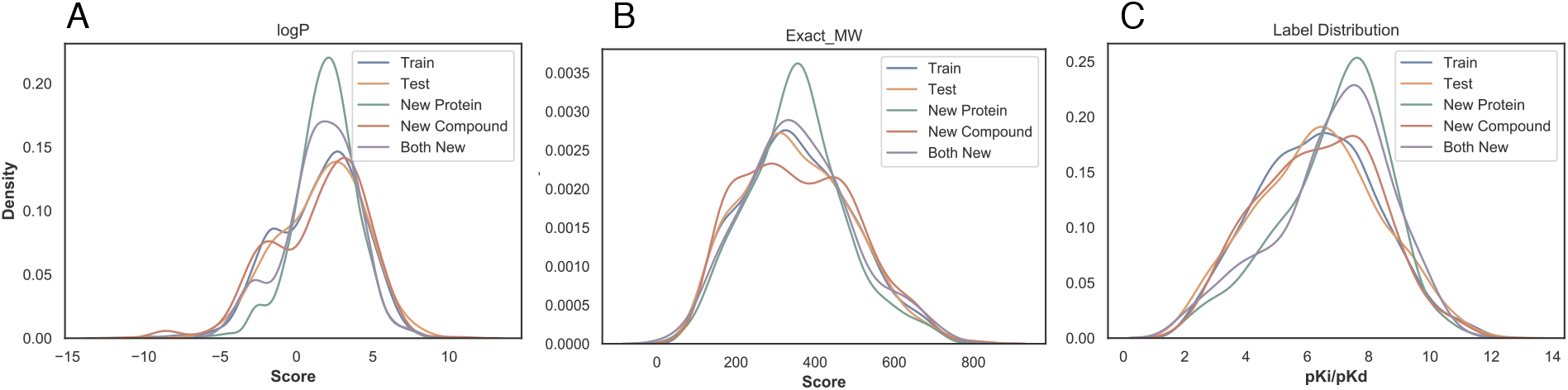
The distributions of compound properties across various subsets: A. logP; B. exact molecule weight; and C. p*K_i_*/p*K_d_* labels.

The dataset is split into training including validation (2,334), test (591), new-protein (795), new-compound (521), and both-new sets (205), as illustrated in Figure 1. Compared to the test set, the three generalization sets not only contain new proteins or/and compounds but also mainly consist of very dissimilar proteins or/and compounds compared to the training set, which suggest their challenges for machine learning. For instance, the new-protein set only contains proteins not present in the training set. 454 (57.1%) pairs in the set involve new proteins whose global sequence identities to the closest training proteins are below 30%. And 452 (56.8%) pairs involve new proteins whose local binding *k*-mer identities are below 30% (note that only around 10% residues of an average binding *k*-mer are binding residues). Similarly, 414 (79.5%) new-compound pairs involve new compounds whose Tanimoto scores (details in the supplemental Sec. 1.4) to the closest training compounds are below 0.5. The both-new set only contains pairs of new proteins and new compounds with similarly low resemblance to the training set. 98 (47.8%) pairs involve new proteins with sequence identity below 30% *and* new compounds with Tanimoto scores below 0.5. So the both-new set is expected to be the most challenging set among the four for the generalizability of machine learning models. Pair breakdowns are visualized in part of Figure 6 (counts). In addition, Jensen-Shannon distances between compound properties of training and those of the other sets are given in Table S1, similarly revealing the most challenging both-new set.

**Figure 4:**
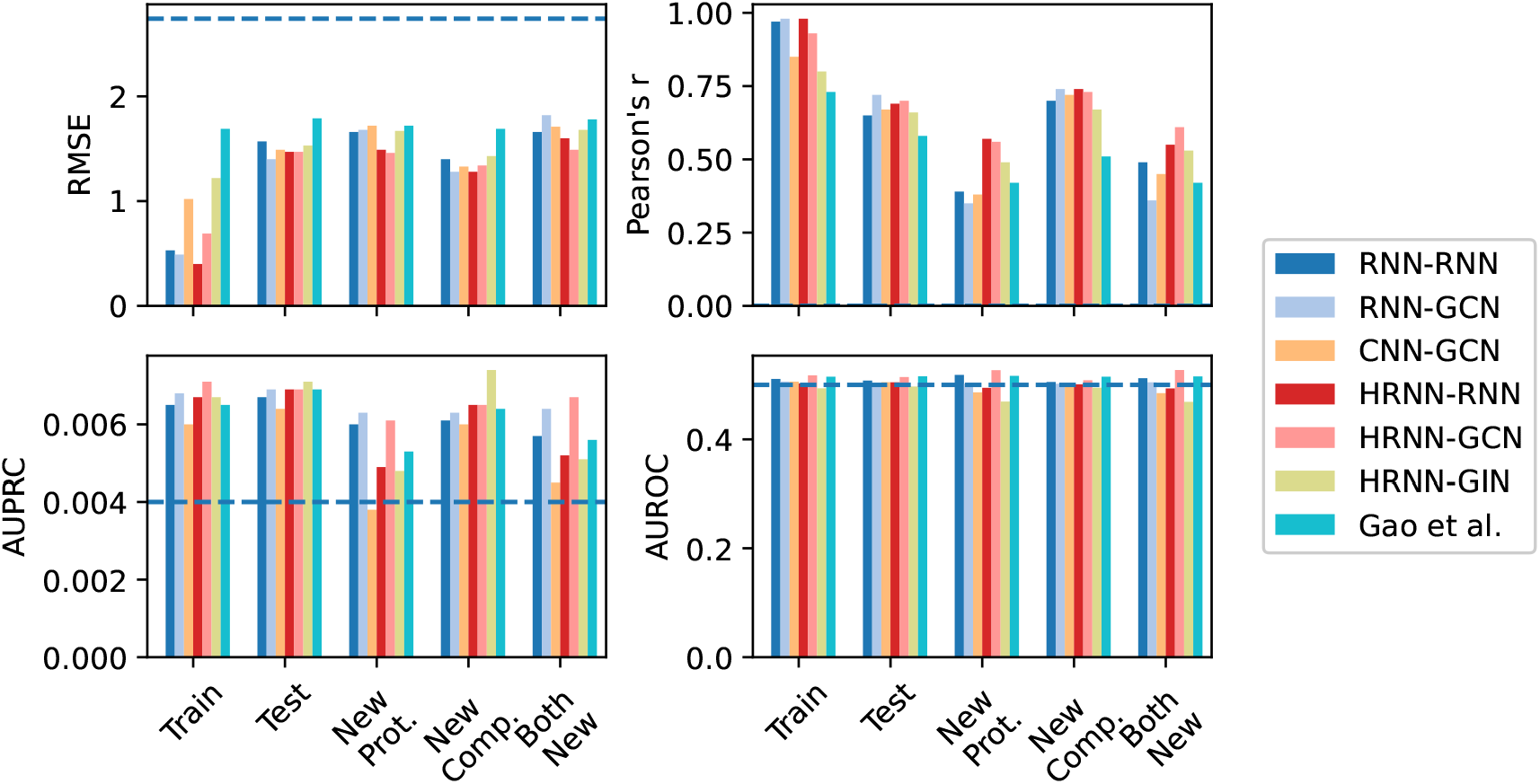
Comparing accuracy and interpretability among various versions of DeepAffinity with unsupervised joint attention mechanisms as well as another interpretable method (Gao et al.). Separated by hyphens in legends are neural network models for proteins and compounds respectively. A horizontal dashed line indicates the performance of a random predictor.

**Figure 5:**
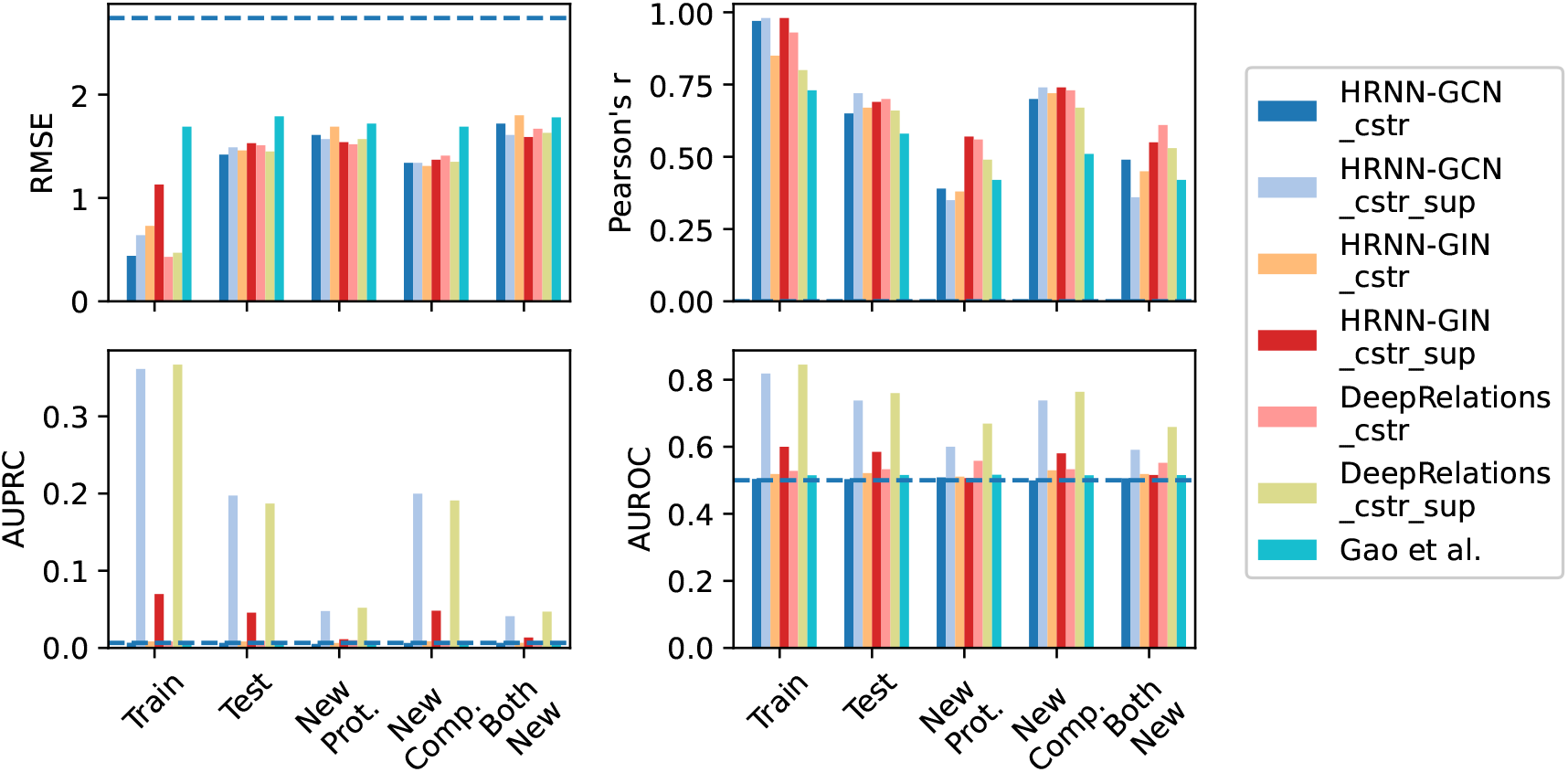
Comparing accuracy and interpretability among various versions of DeepAffinity+ (DeepAffinity with regularized and supervised attentions) and DeepRelations. “cstr” in legends indicates physical constraints imposed on attentions through regularization term *R*_2_(·), whereas “sup” indicates supervised attentions through regularization term *R*_3_(·). A horizontal dashed line indicates the performance of a random predictor.

**Figure 6:**
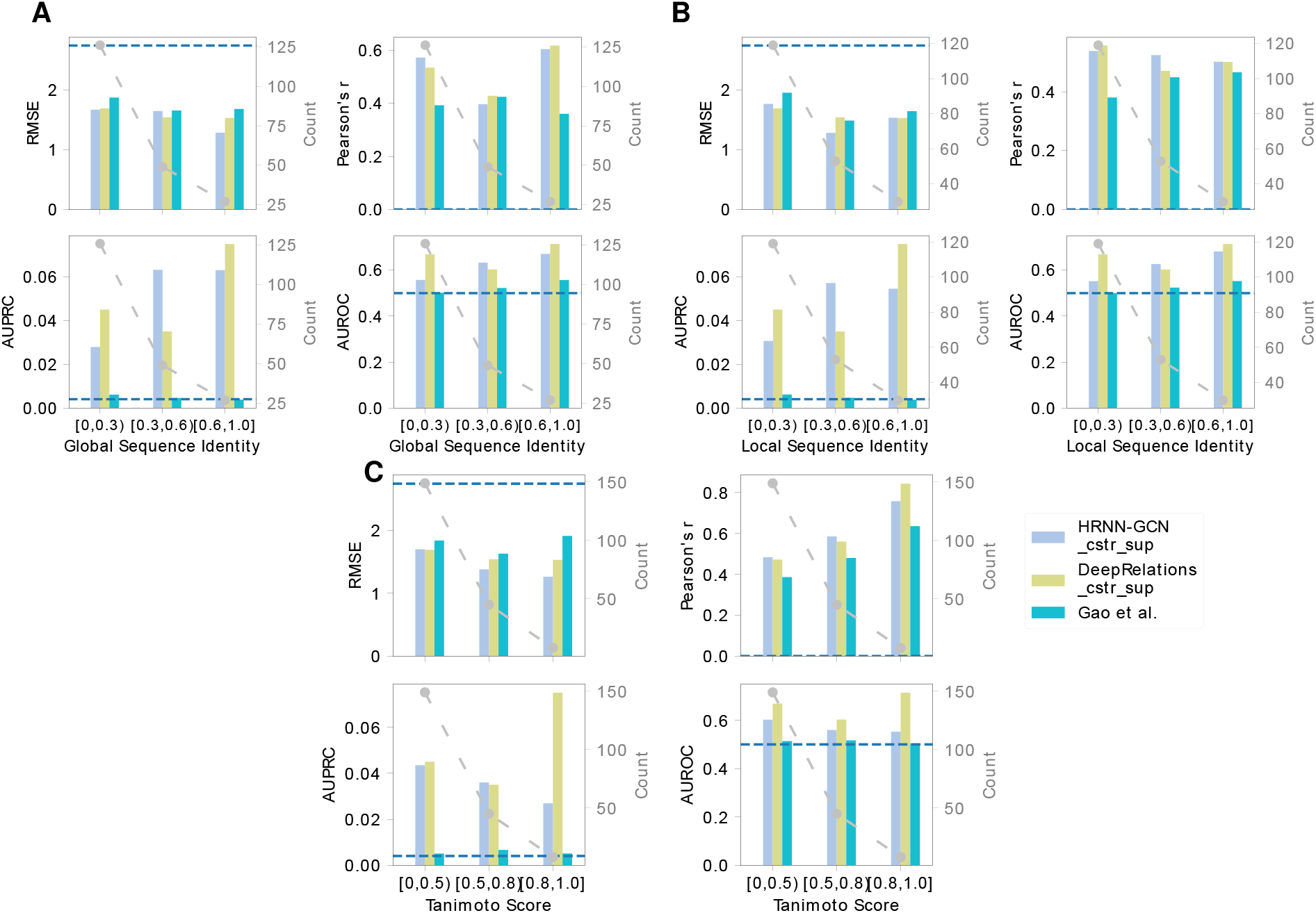
Comparing DeepAffinity+, DeepRelations, and Gao’s method in the generalizability of affinity prediction (RMSE and Pearson’s r) and contact prediction (AUPRC and AUROC) to molecules unlike training data. A horizontal dashed line indicates the performance of a random predictor.

### Attentions alone are inadequate for interpreting compound-protein affinity prediction

Now that we have established the accuracy of attention-embedded DeepAffinity and constructed a suitable dataset, our first task for interpretability is to systematically assess the adequacy of attention mechanisms for interpreting model-predicted compound-protein affinities. To that end, using our newly curated benchmark set for both affinity and contact prediction, we have tested six DeepAffinity variants for protein-compound pairs (including RNN-RNN, RNN-GCN, CNN-GCN, HRNN-RNN, HRNN-GCN, and HRNN-GIN) as well as the only other interpretable method (Gao et al.) that is also attention-based and adapted by us from a classifier to a regressor. All models are re-trained using the new training set with details in Methods. The first two DeepAffinity (RNN-RNN and RNN-GCN) models’ attentions on proteins are at the secondary structure levels. Their joint attentions were thus converted to residue-atom matrices, using equal weights across all residues within a secondary structure, in the post-analysis of interpretability. The rest have joint attentions at the level of pairs of protein residues and compound atoms.

The accuracy of affinity prediction, measured by RMSE and Pearson’s *r* in p*K_i_*/p*K_d_*, is summarized for the DeepAffinity variants in the top panel of Figure 4 and Table S3. Overall, all variants have shown affinity RMSE (Pearson’s r) around 1.5 (0.65), 1.6 (0.50), 1.4 (0.70), and 1.7 (0.50) for the default test, new-protein, new-compound, and both-new sets, respectively. In particular, the HRNN-GCN version achieved an RMSE (Pearson’s r) of 1.47 (0.70), 1.46 (0.56), 1.34 (0.73), and 1.49 (0.61) for the four sets, respectively, showing a robust accuracy profile. In contrast, the competing method (Gao et al.) has worse RMSE values between 1.72 and 1.87 and worse Pearson’s r between 0.42 and 0.58.

The interpretability of affinity prediction is assessed against ground truth of intermolecular residue-atom contacts, as shown in the bottom panel of Figure 4 and Table S3. Specifically, we use joint attention scores to classify all possible residue-atom pairs into contacts or non-contacts. As contacts only represent a tiny portion 0.0040±0.0029 in our dataset) of all possible pairs, we use the the area under the precision-recall curve (AUPRC) as the major metric and the area under the receiver operating characteristic curve (AUROC) as a reference, to assess such binary classification. Here AUPRC/AUROC is averaged over all pairs involved in the corresponding set. Interestingly, compared to chance (AUPRC=0.004 and AUROC=0.5), all attention-based models including DeepAffinity variants and Gao et al. only had slightly better AUPRC (around 0.006 albeit a 50% improvement) except CNN-GCN for the new-protein set. The best DeepAffinity variant, HRNN-GCN, did improve against Gao et al..

From the results above, we conclude that attention mechanisms alone are inadequate for the interpretability of compound-protein affinity predictors, regardless of the choice of commonly used, generic neural network architectures.

### Regularizing attentions with physical constraints modestly improves interpretability

Our next task is to enhance the interpretability of compound-protein affinity prediction beyond the level achieved by attention mechanisms alone. The first idea is to incorporate domain-specific physical constraints into model training. The rationale is that, by bringing in the (predicted) structural contexts of proteins and protein-compound interactions, attentions can be guided in their sparsity patterns accordingly for better interpretability.

We start with the two best-performing DeepAffinity variants so far (HRNN-GCN and HRNN-GIN) where protein amino-acid sequences are modeled by hierarchical RNN and compound graphs by various GNNs (including GCN and GIN). And we introduce structure-aware sparsity regularization *R*_2_(·) to the two models to make “DeepAffinity+” variants. The resulting HRNN-GCN_cstr and HRNN-GIN_cstr models with physical constraints are assessed in Figure 5 and Table S4. Compared to the the non-regularized counterparts in Figure 4 and Table S3, both models achieved similar accuracy levels across various test sets for affinity prediction. As to their interpretability, HRNN-GCN_cstr had similar AUPRC as before regularization (0.006) and HRNN-GIN_cstr slightly improved AUPRC to around 0.008, although both were still close to the baseline (0.004). These results suggest that incorporating physical constraints to structurally regularize the sparsity of attentions is useful for improving interpretability but may not be enough.

### Supervising attentions significantly improves interpretability

As regularizing attentions with physical constraints was not enough to enhance interpretability, our next idea is to additionally supervise attentions with ground-truth contact data available to some but not all training examples. Again we introduce “DeepAffinity+” models starting with HRNN-GCN and HRNN-GIN, by both regularizing and supervising attentions (using *R*_2_(·) and *R*_3_(·)).

The performances of resulting HRNN-GCN_cstr_sup and HRNN-GIN_cstr_sup models are shown in Figure 5. Importantly, HRNN-GCN_cstr_sup (light blue) significantly improved interpretability of affinity prediction without the sacrifice of accuracy. The average AUPRC improved to 0.197, 0.048, 0.200, and 0.041 for the default test, new-protein, new-compound, and both-new sets, representing a 30.4, 9.2, 31.2, and 6.3-fold increase, respectively, compared to the version with just regularization but not supervision of attentions (HRNN-GCN_cstr). The performances also represented a 32.9, 9.9, 35.1, and 8.6-fold increase, respectively, compared to Gao et al.. Interestingly, supervising attentions in HRNN-GIN did not see as significant improvement in interpretability.

### Building explainability into DeepRelations architecture further improves interpretability

Toward better interpretability, besides regularizing and supervising attentions, we have further developed an explainable, deep relational neural network named DeepRelations. Here atomic “relations” constituting physical bases and explanations of compound-protein affinities are explicitly modeled in the architecture with multi-stage gradual “zoom-in” to focus attentions. In other words, the model architecture itself is intrinsically explainable by design.

The performances of the resulting DeepRelations (with both regularized and supervised attentions) are shown in Figure 5 (yellow-green “DeepRelations_cstr_sup”). With equally competitive accuracy in affinity prediction as all previous models, DeepRelations achieved further improvements in interpretability. The AUPRC values were similar to the best Deep-Affinity+ model (HRNN-GCN_cstr_sup): 0.187, 0.052, 0.191, and 0.047 for the default test, new-protein, new-compound, and both-new sets, respectively. The AUROC values improved to 0.76, 0.67, 0.76, and 0.66 for the four sets, representing an increase of 0.03, 0.07, 0.03, and 0.07 compared to those of the best DeepAffinity+, respectively.

To disentangle various components of DeepRelations and understand their relative contributions to DeepRelations’ improved interpretability, we removed components from DeepRelations for ablation study. Besides regularized and supervised attentions, we believe that the main contributions in the architecture itself are (1) the multi-stage “zoom-in” mechanisms that progressively focus attentions from surface, binding *k*-mers, binding residues to binding residue-atom pairs; and (2) the explicit modeling of atomic relations that can explain the structure feature-affinity mappings consistently with physics principles.

We thus made three DeepRelations-variants: DeepRelations without multi-stage focusing, without explicit atomic relations, or without both. And we compare them with DeepRelations in Figure S1. Consistent with our conjecture, we found that, the explicit modeling of atomic relations was the main contributor as its removal led to worse affinity and contact predictions in new-protein and both-new sets. The multi-stage focusing also contributes as its removal led to worse affinity prediction for both new-compound and both-new sets.

### Validation of affinity prediction

To validate the affinity accuracy of our two final models HRNN-GCN_cstr_sup (DeepAffinity+ hereinafter) and DeepRelations_cstr_sup (DeepRelations hereinafter), we have performed several randomization tests. First, using random sampling of the training set would lead to affinity RMSEs above 2.7 and Pearson’s *r* around 0; whereas using the sample mean would lead to affinity RMSEs between 1.85 and 2.02 and an undefined Pearson’s *r*. Both random affinity predictors performed considerably worse than DeepAffinity+ and DeepRelations (RMSE between 1.3 and 1.6 and Pearson’s *r* between 0.5 and 0.7). Second, *Y*-randomization tests^67^ of DeepAffinity+ and DeepRelations (20 trials each) led to much worse affinity prediction (RMSE between 2.20 and 2.45 and Pearson’s r around 0). Compound-randomization tests of our two models had similar results (RMSE between 1.95 and 2.22 and Pearson’s *r* around 0 for new proteins). More details can be found in Tables S5-7. Therefore, we conclude that our models’ affinity accuracy is significantly better than chance correlations.

To further improve the accuracy of affinity prediction, we have constructed ensembles of DeepAffinity+, DeepRelations, and both, by using combinations of hyper-parameters (such as the dropout ratio, λ_bind_, and the width of fully-connected layers). More details can be found in the supplemental Sec. S2.5. Notably, the DeepAffinity+ ensemble decreased affinity RMSE from 1.49 to 1.29, increased Pearson’s r from 0.68 to 0.77, and increased predictive *R*^2^ from 0.45 to 0.59 for the test set. It similarly improved the accuracy of affinity prediction, albeit to a lesser extent, for other sets involving new molecules. More results are reported in Table S8.

### Better interpretability helps better accuracy and generalizability of affinity prediction

To examine whether the more interpretable affinity predictors are also more accurate in affinity prediction, we compare our two final models HRNN-GCN_cstr_sup (DeepAffinity+ hereinafter) and DeepRelations_cstr_sup (DeepRelations hereinafter) to the competing interpretable affinity predictor Gao et al. Re-examining earlier results (Figure 5 and Table S4) shows that DeepAffinity+ and DeepRelations with much better interpretability (AUPRC increase between 8.6 and 59-fold) than Gao et al. are also more accurate in affinity prediction (RMSE drop between 0.15 and 0.42 and Pearson’s r increase around 0.25) over all sets considered. Even when we compare DeepAffiity+ and DeepRelations to their attention-unsupervised counterparts (HRNN-GCN_cstr and DeepRelations_cstr), we find that better interpretability (contact prediction) leading to better accuracy (lower RMSE and higher Pearson’s *r* for affinity prediction) in 6 of 8 cases where the only exceptions occurred when AUPRC values were low.

Here we further compare DeepAffinity+ and DeepRelations to Gao et al. in affinity and contact prediction over multiple difficulty ranges (measured by protein global sequence identity, protein local binding *k*-mer sequence identity, or compound Tanimoto scores) of the new-compound, new-protein, and both-new sets. The results are reported in Figure 6 as well as Figures S2–7 and Table S9. We find that the same conclusion (better interpretability leads to better accuracy) also applies where model generalizability is needed the most: pairs involving very dissimilar proteins (global or local sequence identity below 30%) or/and compounds (Tanimoto scores below 0.5) compared to training molecules. Importantly, in those cases demanding generalizability the most, DeepAffinity+ and DeepRelations have much better accuracy (affinity-prediction RMSE decrease between 0.14 and 0.40 and Pearson’s r increase between 0.10 and 0.18) as well as significantly improved interpretability (contact-prediction AUPRC increase between 5.9 and 33.3-fold) compared to Gao et al..

DeepAffinity+ and DeepRelations also showed competitive generalizability in both affinity and contact prediction. From the most similar proteins (sequence identity above 60%) to the least (sequence identity below 30%), affinity-prediction RMSE values of DeepAffinity+ (DeepRelations) only increased 0.13 (0.08) for the new-compound set and increased 0.00 (0.16) for the most challenging both-new set. From the most similar compounds (Tanimoto scores above 0.8) to the least (Tanimoto scores below 0.5), affinity-prediction RMSE values of DeepAffinity+ (DeepRelations) only increased 0.14 (0.08) for the new-compound set and increased 0.43 (0.48) for the most challenging both-new set. Similar conclusions can be made about their generalizability in contact prediction.

### Case studies

Now that we have established and explained how DeepAffinity+ and DeepRelations significantly improve the interpretability of compound-protein affinity prediction, we went on to delve into their affinity and contact predictions in comparison to Gao et al. using a series of cases studies of increasing difficulty. Summary performances of the five cases are reported in Table 3. DeepAffinity+ and DeepRelations had better affinity and contact prediction in all cases compared to the competing method whose top-10 predicted contacts failed to produce any native contacts. In order to understand model behaviors, our analysis next would focus on the patterns of top-10 contacts predicted by DeepAffinity+ and DeepRelations compared to Gao et al..

**Table 3:** Performance summary of three interpretable methods for five case studies.

|  |  | DeepAffinity+ |  |  |  | DeepRelations |  |  |  | Gao et al. |  |  |  |
| --- | --- | --- | --- | --- | --- | --- | --- | --- | --- | --- | --- | --- | --- |
| Protein | Ligand | Affinity Error | Contact AUROC | Contact AUPRC | Top-10 Contact Precision | Affinity Error | Contact AUROC | Contact AUPRC | Top-10 Contact Precision | Affinity Error | Contact AUROC | Contact AUPRC | Top-10 Contact Precision |
| Two compounds bind to the same pocket of a new, non-homologous protein (different affinity-prediction quality) |  |  |  |  |  |  |  |  |  |  |  |  |  |
| CA2 | AL1 | 1.89 | 0.658 | 0.284 | 0.5 | 2.70 | 0.828 | 0.075 | 0.6 | 3.28 | 0.500 | 0.006 | 0.0 |
|  | IT2 | 2.92 | 0.601 | 0.034 | 0.3 | 3.03 | 0.780 | 0.309 | 0.5 | 3.09 | 0.630 | 0.009 | 0.0 |
| Two new compounds bind to distinct pockets of a protein |  |  |  |  |  |  |  |  |  |  |  |  |  |
| PYGM | CPB | 0.10 | 0.552 | 0.006 | 0.1 | 0.39 | 0.513 | 0.005 | 0.0 | 0.61 | 0.522 | 0.001 | 0.0 |
|  | T68 | 0.68 | 0.944 | 0.675 | 1.0 | 0.66 | 0.908 | 0.610 | 1.0 | 1.80 | 0.635 | 0.006 | 0.0 |
| A new compound very dissimilar to training compounds binds to a new protein non-homologous to training proteins |  |  |  |  |  |  |  |  |  |  |  |  |  |
| LCK | LHL | 2.12 | 0.500 | 0.053 | 0.4 | 1.30 | 0.702 | 0.053 | 0.4 | 2.89 | 0.540 | 0.005 | 0.0 |

#### Two compounds bind to the same pocket of a new protein non-homologous to training examples

Our first case study involves a protein from the new-protein set, human carbonic anhydrase II (CA2, UniProt ID: P00918), that has no close homolog in the training set. Specifically, the closest training protein would be human carbonic anhydrase IV (CA4, UniProt ID: P22748) with a sequence identity below the 30% threshold (29%). We choose two compounds (HET IDs: AL1 and IT2) that bind to the same pocket of CA2 with distinct sizes (AL1 is larger by 14 heavy atoms) and affinity-prediction quality (see Table 3).

We compare in Figure 7 the top-10 contacts between protein residues and compound atoms that are predicted by three methods. Top-predicted contacts by Gao et al. were scattered across protein residues that are far from the binding site, failing to match any native contact. In contrast, those top-10 contacts predicted by DeepAffinity+ and DeepRelations were more focused in or near the binding site, containing 3 to 6 native contacts that are direct, first-shell contacts. Between our two models, DeepRelations showed better contact prediction in these two cases: its top-10 predictions were more focused in the binding site and contained 60% and 50% native contacts for compounds AL1 and IT2, respectively. The more focused contact prediction of our methods could be attributed to structure-aware regularization using protein residue-residue contact maps. DeepRelations had better focus than DeepAffinity+, possibly due to the multi-stage focusing strategy.

**Figure 7:**
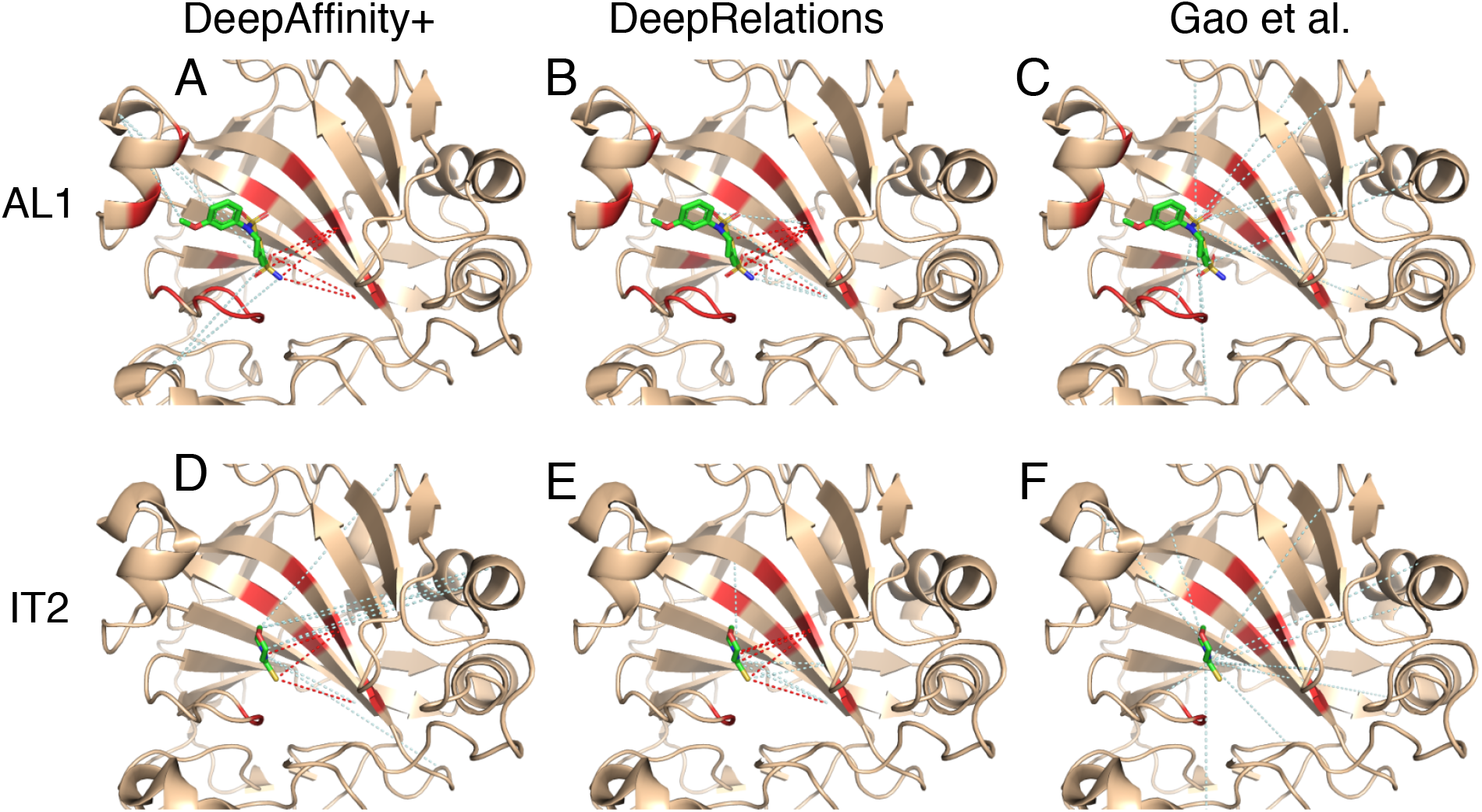
Structural visualization of top-10 intermolecular contacts predicted by DeepAffinity+ (left), DeepRelations (middle) and Gao et al. (right) for two test cases. Here two compounds (AL1: top panels A–C and IT2: bottom panels D–F; stick representations) bind to the same pocket of the human carbonic anhydrase II that is new and non-homologous to training data (wheat cartoons where binding residues are highlighted in red). Shown in dashed lines are top-10 predicted contacts (interactions between protein residues and compound atoms). The dashed lines in red and pale cyan highlight correct and incorrect predictions, respectively, according to native, direct contacts retrieved by LigPlot.

Even the incorrect predictions of DeepRelations can correspond to residue-atom pairs that are close (but above the 4 Å-cutoff used in the first-shell contact definition). For instance, in the case of compound AL1, the 4 incorrect predictions all corresponded to correct binding-residues that were paired to wrong compound atoms. In the case of compound IT2, the 5 incorrect predictions included 2 that paired correct binding-site residues to wrong atoms and 3 that included (the very next) sequence neighbors of correct binding-site residues.

These two cases also provided examples to interpret the values of AURPC and top-10 contact precision. A seemingly “low” AUPRC value of 0.075 can lead to 5 of 10 top predictions being correct. The reason is that native contacts represent a rare minority (0.004) among all possible residue-atom pairs and an AUPRC value of 0.075 actually represents over 18-fold increase compared to the baseline AUPRC by chance. Meanwhile, a top-10 contact precision of 0.4 predicted by our structure-free methods is close to the average level (0.44) achieved by a popular structure-based protein-ligand docking program, AutoDock Vina, ^68^ under default settings.^69^

#### Two new compounds bind to distinct pockets of a protein

Our next case study involves two compound-protein pairs from the new-compound set, where two compounds (HET ID: CPB and T68) not present in the training set bind to two distinct pockets of the rabbit glycogen phosphorylase (PYGM, UniProt ID: P00489). The protein is present in the training set with 38 ligands (all but one are occupying the same pocket as T68). In addition, the compound CPB does not resemble its closest training example interacting with the same protein (HET ID: 62N) when 62N rather occupies the same pocket as T68. Therefore, contact prediction for the CPB case would be much more challenging. Indeed, our results supported the conjecture (Tabe 3). In their top-10 contact predictions our both models achieved 100% native contacts for T68 but just 10% (DeepAffinity+) or even 0% (DeepRelations) for CPB. They had good estimation of binding affinity for both cases.

A closer look into their contact predictions reveal more insights. As seen in Figure 8, consistent with our earlier observations, Gao et al.’s contact predictions are dispersed across the whole protein whereas ours are focused. In the case of T68, our predictions are focused in the correct binding site (and even the correct binding residues). However, in the case of CPB, our predictions are actually still focused in the same site as they did for T68, only being wrong this time. Interestingly many falsely-predicted contacts for CPB were not only in the other binding site (circled area) but also with the T68 binding residues. This model behavior is understandable when almost all training examples, including a very similar compound, are indicating a different site. It also reveals a situation that would challenge more generalizability and demand more explainability from machine learning methods. Intriguingly, DeepAffinity+ still managed to make one correct contact prediction (pointed at by a red arrow).

**Figure 8:**
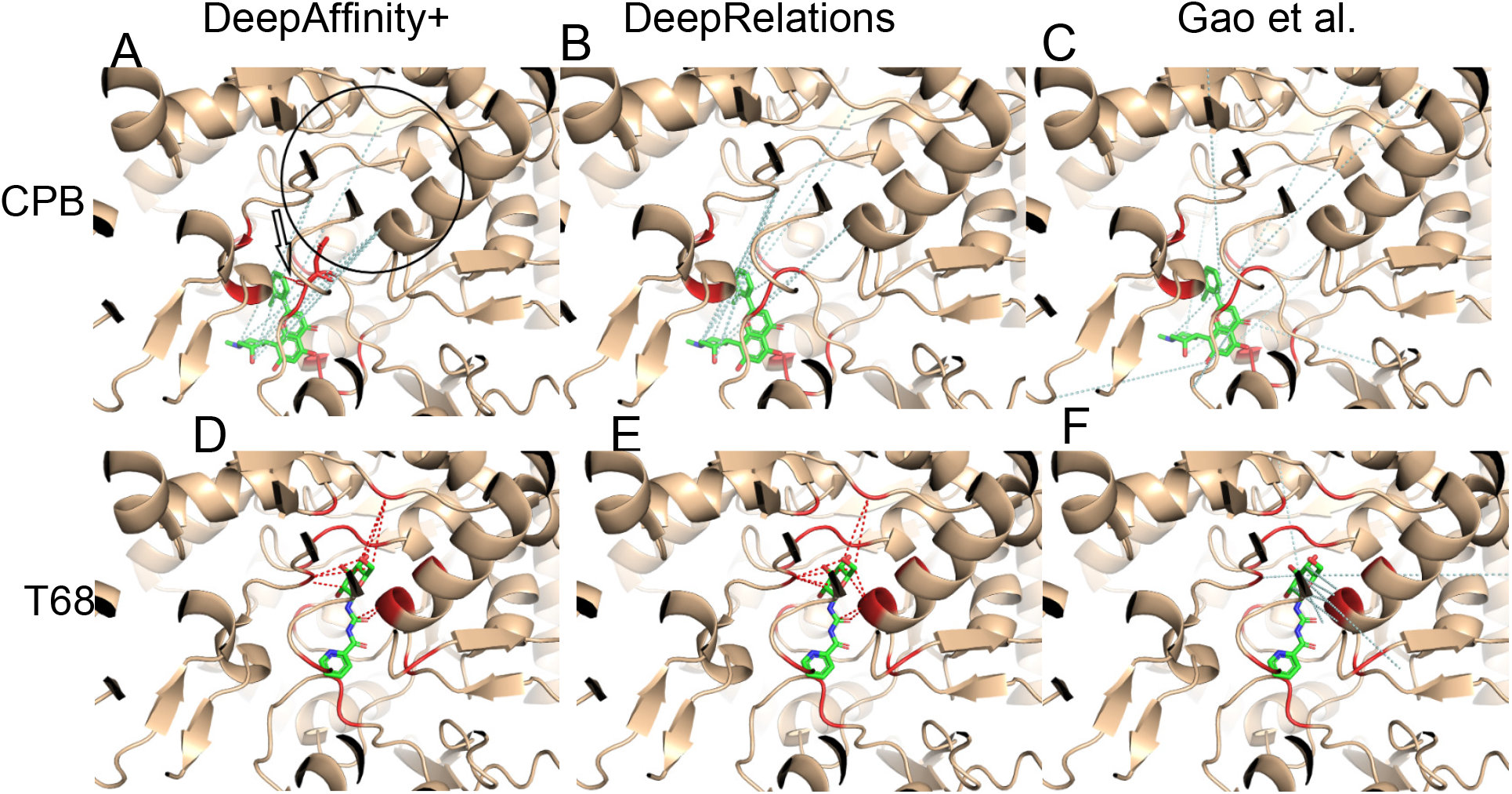
Structural visualization of top-10 intermolecular contacts predicted by DeepAffinity+ (left), DeepRelations (middle) and Gao et al. (right) for another two test cases. Here two compounds that are new to training data (CPB: top panels A-C and T68: bottom panels D-F; stick representations) bind to distinct pockets of the human glycogen phosphorylase (wheat cartoons where binding residues are highlighted in red). Shown in dashed lines are top-10 predicted contacts (interactions between protein residues and compound atoms), including correct (red) and incorrect (pale cyan) ones according to LigPlot’s definition of native, direct contacts. The black hollow arrow in panel A points to the only correct prediction by DeepAffinity+ and the black circle there indicates the binding site for T68. Interestingly, many incorrect predictions by DeepAffinity+ and DeepRelations for CPB were with binding residues to T68.

#### A pair of new protein and new compound very dissimilar to training examples

Our last case study is even more challenging in that both the protein (the human tyrosine-protein kinase Lck, LCK in short, UniProt ID: P06239) and the compound (HET ID: LHL) are new and they don’t even resemble training examples. Specifically, the most similar training protein would be the human tyrosine-protein kinase BTK, BTK in short, UniProt ID: Q06239) with sequence identity at 28%. And the most similar training compound would be K60 (HET ID) with Tanimoto score at 0.12. Indeed our results (Table 3) showed that contact AUPRC is just around 0.053. Given the explanation to interpret AUPRC and top-10 contact precision in the first case study, one would notice that the AUPRC value is 14-fold of the baseline (0.004) and 40% of our top-10 contact predictions were true positives (a level close to average protein-ligand docking performances).

As seen in Figure 9, again, our contact predictions are more focused in or near the binding site compared to the competing methods, which can be attributed to our structure-aware attention regularization (and supervision). A closer look into the false positives reveal more into our methods. Take DeepAffinity+ as an example. Among the 6 false-positive contact predictions, 4 were pairing correct binding residues with wrong compound atoms, 1 was paired to a protein residue that is a close sequence neighbor (2 residues away) of a correct binding residue, and 1 was paired to a protein residue that is not present in the co-crystal structure but predicted to be spatially close to a correct binding residue. In other words, the origins of false positives in contact prediction include (but are not limited to) pairing with other (nearby) compound atoms and pairing with sequential or predicted spatial neighbors of protein binding-residues. When the criterion of native contacts is relaxed from direct, first-shell contacts within 4Å to more contacts within longer distance cutoffs, the precision level would further increase, which is detailed next.

**Figure 9:**
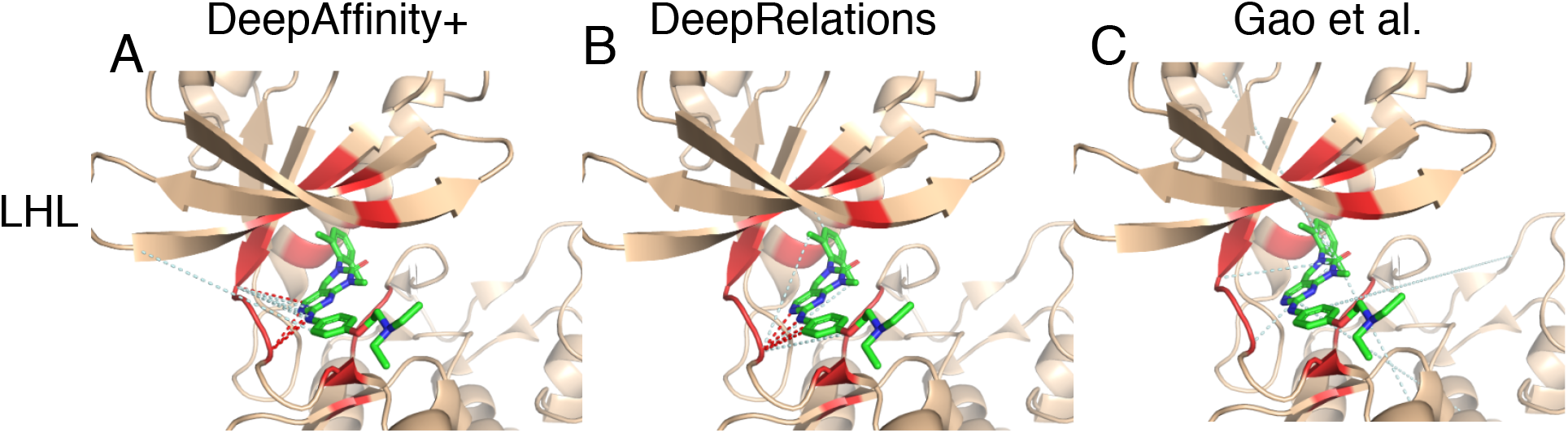
Structural visualization of top-10 intermolecular contacts predicted by (A) DeepAffinity+, (B) DeepRelations and (C) Gao et al. for a difficult test case. Here both the compound (LHL, in sticks) and the protein (tyrosine-protein kinase Lck, in wheat cartoons with binding residues highlighted in red) are new and very dissimilar to training data. The red and pale cyan dashed lines represent correct and incorrect top-10 predicted contacts. DeepAffinity+ and DeepRelations still managed to achieve the precision of 40% in their top-10 contact predictions.

#### Global patterns of top-10 predicted contacts

We extended the analysis of the patterns of predicted contacts to all test cases. Considering that the native contacts are defined strictly as direct, first-shell contacts within 4Å, we assess 4~10-Å distance-distributions of residue-atom pairs predicted by DeepAffinity+ (HRNN-GCN_cstr_sup) and DeepRelations in comparison with Gao et al.. As seen in the global analysis in Figure 10 and Table S10, DeepAffinity+ and DeepRelations significantly outperform the competing method in all distance ranges over all test sets. Specifically, among their top-10 contact predictions, around 40% for the default test and new-compound sets were first-shell contacts within 4Å and the ratios increased to about 70% when considering contacts within 10Å. For the more challenging cases of new-protein and both-new sets, the ratios were around 20% and 50%, respectively. These results significantly outperformed the competing method whose such ratios were merely 4~6% over all sets. Between our two models, DeepRelations behaved similarly as DeepAffinity+ and had more top-10 predictions falling in the long range of 8~10Å.

**Figure 10:**
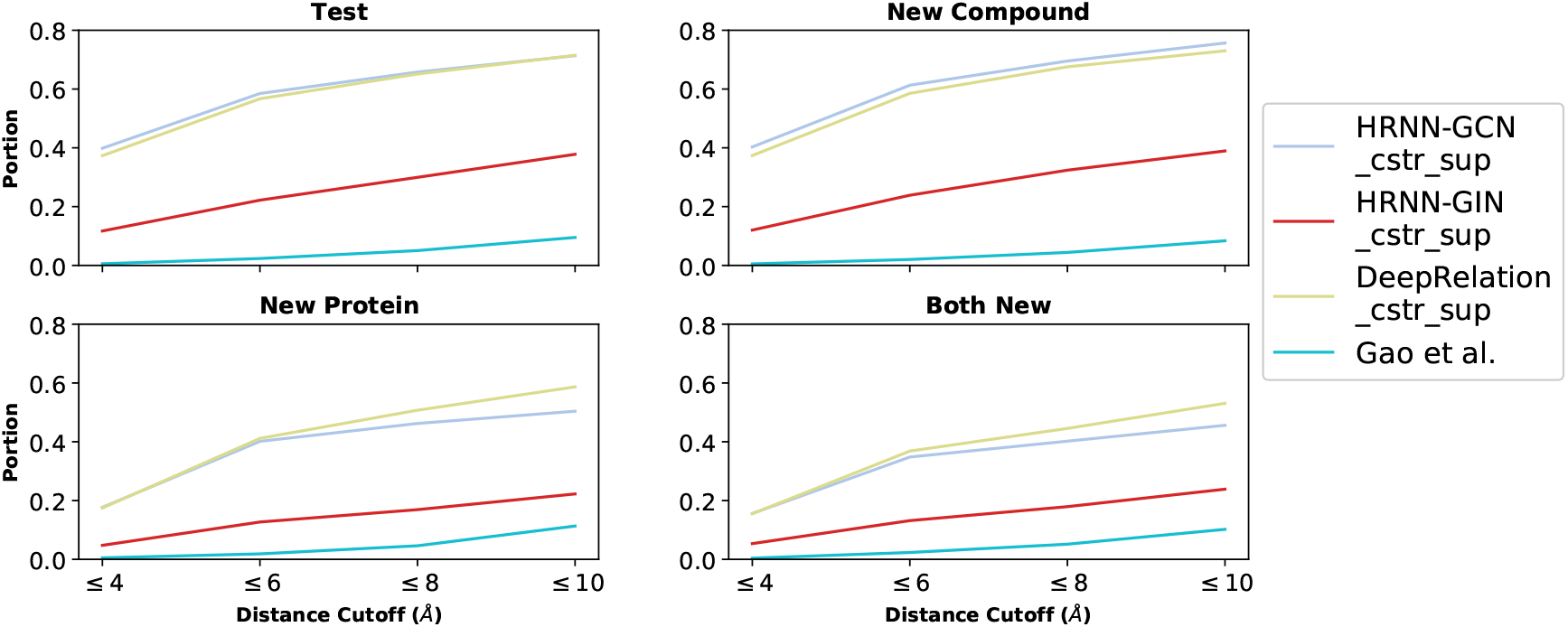
Distributions of top-10 contacts, predicted by DeepAffinity+, DeepRelations, and Gao’s method, in various distance ranges.

#### Predicted contacts assist and improve protein-ligand docking

From the case studies and the global analysis above, we have concluded that top-10 contact predictions by our methods are enriched with native contacts within 4 Å (20~40%) as well as dominated by longer-range “contacts” within 10 A (50~70%). We therefore test how much the top-10 contact predictions, including false positives, could make a positive impact in the drug discovery process. Picking a typical task - protein-ligand docking and a popular tool – AutoDock Vina,^68^ we assess how our contact predictions could assist the task by reducing the search space.

Specifically, we chose the five case studies (except the case where DeepRelations made no correct contact prediction) and performed unbound protein-ligand docking (all protein structures are unbound except PYGM whose structure is co-crystalized with its cognate phosphate AMP). Each pair (rigid protein and flexible ligand) is docked twice: one with the default procedure to define a search “box” covering the entire protein, and the other using a restricted box that barely covers *all* residues in the top-10 DeepRelations contact predictions (including false positives) and then has 20 Å-padding. All the other docking parameters in AutoDock Vina are default, including a total of 9 protein-ligand complex models ordered and reported at the end. Docking performances were evaluated by ligand RMSD of the top few models using the software DOCKRMSD.^70^

Results in Table 4 show that AutoDock Vina assisted by DeepRelations top-10 contact predictions had much improved docking performance compared to otherwise. When the top-10 contact precision was 40%, 50%, 50%, and 100%, respectively, the best ligand RMSD (among all 9 complex models) reduced from 2.77 Å, 4.01 A, 16.62 Å, and 18.75 Ådown to 2.45 Å, 1.59 Å, 4.73 A, and 1.88 Å, respectively. The quality of the top-1 models also drastically improved in 3 of 4 cases. Although the way to incorporate predicted contacts into protein-ligand docking remains to be optimized, these results have proved that the precision and spatial pattern of our structure-free contact prediction is at a level useful to assist and improve structure-based protein-ligand docking for pose prediction.

**Table 4:** Ligand docking performances for case studies. The default Autodock Vina is compared with that assisted by DeepRelations top-10 contact predictions.

| Protein<br>(UniProt, PDB) | Ligand | Complex<br>PDB | Top-10 Contact<br>Precision | RMSD (Å) – Vina |  |  |  | RMSD (Å) – Contact-Assisted Vina |  |  |  |
| --- | --- | --- | --- | --- | --- | --- | --- | --- | --- | --- | --- |
|  |  |  |  | Top 1 | Top 3 | Top 5 | Best | Top 1 | Top 3 | Top 5 | Best |
| LCK (P06239, 3LCK) | LHL | 3KMM | 40% | 3.02 | 3.02 | 2.77 | 2.77 | 4.65 | 2.98 | 2.45 | **2.45** |
| CA2 (P00918, 2CBA) | AL1 | 1BNN | 50% | 18.55 | 16.62 | 16.62 | 16.62 | 4.78 | 4.78 | 4.73 | **4.73** |
| CA2 (P00918, 2CBA) | IT2 | 3P5A | 50% | 15.98 | 15.98 | 4.01 | 4.01 | 3.65 | 3.65 | 3.65 | **1.59** |
| PYGM (P00489, 8GPB) | T68 | 3ZCU | 100% | 36.40 | 18.75 | 18.75 | 18.75 | 9.08 | 2.23 | 2.23 | **1.88** |

#### Affinity prediction for target prioritization

Using the two aforementioned CA2 (human carbonic anhydrase II) compounds (AL1 and IT2) in the first case study, we also explore the utility of our models for target prioritization for given compounds. As no affinity data was observed in our dataset for AL1 or IT2 with proteins other than CA2, we approximate the set of “off-targets” with all the 1,286 non-CA2 proteins in our dataset. For either compound AL1 and IT2, we assessed the distribution of its off-target affinities predicted by DeepAffinity+ and compared the distribution (see Figure S8) to its predicted on-target (CA2) affinity. As shown in Figure S8, 83.1% (100%) and 88.2% (99.7%) of predicted off-target affinities are weaker than the predicted (actual) affinity to the target CA2, for compounds AL1 and IT2, respectively. Removing CA2 homologs (4 in total) from the non-CA2 proteins led to nearly the same results (data not shown). We note that this case is particularly challenging because no homologs of the target CA2 are in the training set and the errors of target affinity prediction are higher than average. More systematic and dedicated studies are needed for this topic in future.

### More utilities from explainable affinity prediction

In the last part of the results, we explore additional utilities of our methods toward facilitating drug discovery: binding-site prediction for proteins and structure-activity relationship (SAR) for compounds. Our methods do not demand protein structures or protein-ligand docking to make these predictions. Rather, they simply aggregate predicted attentions (or predicted weights of residue-atom contacts) or/and decompose predicted affinities. Although not directly designed or optimized for these tasks, our explainable models have shown promising potentials in the tasks toward rational drug discovery.

#### Binding site prediction

The first extended utility we aim at is structure-free and ligand-specific binding-site prediction for proteins. To this end, we feed an arbitrary pair of protein and compound to the trained DeepAffinity+ and DeepRelations models and predict the weights of residue-atom pairs (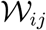 where *i* and *j* are the indices of a protein residue and a compound atom, respectively). We then calculate the max-marginal attention (max_*j*_ 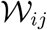) for each residue *i* as a weight for ranking. The performances of the residue weights toward ligand-specific binding site prediction are summarized in Figure 11 and Table S11. Here binding-site residues of a protein are strictly defined as those making direct, first-shell contacts with a paired compound. Without the help of protein structures, predicted residue-contact maps, or proteinligand docking, our methods on average achieved AUPRC (AUROC) of around 0.43 (0.77) for the default test and new-compound sets as well as AUPRC (AUROC) of around 0.18 (0.69) for the more challenging new-protein and both-new sets. In contrast, the competing method had AUPRC and AUROC close to the random performances of 0.004 and 0.50, respectively.

**Figure 11:**
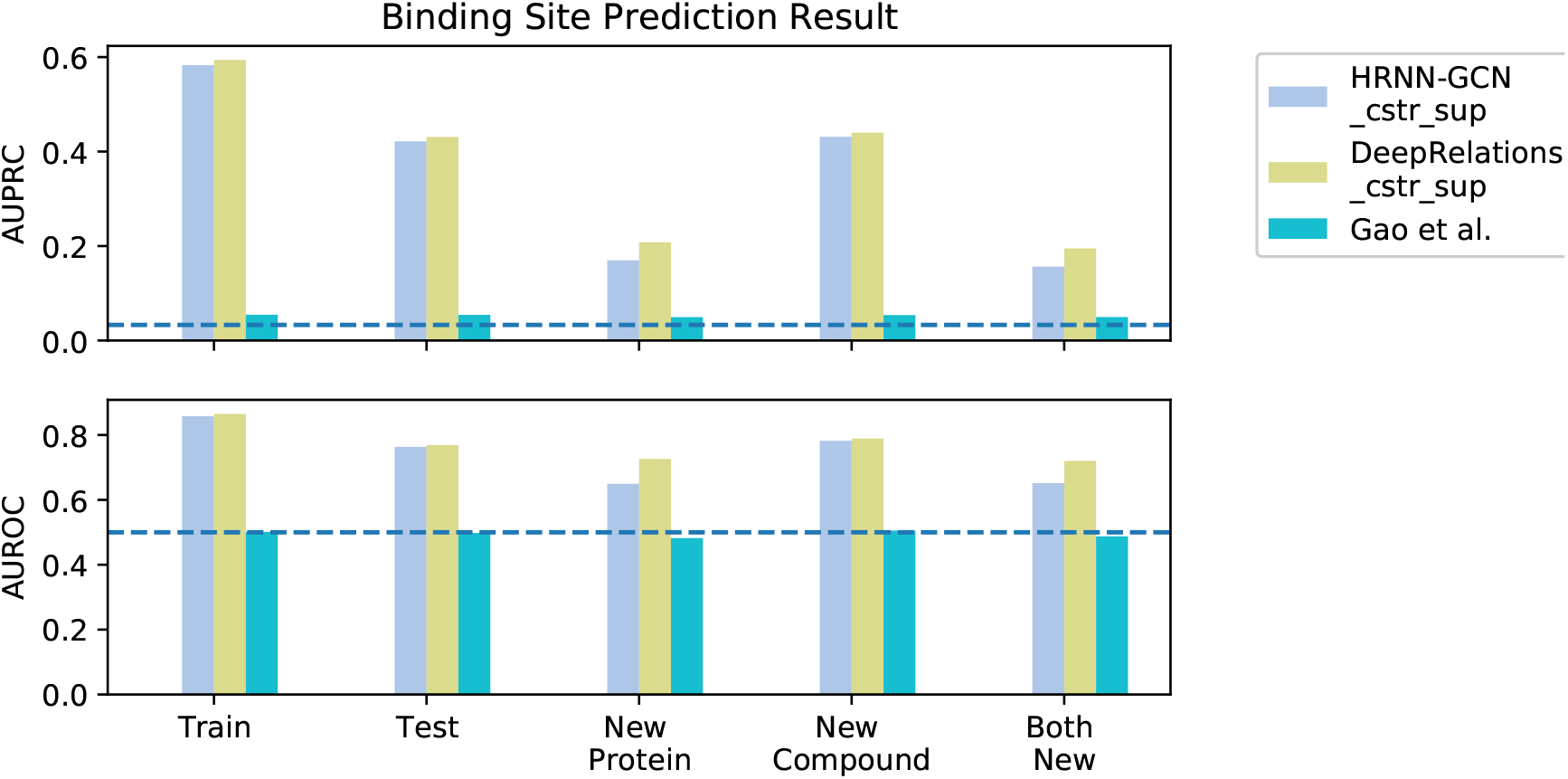
Comparing three interpretable methods (DeepAffinity+, DeepRelations, and Gao et al.) in binding-site prediction.

Between DeepAfinity+ and DeepRelations, we noticed that the latter had better performance in predicting binding sites for new proteins. Specifically, the AUPRC (AUROC) increased from 0.17 (0.65) to 0.21 (0.73) for the new-protein set and did from 0.16 (0.65) to 0.20 (0.72) for the both-new set.

#### Structure activity relationship (SAR)

The second extended utility we aim at is SAR for compounds. To test the utility we choose two subchallenges (SC3 and SC4) from Grand Challenge 3 of D3R^71^, Janus kinase 2 (JAK2) and Angiopoietin-1 receptor (TIE2), that were excluded in our training set (thus new proteins). The most similar proteins to JAK2 and TIE2 in our training set are calcium/calmodulin-dependent protein kinase kinase 2 (CAMKK2, sequence identity 48%) and cyclin-dependent kinase 2 (CDK2, sequence identity 39%), respectively. The two datasets include 17 and 18 congeneric compounds, respectively, with *K_d_* values measured. They were meant to “detect large changes in affinity due to small changes in chemical structure” (https://drugdesigndata.org/about/grand-challenge-3). In other words, the datasets focus on the sensitivity of methods targeting SAR. Chemical graphs, actual p*K_d_*, and DeepRelations-predicted p*K_d_* of the JAK2 and TIE2 compounds are in Figure S9 and S10, respectively.

Here we compare our DeepAffinity+ and DeepRelations not only to structure-free Gao et al. but also to 18 structure-based methods from the community that participated in the sub challenges. The assessment metrics for affinity ranking are Kendall’s *τ* and Spearman’s *ρ* as in D3R. A summary of the performances is in Table 5. In the case of JAK2, the 18 structure-based methods had *τ* ranging from 0.71 to −0.56 and ρ ranging from 0.86 to −0.70, including 8 methods with negative *τ* and *ρ* (see details in Table S13). As to the structure-free affinity predictors, Gao et al. had *τ* = −0.42 and *ρ* = −0.54 whereas our DeepAffinity+ had slightly better *τ* = −0.36 and ρ = −0.47, both outperforming just 1 structure-based method. However, our DeepRelations achieved *τ* = 0.15 and *ρ* = 0.21, outperforming 12 (two-thirds) of the structure-based methods. In the case of TIE2, the 18 structure-based methods had *τ* ranging from 0.57 to −0.57 and *ρ* ranging from 0.76 to −0.69, including 8 methods with negative *τ* and *ρ* (see details in Table S14). Interestingly, the best structure-based method for JAK2 only placed 12^th^ among 18 with slightly negative *τ* and *ρ* for TIE2. In contrast, all the structure-free affinity predictors performed well for TIE2: Gao et al., DeepAffinity+, and DeepRelations had *τ* (*ρ*) reaching 0.60 (0.74), 0.65 (0.79), and 0.61 (0.72), respectively; and they all outperformed the best structure-based method. The scatter plots of actual versus our predicted p*K_d_* are in Figure S11. We note that all 18 structure-based methods used crystal structures of proteins and often-expensive ligand docking whereas structure-free methods did not. Our methods only cost a fraction of a second when making quality predictions for tens to hundreds of compound-protein pairs, thus a useful complement to structure/docking-based methods toward virtual screening.

**Table 5:**
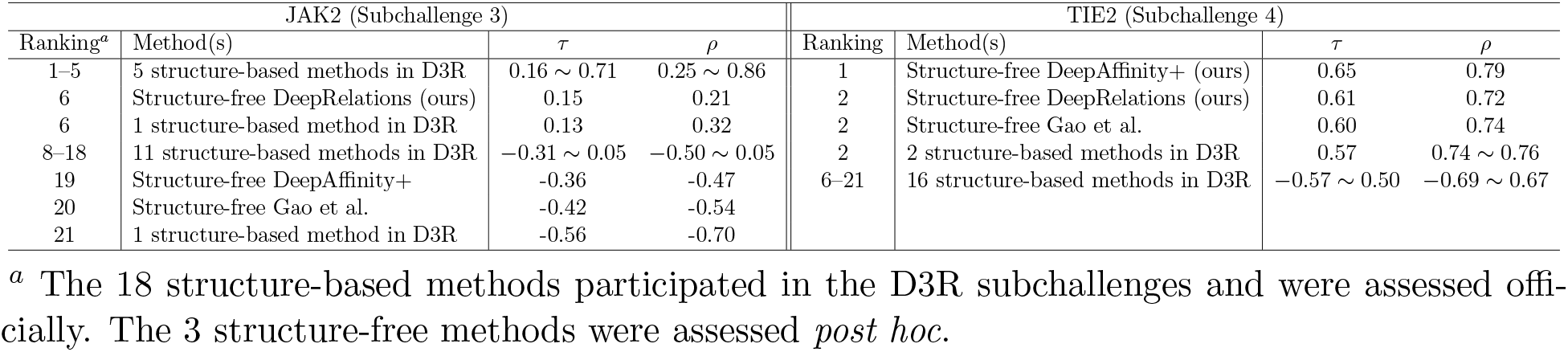
Summary of scoring performances among three structure-free methods (including our DeepAffinity+ and DeepRelations and eighteen structure-based methods.

| JAK2 (Subchallenge 3) |  |  |  | TIE2 (Subchallenge 4) |  |  |  |
| --- | --- | --- | --- | --- | --- | --- | --- |
| Ranking<sup>a</sup> | Method(s) | $\tau$ | $\rho$ | Ranking | Method(s) | $\tau$ | $\rho$ |
| 1-5 | 5 structure-based methods in D3R | 0.16 ~ 0.71 | 0.25 ~ 0.86 | 1 | Structure-free DeepAffinity+ (ours) | 0.65 | 0.79 |
| 6 | Structure-free DeepRelations (ours) | 0.15 | 0.21 | 2 | Structure-free DeepRelations (ours) | 0.61 | 0.72 |
| 6 | 1 structure-based method in D3R | 0.13 | 0.32 | 2 | Structure-free Gao et al. | 0.60 | 0.74 |
| 8-18 | 11 structure-based methods in D3R | -0.31 ~ 0.05 | -0.50 ~ 0.05 | 2 | 2 structure-based methods in D3R | 0.57 | 0.74 ~ 0.76 |
| 19 | Structure-free DeepAffinity+ | -0.36 | -0.47 | 6-21 | 16 structure-based methods in D3R | -0.57 ~ 0.50 | -0.69 ~ 0.67 |
| 20 | Structure-free Gao et al. | -0.42 | -0.54 |  |  |  |  |
| 21 | 1 structure-based method in D3R | -0.56 | -0.70 |  |  |  |  |

<sup>a</sup> The 18 structure-based methods participated in the D3R subchallenges and were assessed officially. The 3 structure-free methods were assessed *post hoc*.

Beyond affinity scoring we further examine DeepRelations in extracting SAR knowledge toward drug discovery. A central question in lead optimization is where and how to modify a lead compound to improve its property (affinity here). As a stepping stone, we construct predictors from our DeepRelations in order to anticipate the affinity changes when a functional-group substituent is introduced to a lead. Specifically, we regard our predicted 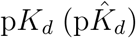 as estimated binding energy for a compound-protein pair and our predicted jointattention 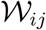 as the fraction of contribution between protein residue *i* and compound atom *j*. Borrowing the idea of energy decomposition, we calculate the binding-energy contribution of a functional group *R* as the product of the predicted binding-energy and the sum-marginals of joint attention: 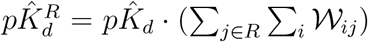. In this way the difference of this *R*-group contribution, 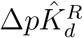, can be a predictor of affinity change when introducing a substituent *R*-group to a compound.

To test our predictor for lead optimization, we use the JAK2 dataset involving 17 compounds that share a common scaffold and have distinct combinations of two functional groups (3 choices for *R*_1_ and 10 for *R*_2_, see Figure 12A and Figure S9). We construct 121 pairs of compounds between a weaker binder (origin) and a stronger binder (end). 7, 36, and 78 of the structural changes from the origin to the end compound involve *R*_1_, *R*_2_, and both-*R* substitutions, respectively. And we compare three methods in predicting these 121 affinity changes with assessment metrics including Pearson’s *r* (main assessment), Spearman’s *ρ* and Kendall’s *τ* (Figure 12B-D). A straightforward predictor using DeepRelations’ △*pK_d_* without decomposition had *r* = 0.218; whereas the decomposed affinity-change predictor 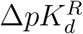 improved *r* to 0.361. If one has access to the protein in complex with a previously discovered compound and can have an accurate estimate of the binding residues, the summation of protein residue *i* in 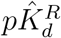 can be just over binding residues rather than all residues. In that case, the new 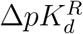 can slightly improve r further to 0.363. Our decomposed affinity-change predictor 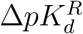 similarly improved *ρ* and *τ*. Compared to the 18 structure-based competing methods that participated in the D3R JAK2 subchallenge, our structure-free predictor with decomposition outperformed 15 (five sixths) of them in *r, ρ*, and *τ*, as detailed in Tables S15 and S16.

**Figure 12:**
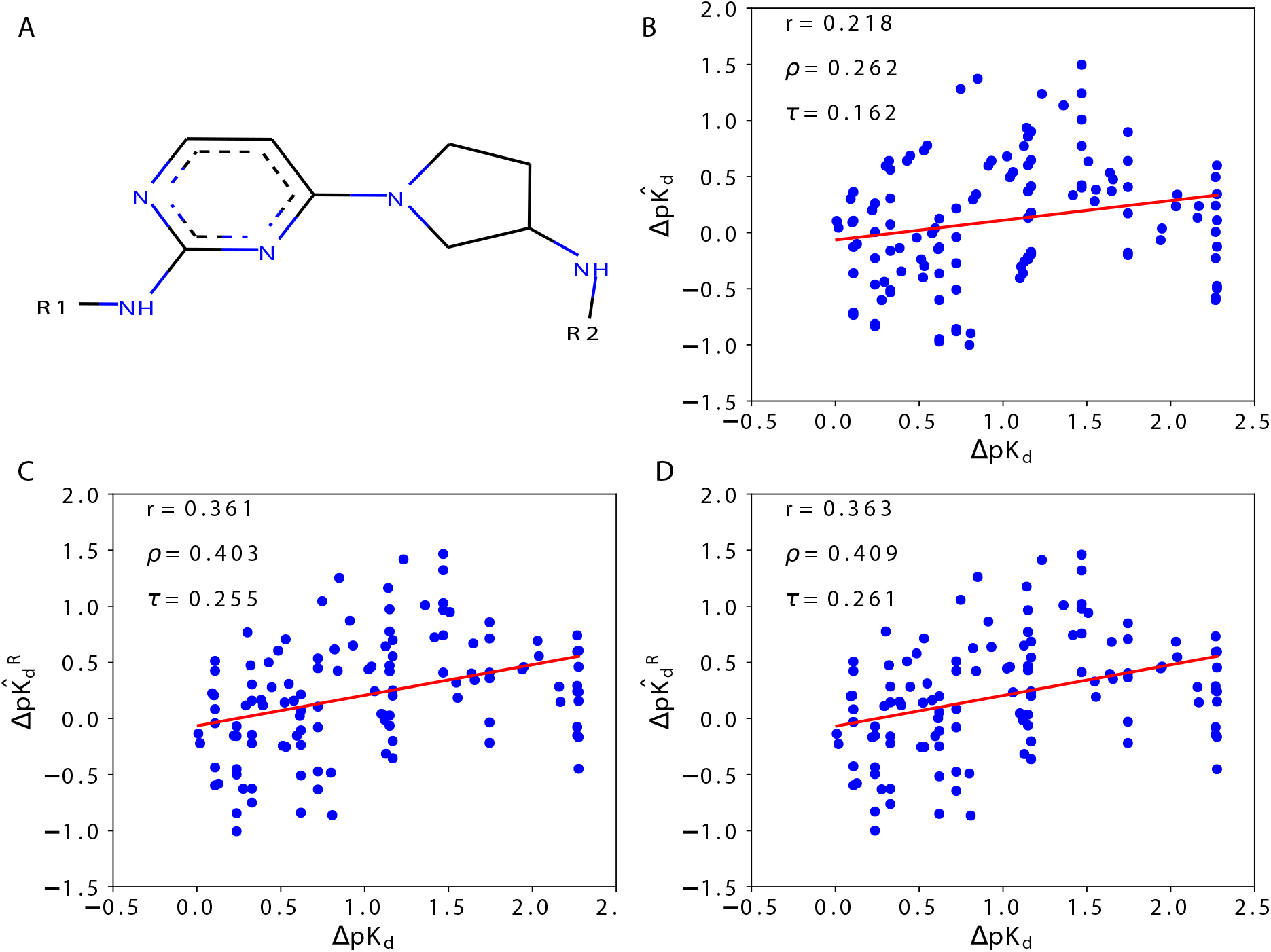
Actual (*x*-axis) *versus* DeepRelations-predicted (*y*-axis) affinity changes when introducing functional-group substitutions (*R*_1_, *R*_2_ or both in A) to lead compounds for JAK2. The three predictors are: B. predicted affinity change 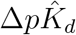; C. group-decomposed affinity change 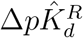 using all protein residues and the substituent group *R* alone; and *D*. group-decomposed affinity change 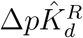 using estimated protein binding residues and the substituent group *R* alone.

When we split the analysis into 3 series involving *R*_1_, *R*_2_, and both-*R* separately, we observed that 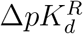 improved r from 0.267 to 0.753, −0.081 to 0.137, and 0.244 to 0.377, respectively (Figure S13). Using the binding-residue information could slightly improve the correlation further. Interestingly, when both *R*-groups are substituted (78 cases), 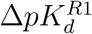 had a better Pearson’s correlation (0.405) with the actual affinity changes than 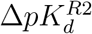 (−0.121) and even 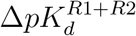 (0.375) did (Figure S14), potentially suggesting that *R*_1_ group could be explored first for affinity optimization. Once a functional group *R* is chosen, affinity changes upon any proposed substitution can be predicted using our group-decomposed 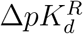.

## Conclusions

Toward accurate and interpretable machine learning for structure-free prediction of compound-protein interactions, we have curated compound-protein interaction dataset annotated with both affinities and intermolecular atom-contacts, assessed the adequacy of current attention-based deep learning models for both accuracy and interpretability, and developed novel machine-learning models (in particular, DeepAffinity+ and DeepRelations) to remarkably enhance interpretability without sacrificing accuracy. We have also shown that our methods’ accuracy for affinity prediction is comparable or better than competing (non-interpretable) methods using established benchmark datasets. This is the first study with dedicated model development and systematic model assessment for interpretability in affinity prediction.

Our study has found that commonly-used attention mechanisms alone, although better than chance in most cases, are not satisfying in interpretability. The most attended proteinligand contacts in affinity prediction do not reveal native contacts underlying affinities at a useful level. The conclusion maintains regardless of the representation of molecules (sequences/strings or graphs) the architecture of neural networks. We have tackled the challenge with three innovative, methodological advances. First, we introduce structure-aware constraints to regularize attentions (or guide their sparsity patterns), using sequence-predicted structural contexts such as protein surfaces and protein residue-residue contact maps. Second, we exploit available native contacts to supervise novel joint attentions, i.e., to teach neural network how to weigh residue-atom pairs when making affinity predictions. Lastly, we build intrinsically explainable model architecture where various atomic relations, reflecting physics laws, are explicitly modeled and aggregated for affinity prediction. Joint attentions are embedded over residue-atom pairs for individual and overall relations. And a multi-stage hierarchy, trained end-to-end, progressively focuses attentions on protein surfaces, binding *k*-mers and residues, and residue-atom contact pairs. The first two advances are introduced in both DeepAffinity+ and DeepRelations; and the last is additionally introduced in DeepRelations. Their best versions involve hierarchical recurrent neural networks (HRNN) to embed protein sequences and graph convolutional networks (GCN) to embed compound graphs.

Empirical results demonstrate the superiority of DeepAffinity+ and DeepRelations in interpretable and accurate prediction of compound-protein interactions. Their affinity prediction shows generalizability to compounds or/and proteins that are new or even dissimilar to training data. Compared to a competing interpretable method, they boosted the AUPRC for contact prediction (a measure of interpretability) by around 33, 10, 35, and 9-fold for the default test, new-protein, new-compound, and both-new sets, respectively. Importantly, improved model interpretability has shown to contribute to improved model accuracy and generalizability.

Case studies suggest that DeepAffinity+ and DeepRelations predict not only more correct but also more well-patterned contacts that are focused in or near binding sites, which is thanks to the structure-aware regularization and supervision of joint attentions. A global analysis indicates that around 40% (20%) of our top-10 predicted contacts are native contacts that are direct and first-shell for the test and the new-compound set (the new-protein and both-new set). Many “incorrect” predictions due to the strict definition of native contacts were within reasonable ranges — in fact, around 70% (50%) of the top-10 predicted contacts correspond to residue-atom pairs within 10 Å when the set does not (does) involve a new protein. With the precision level and the focused pattern, our top-10 contact predictions (including false positives) have demonstrated their value in assisting and improving proteinligand docking, while the protocol to incorporate the predictions into docking remains to be optimized.

By aggregating joint attentions and decomposing predicted affinities, we also demonstrate additional utilities of our explainable affinity and contact predictor, toward drug-discovery tasks such as binding site prediction, SAR (scoring) and SAR (lead optimization). Although not directly designed nor optimized for these tasks, our methods and analyses have shown great potentials in these tasks toward facilitating drug discovery.

An additional benefit of our structure-free methods is their broad applicability toward the vast chemical and proteomic spaces. They do not rely on 3D structures of compound-protein complexes or even proteins alone when such structures are often unavailable. The only inputs needed are protein sequences and compound graphs. Meanwhile, they adopt the latest technology to predict structural contexts from protein sequences (such as surfaces, secondary structures, and residue-residue contact maps). And they introduce structure-aware regularization to incorporate the predicted structural contexts into affinity and contact predictions. When structure data are available, DeepRelations can readily integrate such data by using actual rather than predicted structural contexts. We tested the use of actual versus predicted protein residue-residue contact maps and did not observe significant performance differences in our cases (Table S12).

Our study demonstrates that, it is much more effective to directly teach explainability to machine learning models (such as our structure-aware regularization and supervision of joint attentions) and build explainability into model architectures (such as our explicit modeling of atomic relations in DeepRelations) than to demand explainability from general-purpose models (such as seeking contact-interpretation from unsupervised, generic attention mechanisms). In other words, designing intrinsically interpretable machine learning models incorporated with domain knowledge, although more difficult, can be much more desired than pursuing interpretability in a *post hoc* manner.

## Associated Contents

### Supporting Information

The Supporting Information is available free of charge at http://pubs.acs.org.

Methods including data curation, compound pre-processing, protein residue-residue contact map prediction, compound similarity calculation, compound property distributions, and atom features for compounds. Results including numeric details for Figures 4–6, ablation study for DeepRelations, numeric details for Figures 10–11, SAR performance details of individual methods, compounds in JAK2 and TIE2 subchallenges, and lead-optimization performance details.

## Acknowledgement

This work was supported in part by the National Institutes of Health (R35GM124952) and the National Science Foundation (CCF-1943008). Part of the computing time was provided by the Texas A&M High Performance Research Computing.

---

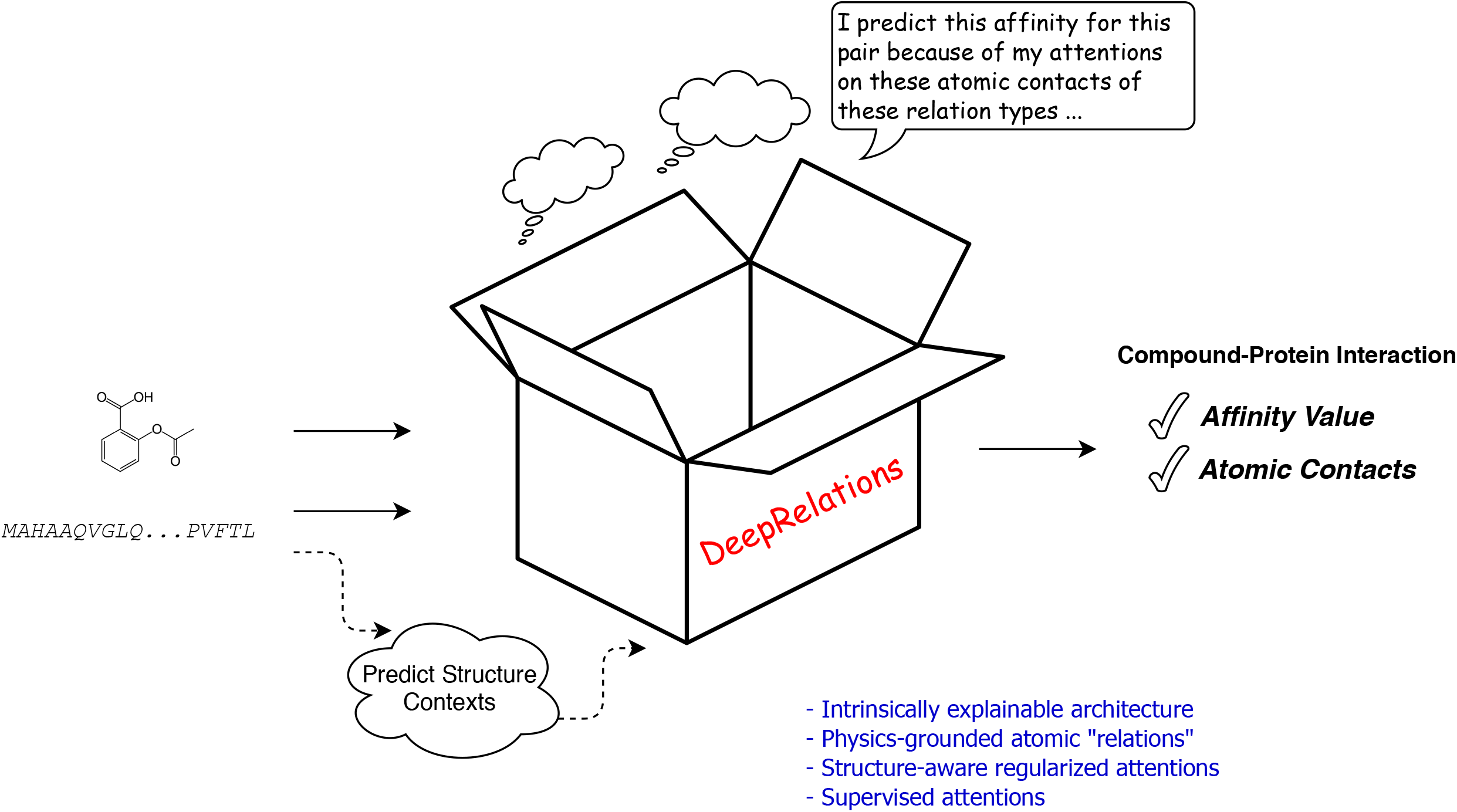

