## Supporting Information for "Explainable Deep Relational Networks for Predicting Compound-Protein Affinities and Contacts"

### 1 Data

#### 1.1 Data curation and splitting

During our training and testing, we used two groups of dataset. The first group is for benchmarking affinity prediction only. It includes three established benchmark datasets: Davis, KIBA and PDBbind. Then based on PDBbind and BindingDB dataset, we curated the second group with intermolecular contacts data for benchmarking both affinity and contact prediction (our current definition of accuracy and interpretability). In general, our data is composed of four portions: protein sequence, compound SMILEs, affinity labels and interolecular interaction/contact data (only used for assessing interpretability). To embed protein sequences, we used SPS formats in RNNs and FASTA sequences from UniProt in HRNNs or CNNs. To embed compounds, we used the canonical SMILES from Pubchem in RNNs and RDKit-converted graphs in GCNs or GINs. Cases are removed when multiple ligands simultaneously interact with one protein or protein/compound identities are unclear.

All of our interaction data are curated from the LigPlot service of PDBsum(Laskowski *et al.*, 2018). PDBsum provides an overview of each 3D macromolecular structure deposited in the Protein Data Bank. Based on the HET ID of the compound and the PDB ID of the protein (complex), we downloaded corresponding interaction data from PDBsum for the compound-protein pair. PDBsum only provides the interaction list. So we mapped each residue of the protein (each atom of the compound) between FASTA sequences (SMILES and graphs) to that in the list file derived from PDB files. After this, we converted the list to a native contact matrix with indexed protein residues and compound atoms. During this process, we removed cases which can't be mapped because of incomplete information in PDB files.

**Davis**: The Davis dataset (He *et al.*, 2017) contains all 30,056 Kd-labeled pairs between 68 kinase inhibitors (including FDA-approved drugs) and 442 kinases, randomly split into 25,046 for training and 5,010 for testing.

**KIBA**:The Kinase Inhibitor BioActivity (KIBA) dataset (Tang *et al.*, 2014) contains 118,254 pairs between 2,111 kinase inhibitors and 229 kinases,including 98,545 for training and 19,709 for testing.

**PDBbind**: We downloaded the v2019 of PDBBIND database (Liu *et al.*, 2015). PDBbind database contains 17,679 protein-ligand complexes which are referred to as the "general set". A "refined set" has been compiled which includes better quality samples in comparison to the "general set". The refined set consists of 4,852 protein-ligand complexes with $K_i/K_d$ affinity labels. We utilized the UniProt (Consortium, 2015) Retrieving API (`https://www.uniprot.org/uploadlists/`) to retrieve the UniProt FASTA sequences (canonical) and GO terms for proteins involved. Some cases are with the same pair of UniProt ID and compound but with different labels, which could be due to protein mutations and removed here. In addition, we set 1,000 and

100 as the length-cutoffs for protein sequence and compound SMILES, respectively, to remove extremely long sequences/strings.
After filtering, we have 3,505 pairs between 1149 proteins and 2870 compounds. The pairs include 62 cases involving nuclear receptors (6 of them are estrogen receptors [ER]), 33 involving G protein-coupled receptor(GPCR), 106 involving ion channels and 2,157 involving enzymes (EC1: 77, EC2: 557, EC3: 1,014, EC4: 400, EC5: 78, EC6: 49, EC7: 2 overlapping; 72 kinases). They are randomly split into 2,921 pairs for training and 584 for testing affinity alone.
**BindingDB**: BindingDB(Liu *et al.*, 2006) is a public, web-accessible database of measured binding affinities. We used part of data whose $K_i$ or $K_d$ label is available from the previously-curated BindingDB dataset(Karimi *et al.*, 2019). In our previous $K_d$-labeled set, there were 8,778 pairs for the training set, 3,811 for testing, 4 for ER, 2554 for GPCR, 366 for ion channels and 2306 for kinases, only 721 among which is found to have interaction data. In our previously curated $K_i$-labeled set, there were 101,135 pairs for the training set, 43,392 for testing, 516 for ER, 77,994 for GPCR, 8,101 for ion channels and 3,354 for kinases, only 1,627 of which is now found to have interaction data.
**Curated dataset for affinity and contact prediction:** Firstly, we merged PDBbind dataset and BindingDB dataset and applied 1,000 and 100 as the length-cutoffs for protein sequences and compound SMILES again. After generating their interaction data from PDBsum, we filtered out cases which didn't have corresponding contact information. In the end, we reached 4,446 pairs, with both affinity ($K_i$ or $K_d$) and contacts (as in crystal structures of protein-ligand complexes) available, between 1,287 proteins and 3,672 compounds. The pairs include 105 interactions with nuclear receptors (21 of which are with ER), 89 with G protein-coupled receptor(GPCR), 111 with ion channels and 2,913 with enzymes (including 114 with kinases). In particular, those pairs involving enzymes can be split (with overlaps) into EC classes: 222 (EC1), 865 (EC2), 1,218 (EC3), 500 (EC4), 92 (EC5), 52 (EC6) and 2 (EC7).
The set is randomly split into four folds where fold 1 do not overlap with fold 2 in compounds, do not do so with fold 3 in proteins, and do not do so with fold 4 in either compounds or proteins. Folds 2, 3, and 4 are referred to as new-compound, new-protein, and both-new sets for generalizability tests; and they contain 521, 795 and 205 pairs, respectively. Then we randomly split fold 1 into training (2,334) and test (591) sets. During this process, we followed the algorithm in 1. Ideally, we expect to split 70% cases into training + test sets and at least 5% into both-new. In addition, we aim to balance the compound and protein unique set. In this case, the ideal percentage of new-compound and new-protein sets would be 10% - 15%.

#### 1.2 Compound preprocessing

At first we used open babel software (O'Boyle *et al.*, 2011) for ionization of compounds. Then, We used "chem.SanitizeMol()" from RDkit open source software (Landrum *et al.*, 2006) for compound sanitization, which includes converting certain neutral atoms to Zwitterionic forms or aromatic atoms to Kekulized forms where applicable. Citing the RDKit book (`https://www.rdkit.org/docs/RDKit_Book.html#molecular-sanitization`), the detailed steps are:

- "Standardizes a small number of non-standard valence states which include ionization.
- Calculates the explicit and implicit valences on all atoms.
- Symmetrized based on smallest set of smallest rings algorithm
- Converts aromatic rings to their Kekule form.
- Determines the number of radical electrons
- Identifies the aromatic rings and ring systems
- Identifies which bonds are conjugated
- Calculates the hybridization state of each atom
- Removes chiral tags from atoms that are not sp3 hybridized.
- Adds explicit Hs where necessary to preserve the chemistry. This is typically needed for heteroatoms in aromatic rings."

**Algorithm 1** Data Splitting

```
Input: Data from PDBbind and BindingDB with Ki+Kd labels
Output: Five folds data sets: training set, test set and three generalization sets
Initialize: ProteinPercentage and CompoundPercentage as 15%,
Initialize: Result as Empty
Initialize: Iter = 0
while Result is Empty do
  while Iter < 5 do
    Shuffle Data
    Randomly Initialize ProteinPercentage protein from Protein List to UniqueProteinList
    Randomly Initialize CompoundPercentage proteins from Compound List to UniqueCompoundList
    for Each case in Data do
      if Protein in UniqueProteinList and Compound in UniqueCompoundList then
        Store case in DoubleUniqueSet
      else if Protein in UniqueProteinList then
        Store case in ProteinUniqueSet
      else if Compound in UniqueCompoundList then
        Store case in ComoundUniqueSet
      else
        Store case in Rest
      end if
    end for
    Randomly split 20% of Rest to TestSet and 80% to TrainingSet
    if Percentage of Rest ∈ [65%, 70%] and Percentage of DoubleUniqueSet ∈ [5%, 10%] then
      Store splitting result to Result
    end if
    Iter = Iter + 1
  end while
  if Percentage of ProteinUnique > 15% then
    ProteinPercentage = ProteinPercentage - 1%
  else if Percentage of ProteinUnique < 10% then
    ProteinPercentage = ProteinPercentage + 1%
  end if
  if Percentage of CompoundUnique > 15% then
    CompoundPercentage = CompoundPercentage - 1%
  else if Percentage of CompoundUnique < 10% then
    CompoundPercentage = CompoundPercentage + 1%
  end if
  Iter = 0
end while
return Splitting result with most cases in DoubleUniqueSet among Result
```

#### 1.3 Predicting protein residue-residue contact maps

We used the standalone software RaptorX-contact for contact map prediction that is eventually for distance-based protein structure prediction (Xu, 2019). The software was provided at the following link: https://github.com/j3xugit/RaptorX-Contact. RaptorX-contact utilized various 1D and 2D sequence information as features for their deep learning model. In the following, the feature generation for RaptorX-contact is explained in details:

- Protein sequence: primary sequence represented as a string of letters (upper case).

- Multiple sequence alignment (MSA): For each protein, we generated four MSAs by running HHblits (Remmert *et al.*, 2012) with 3 iterations and E-value set to 0.001 and 1 for UniClust30 library (Mirdita *et al.*, 2017) created in October 2017 and the UniRef sequence database (Suzek *et al.*, 2015) created in early 2018. For each MSA, we calculate the position-specific frequency matrix (PSFM) and position-specific scoring matrix (PSSM).

- Protein property prediction: For each of the protein sequences, we predict its disorder (Wang *et al.*, 2016a) its secondary structure (SS3 and SS8)through DeepCNF (Wang *et al.*, 2016b), and its solvent accessability through Acconpred (Ma and Wang, 2015).
- CCMpred: For each of the MSA, we predict its normalized contact map through CCMpred (Ekeberg *et al.*, 2013).
- MetaPSICOV: For each of the MSA, we calculate three matrices of pairwise relationship generated by alnstats in MetaPSICOV (Jones *et al.*, 2015).

In summary, 1 protein has 4 sets of input features and accordingly 4 predicted distance matrices, which are then averaged to obtain the final prediction.

#### 1.4 Compound similarity calculation

Molecular fingerprints provide a mathematical representation of compounds which is very useful for compound similarity calculation (Cereto-Massagué *et al.*, 2015). For virtual screening (and protein-ligand interactions), circular (that is topological) fingerprints are better (Hert *et al.*, 2004) such as Extended-connectivity fingerprints (ECFPs) (Rogers and Hahn, 2010). We have used the open source software RDkit (Landrum *et al.*, 2006) for implementation of ECFP4 with 1024 bits. We used Morgan fingerprint with radius 2 which is roughly equivalent to ECFP4 (Rogers and Hahn, 2010). To assess the similarity between molecular fingerprints, we used the standard Tanimoto/Jaccard coefficient (Cereto-Massagué *et al.*, 2015). Tanimoto score considers the common number of 1 bits divided by the total number of 1 bits. Tanimoto scores range between 0 (no similarity) and 1 (highly similar). 0.85 has been chosen as the Tanimoto coefficient threshold above which molecules will be considered similar enough (Martin *et al.*, 2002; Patterson *et al.*, 1996).

#### 1.5 Compound's property and label distribution

| Training set compare to | Test | New Protein | New Compound | Both New |
|---|---|---|---|---|
| LogP | 0.1707 | 0.1781 | 0.2350 | 0.3077 |
| Exact_MW | 0.1350 | 0.2304 | 0.2146 | 0.2906 |
| Label (Affinity) | 0.1721 | 0.2118 | 0.1581 | 0.3065 |

Table S1: Jensen-Shannon distances between the training and the other sets in various property distributions.

#### 1.6 Features used for compounds in graph-based representation

The following table describes the features used for the vertex (atoms) in graph representation (GCN/GIN) for DeepAffinity+ and DeepRelations:

| Feature name | type | length |
|---|---|---|
| Atom being inside aromatic ring | binary | 1 |
| Polarity (based on Gasteiger partial Charges) | continuous | 1 |
| Charge (based on formal Charges) | integer | 1 |
| Type of atoms (C,N ,O ,S ,F ,Si ,Cl ,P ,Br ,I ,B or unknown) | one hot encoding | 12 |
| Hydrogen bonding (F, N, O or not) | binary | 1 |
| Halogen bonding (F, Cl, I, Br or not) | binary | 1 |
| Degree of atom (adopted from (Tang *et al.*, 2020)) | one hot encoding | 6 |
| Number of hydrogen attached (adopted from (Tang *et al.*, 2020)) | one hot encoding | 5 |
| Implicit valence (adopted from (Tang *et al.*, 2020)) | one hot encoding | 6 |
| Radical electrons (adopted from (Tang *et al.*, 2020)) | integer | 1 |
| Hybridization (adopted from (Tang *et al.*, 2020)) | one hot encoding | 6 |

Table S2: Features used for compounds in graph-based representation of DeepAffinity versions

### 2 Results

#### 2.1 Attentions alone are inadequate for interpreting compound-protein affinity prediction.

| Model (Prot.-Comp.) | Assessment | | Training | Test | New-Prot. | New-Comp. | Both-New |
|---|---|---|---|---|---|---|---|
| RNN-RNN | affinity | RMSE | 0.53 | 1.57 | 1.66 | 1.40 | 1.66 |
| | | Pearson's $r$ | 0.97 | 0.65 | 0.39 | 0.70 | 0.49 |
| | contact | AUPRC | 0.0065 | 0.0067 | 0.0060 | 0.0061 | 0.0057 |
| | | AUROC | 0.5108 | 0.5078 | 0.5182 | 0.5053 | 0.5122 |
| RNN-GCN | affinity | RMSE | 0.49 | **1.40** | 1.68 | **1.28** | 1.82 |
| | | Pearson's $r$ | 0.98 | **0.72** | 0.35 | **0.74** | 0.36 |
| | contact | AUPRC | 0.0068 | 0.0069 | 0.0063 | 0.0063 | 0.0064 |
| | | AUROC | 0.5061 | 0.5031 | 0.5023 | 0.5021 | 0.5047 |
| CNN-GCN | affinity | RMSE | 1.02 | 1.49 | 1.72 | 1.33 | 1.71 |
| | | Pearson's $r$ | 0.85 | 0.67 | 0.38 | 0.72 | 0.45 |
| | contact | AUPRC | 0.0060 | 0.0064 | 0.0038 | 0.0060 | 0.0045 |
| | | AUROC | 0.5059 | 0.5049 | 0.4862 | 0.5004 | 0.4848 |
| HRNN-RNN | affinity | RMSE | 0.40 | 1.47 | 1.49 | **1.28** | 1.60 |
| | | Pearson's $r$ | 0.98 | 0.69 | 0.57 | **0.74** | 0.55 |
| | contact | AUPRC | 0.0067 | 0.0069 | 0.0049 | 0.0065 | 0.0052 |
| | | AUROC | 0.5025 | 0.5047 | 0.4946 | 0.5009 | 0.4934 |
| HRNN-GCN | affinity | RMSE | 0.69 | 1.47 | **1.46** | 1.34 | **1.49** |
| | | Pearson's $r$ | 0.93 | 0.70 | **0.56** | 0.73 | **0.61** |
| | contact | AUPRC | 0.0071 | 0.0069 | 0.0061 | 0.0065 | 0.0067 |
| | | AUROC | 0.5174 | 0.5143 | 0.5269 | 0.5085 | 0.5272 |
| HRNN-GIN | affinity | RMSE | 1.22 | 1.53 | 1.67 | 1.43 | 1.68 |
| | | Pearson's $r$ | 0.80 | 0.66 | 0.49 | 0.67 | 0.53 |
| | contact | AUPRC | 0.0067 | 0.0071 | 0.0048 | 0.0074 | 0.0051 |
| | | AUROC | 0.4936 | 0.4969 | 0.4693 | 0.4948 | 0.4688 |
| Gao et al. | affinity | RMSE | 1.79 | 1.87 | 1.72 | 1.75 | 1.79 |
| | | Pearson's $r$ | 0.73 | 0.58 | 0.42 | 0.51 | 0.42 |
| | contact | AUPRC | 0.0062 | 0.0060 | 0.0048 | 0.0057 | 0.0048 |
| | | AUROC | 0.5150 | 0.5157 | 0.5165 | 0.5150 | 0.5155 |

Table S3: Comparing accuracy and interpretability among various versions of DeepAffinity with (unsupervised) joint attention mechanisms as well as a state-of-the-art interpretable method (Gao et al., adapted from binding classifcation to affiity regression). Best affinity predictions in each set are highlighted in boldface.

#### 2.2 Supervising attentions significantly improves interpretability

| Model | Assessment | | Training | Test | New-Prot. | New-Comp. | Both-New |
|---|---|---|---|---|---|---|---|
| HRNN-GCN _cstr | affinity | RMSE | 0.44 | **1.42** | 1.61 | 1.34 | 1.72 |
| | | Pearson's $r$ | 0.95 | **0.71** | 0.40 | 0.73 | 0.32 |
| | contact | AUPRC | 0.0067 | 0.0065 | 0.0052 | 0.0064 | 0.0065 |
| | | AUROC | 0.5041 | 0.5036 | 0.5087 | 0.4996 | 0.5060 |
| HRNN-GCN _cstr _sup (Best DeepAffinity+) | affinity | RMSE | 0.64 | 1.49 | 1.57 | 1.34 | 1.61 |
| | | Pearson's $r$ | 0.95 | 0.68 | 0.45 | 0.73 | 0.51 |
| | contact | AUPRC | 0.3614 | **0.1974** | 0.0477 | **0.1998** | 0.0411 |
| | | AUROC | 0.8184 | 0.7378 | 0.6001 | 0.7380 | 0.5909 |
| HRNN-GIN _cstr | affinity | RMSE | 0.73 | 1.46 | 1.69 | **1.31** | 1.80 |
| | | Pearson's $r$ | 0.93 | 0.69 | 0.47 | **0.74** | 0.43 |
| | contact | AUPRC | 0.0084 | 0.0083 | 0.0064 | 0.0087 | 0.0066 |
| | | AUROC | 0.5187 | 0.5214 | 0.5109 | 0.5297 | 0.5186 |
| HRNN-GIN _cstr _sup | affinity | RMSE | 1.13 | 1.53 | 1.54 | 1.37 | **1.59** |
| | | Pearson's $r$ | 0.81 | 0.65 | **0.55** | 0.70 | **0.57** |
| | contact | AUPRC | 0.0698 | 0.0457 | 0.0113 | 0.0482 | 0.0135 |
| | | AUROC | 0.5999 | 0.5847 | 0.5066 | 0.5803 | 0.5157 |
| DeepRelations _cstr | affinity | RMSE | 0.43 | 1.51 | **1.52** | 1.41 | 1.67 |
| | | Pearson's $r$ | 0.97 | 0.67 | 0.52 | 0.69 | 0.47 |
| | contact | AUPRC | 0.0083 | 0.0091 | 0.0100 | 0.0086 | 0.0089 |
| | | AUROC | 0.528 | 0.533 | 0.558 | 0.533 | 0.552 |
| DeepRelations _cstr _sup (Best DeepRelations) | affinity | RMSE | 0.47 | 1.45 | 1.57 | 1.35 | 1.63 |
| | | Pearson's $r$ | 0.96 | 0.69 | 0.47 | 0.71 | 0.52 |
| | contact | AUPRC | 0.367 | 0.187 | **0.052** | 0.191 | **0.047** |
| | | AUROC | 0.845 | **0.760** | **0.669** | **0.764** | **0.659** |
| Gao et al. | affinity | RMSE | 1.79 | 1.87 | 1.72 | 1.75 | 1.79 |
| | | Pearson's $r$ | 0.73 | 0.58 | 0.42 | 0.51 | 0.42 |
| | contact | AUPRC | 0.0062 | 0.0060 | 0.0048 | 0.0057 | 0.0048 |
| | | AUROC | 0.5150 | 0.5157 | 0.5165 | 0.5150 | 0.5155 |

Table S4: Comparing accuracy and interpretability among various versions of DeepAffinity+ (DeepAffinity with regularized and supervised attentions) and DeepRelations. "cstr" indicates physical constraints imposed on attentions through regularization term $R_2(\cdot)$, whereas "sup" indicates supervised attentions through regularization term $R_3(\cdot)$. Best performances in each set are highlighted in boldface.

#### 2.3 Ablation study for DeepRelations

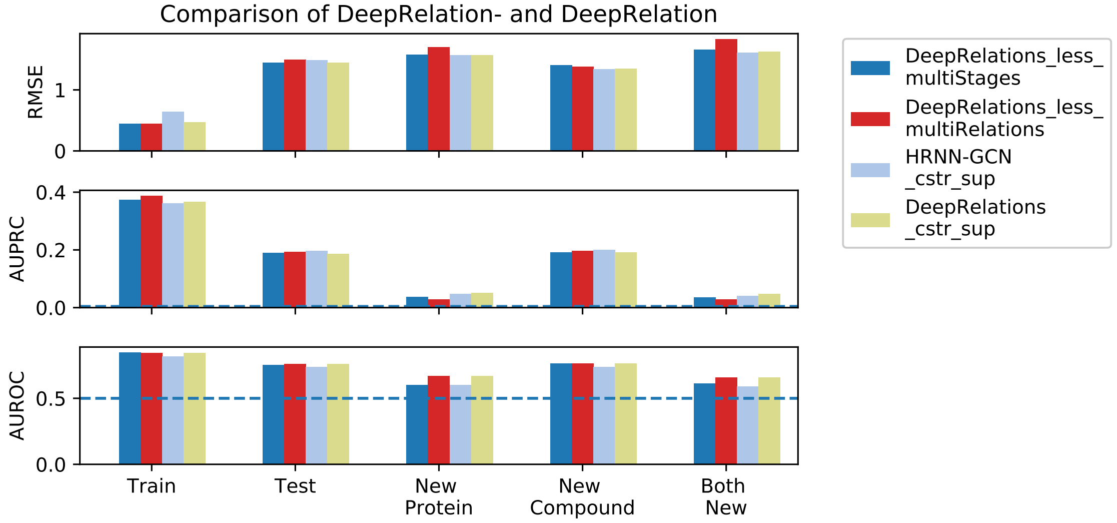


Figure S1: Comparing interpretability between DeepRelations and DeepRelations- (DeepRelations without multi-stage focusing, explicitly-modeled relations, or both).

#### 2.4 Randomization tests for affinity prediction

| | | Training | Test | New-Comp. | New-Prot. | Both-New |
|---|---|---|---|---|---|---|
| Random sampling training set | RMSE | 2.77 ± 0.03 | 2.82 ± 0.06 | 2.74 ± 0.07 | 2.72 ± 0.06 | 2.79 ± 0.11 |
| | Pearson's $r$ | 0.004 ± 0.019 | -0.002 ± 0.037 | -0.001 ± 0.041 | 0.006 ± 0.036 | 0.006 ± 0.068 |
| Random sampling each set | RMSE | 2.78 ± 0.03 | 2.85 ± 0.06 | 2.71 ± 0.07 | 2.51 ± 0.06 | 2.67 ± 0.13 |
| | Pearson's $r$ | -0.004 ± 0.021 | 0.003 ± 0.037 | -0.005 ± 0.043 | 0.004 ± 0.033 | 0.003 ± 0.081 |
| Mean of training | RMSE | 1.95 | 2.02 | 1.91 | 1.85 | 1.95 |
| Mean of each set | RMSE | 1.95 | 2.02 | 1.91 | 1.77 | 1.90 |
| Mean of all data | RMSE | 1.96 | 2.02 | 1.91 | 1.82 | 1.93 |

Table S5: Affinity prediction based on several random schemes.

| | | Permuted Training | Test | New-Comp. | New-Prot. | Both-New |
|---|---|---|---|---|---|---|
| HRNN Y randomization | RMSE | 0.56 ± 0.02 | 2.45 ± 0.06 | 2.39 ± 0.07 | 2.20 ± 0.15 | 2.25 ± 0.16 |
| | Pearson's $r$ | 0.960 ± 0.003 | 0.001 ± 0.054 | 0.001 ± 0.043 | -0.028 ± 0.084 | -0.10 ± 0.112 |
| DeepRelations Y randomization | RMSE | 1.03 ± 0.15 | 2.38 ± 0.16 | 2.29 ± 0.17 | 2.16 ± 0.13 | 2.24 ± 0.14 |
| | Pearson's $r$ | 0.817 ± 0.169 | -0.007 ± 0.063 | -0.006 ± 0.069 | -0.16 ± 0.078 | -0.031 ± 0.095 |

Table S6: Y randomization test for DeepAffinity+ and DeepRelations to validate their affinity prediction.

| | | Permuted Training | Test | New-Comp. | New-Prot. | Both-New |
|---|---|---|---|---|---|---|
| HRNN comp. randomization | RMSE | 0.55 ± 0.04 | 2.03 ± 0.04 | 1.95 ± 0.06 | 2.15 ± 0.15 | 2.22 ± 0.13 |
| | Pearson's $r$ | 0.962 ± 0.007 | 0.154 ± 0.032 | 0.243 ± 0.027 | 0.028 ± 0.097 | 0.017 ± 0.105 |
| DeepRelations comp. randomization | RMSE | 0.86 ± 0.17 | 2.06 ± 0.03 | 1.94 ± 0.12 | 2.20 ± 0.17 | 2.23 ± 0.16 |
| | Pearson's $r$ | 0.877 ± 0.128 | 0.135 ± 0.041 | 0.219 ± 0.039 | 0.012 ± 0.095 | 0.012 ± 0.113 |

Table S7: Compound randomization test for DeepAffinity+ and DeepRelations to validate their affinity prediction.

#### 2.5 Ensemble approach further improves affinity prediction

To further improve the accuracy of affinity prediction for DeepAffinity+ and DeepRelations, we have pursued an ensemble-learning approach and trained 50 models with different hyper-parameters for either DeepAffinity+ and DeepRelations (100 in total). Specifically, for either model, we used 5 different dropout ratios ({0.5, 0.6, 0.7, 0.8, 0.9}), 2 different $\lambda_{bind}$ ({100, 1000}) and 5 different amount of neurons for the 2 last fully connected layers ({(300, 300), (600, 300), (600, 600), (800, 800), (1000, 1000)}). In the end, we consider the ensemble of $5 \times 2 \times 5 = 50$ combinations for either DeepAffinity+ or DeepRelations as well as the ensemble of $50 + 50 = 100$ combinations. We report the results in Table S8 based on three metrics: RMSE, Pearson's $r$ and $R^2_{\text{pred}}$.

| | | Training | Test | New-Comp. | New-Prot. | Both-New |
|---|---|---|---|---|---|---|
| HRNN-GCN single | RMSE | 0.64 | 1.49 | 1.34 | 1.57 | 1.61 |
| | Pearson's $r$ | 0.95 | 0.68 | 0.73 | 0.45 | 0.51 |
| | $R^2_{\text{pred}}$ | 0.89 | 0.45 | 0.51 | 0.27 | 0.32 |
| DeepRelations single | RMSE | 0.47 | 1.45 | 1.35 | 1.57 | 1.63 |
| | Pearson's $r$ | 0.96 | 0.69 | 0.71 | 0.47 | 0.52 |
| | $R^2_{\text{pred}}$ | 0.92 | 0.48 | 0.50 | 0.27 | 0.30 |
| HRNN-GCN ensemble | RMSE | 0.19 | **1.29** | **1.20** | **1.50** | **1.60** |
| | Pearson's $r$ | 0.99 | **0.77** | **0.78** | **0.57** | **0.58** |
| | $R^2_{\text{pred}}$ | 0.99 | **0.59** | **0.60** | **0.34** | **0.34** |
| DeepRelations ensemble | RMSE | 0.29 | 1.35 | 1.27 | 1.59 | 1.65 |
| | Pearson's $r$ | 0.99 | 0.75 | 0.75 | 0.53 | 0.54 |
| | $R^2_{\text{pred}}$ | 0.98 | 0.55 | 0.56 | 0.26 | 0.28 |
| HRNN-GCN + DeepRelations ensemble | RMSE | 0.23 | 1.30 | 1.21 | 1.53 | **1.60** |
| | Pearson's $r$ | 0.99 | 0.76 | 0.77 | 0.56 | 0.57 |
| | $R^2_{\text{pred}}$ | 0.99 | 0.58 | **0.60** | 0.31 | 0.32 |

Table S8: Affinity prediction through ensembles of DeepAffinity+ or/and DeepRelations.

#### 2.6 Model Generalizability

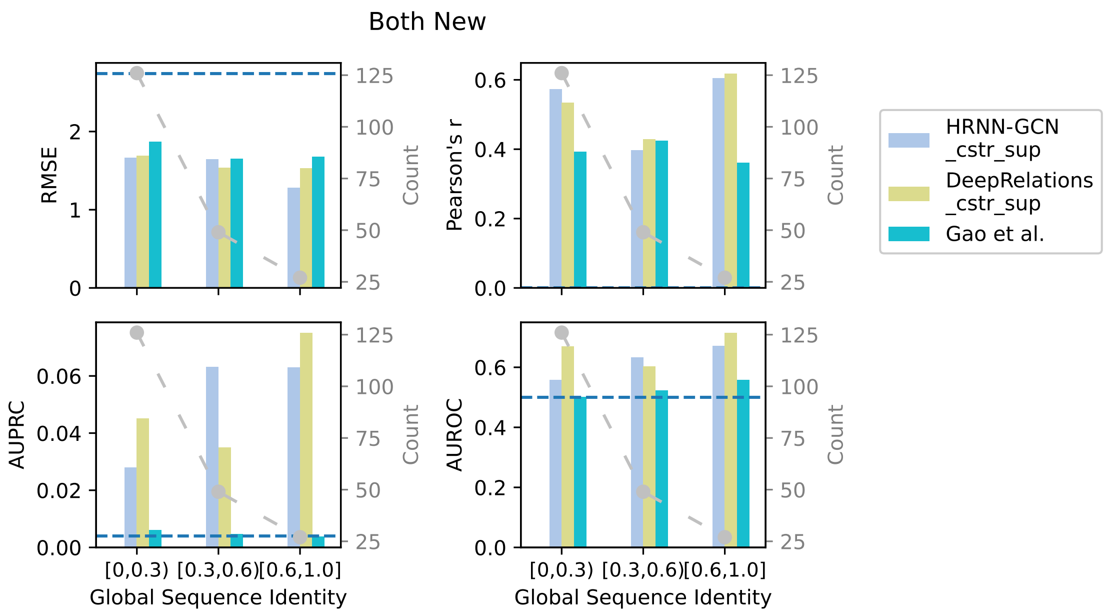


Figure S2: Comparing DeepAffinity+, DeepRelations, and Gao's method in the global protein generalizability of affinity prediction (RMSE and Pearson's $r$) and contact prediction (AUPRC and AUROC) in global sequence identity perspective for the both-new set. Cyan dashed lines indicate the performances of random predictors.

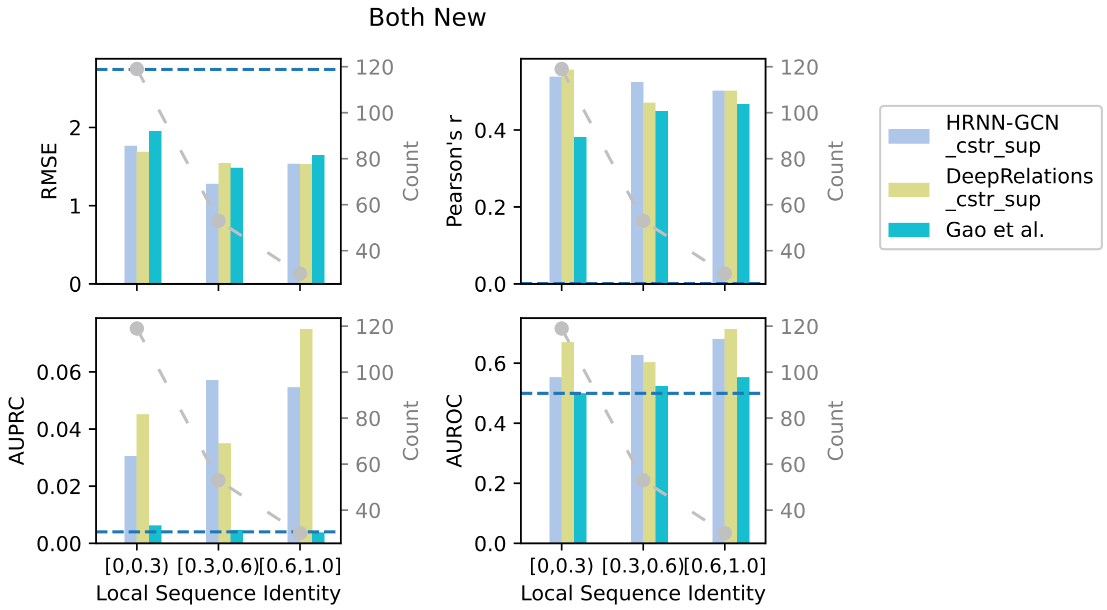


Figure S3: Comparing DeepAffinity+, DeepRelations, and Gao's method in the local protein (binding $k$-mer) generalizability of affinity prediction (RMSE and Pearson's $r$) and contact prediction (AUPRC and AUROC) in local sequence identity perspective for the both-new set.

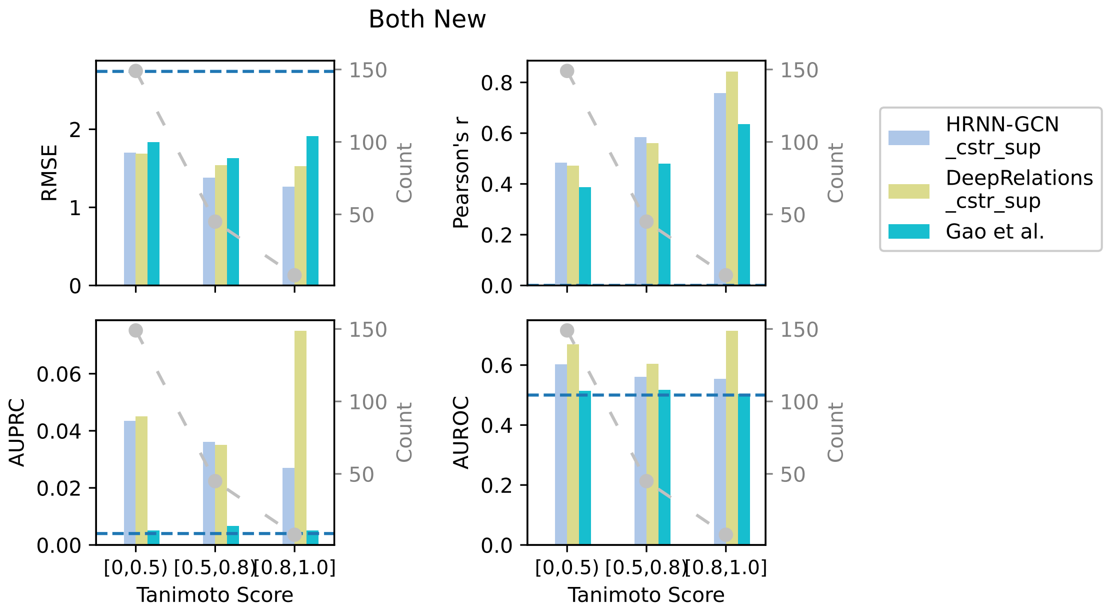


Figure S4: Comparing DeepAffinity+, DeepRelations, and Gao's method in the compound generalizability of affinity prediction (RMSE and Pearson's $r$) and contact prediction (AUPRC and AUROC) in tanimoto score perspective for the both-new set.

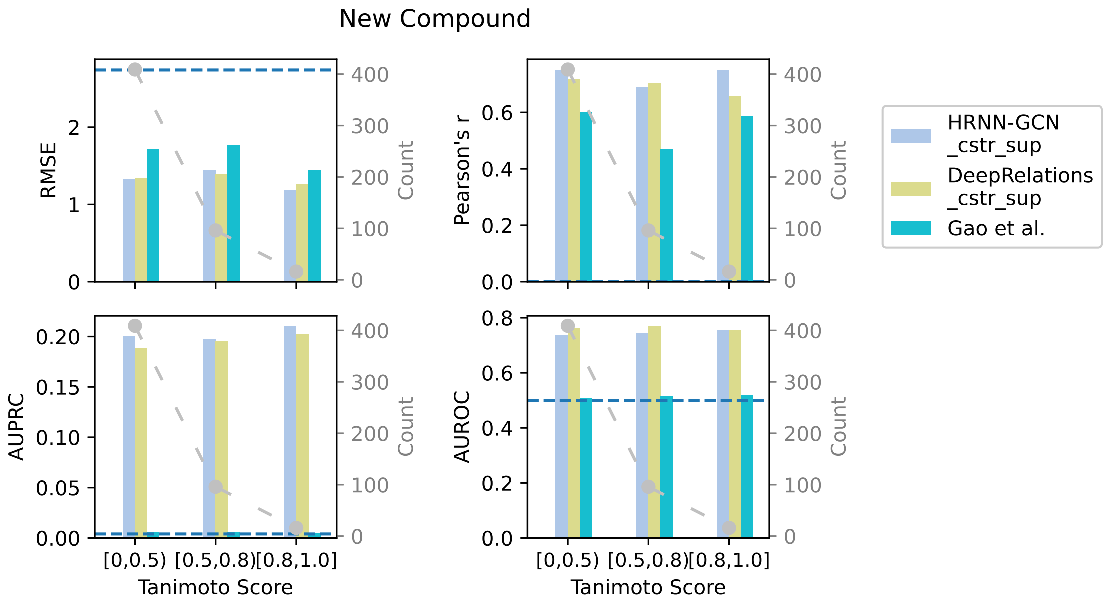


Figure S5: Comparing DeepAffinity+, DeepRelations, and Gao's method in the compound generalizability of affinity prediction (RMSE and Pearson's $r$) and contact prediction (AUPRC and AUROC) in tanimoto score perspective for the new compound set.

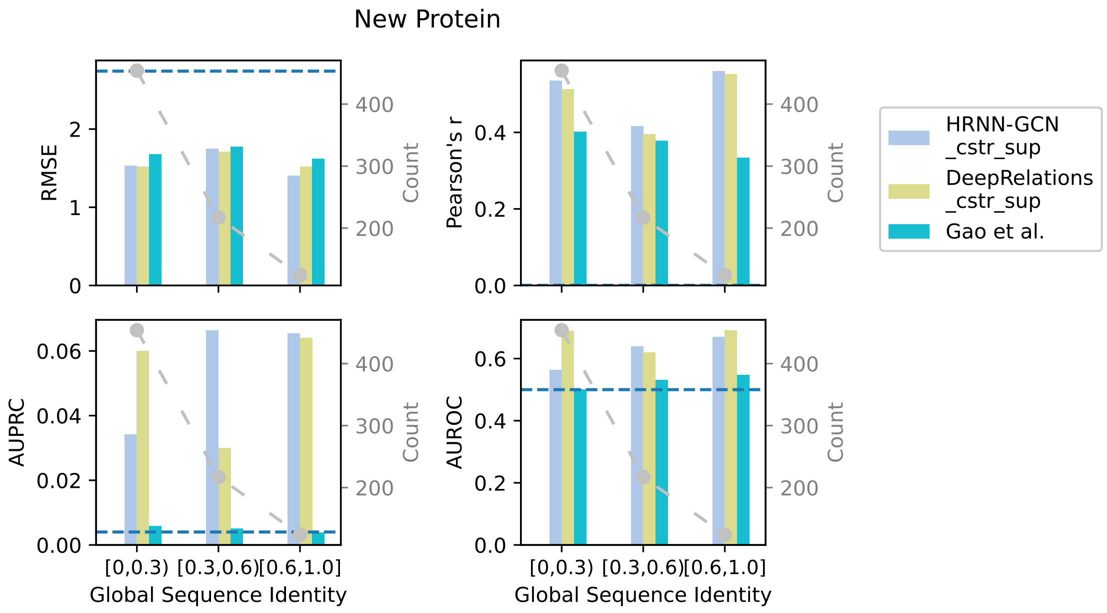


Figure S6: Comparing DeepAffinity+, DeepRelations, and Gao's method in the global protein generalizability of affinity prediction (RMSE and Pearson's $r$) and contact prediction (AUPRC and AUROC) in gloabl sequence identity perspective for the new protein set.

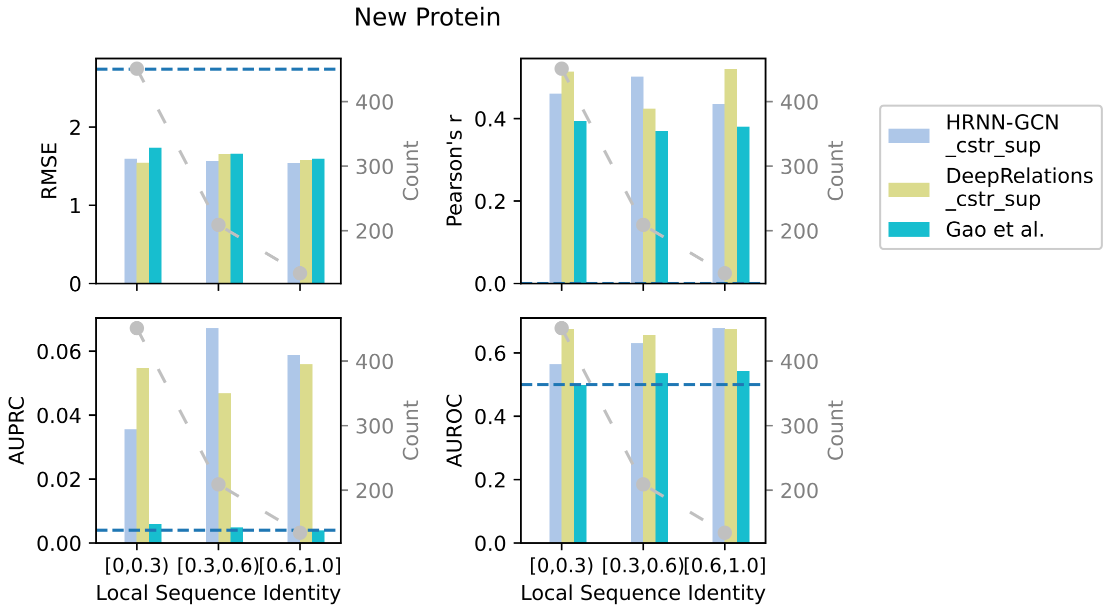


Figure S7: Comparing DeepAffinity+, DeepRelations, and Gao's method in the local protein (binding $k$-mer) generalizability of affinity prediction (RMSE and Pearson's $r$) and contact prediction (AUPRC and AUROC) for the new protein set.

| | | | Both New (seq identity) | Both New (k-mer identity) | Both New (Tanimoto) |
|---|---|---|---|---|---|
| HRNN-GCN _cstr_sup | affinity | RMSE | 1.66, 1.64, 1.28 | 1.76, 1.28, 1.53 | 1.69, 1.37, 1.26 |
| | | Pearson's $r$ | 0.57, 0.39, 0.60 | 0.54, 0.52, 0.50 | 0.48, 0.58, 0.75 |
| | contact | AUPRC | 0.0278, 0.0631, 0.0630 | 0.0306, 0.0571, 0.0545 | 0.0434, 0.0359, 0.0269 |
| | | AUROC | 0.5576, 0.6326, 0.6707 | 0.5520, 0.6269, 0.6814 | 0.6024, 0.5597, 0.5526 |
| DeepRelations _cstr_sup | affinity | RMSE | 1.69, 1.54, 1.53 | 1.73, 1.37, 1.68 | 1.71, 1.44, 1.23 |
| | | Pearson's $r$ | 0.53, 0.42, 0.61 | 0.56, 0.47, 0.50 | 0.47, 0.56, 0.84 |
| | contact | AUPRC | 0.045, 0.035, 0.075 | 0.043, 0.046, 0.065 | 0.050, 0.037, 0.041 |
| | | AUROC | 0.669, 0.603, 0.714 | 0.654, 0.672, 0.660 | 0.658, 0.668, 0.630 |
| Gao et al. | affinity | RMSE | 1.87, 1.65, 1.68 | 1.95, 1.49, 1.64 | 1.83, 1.62, 1.91 |
| | | Pearson's $r$ | 0.39, 0.42, 0.36 | 0.38, 0.44, 0.46 | 0.38, 0.48, 0.63 |
| | contact | AUPRC | 0.0060, 0.0045, 0.0037 | 0.0061, 0.0046, 0.0037 | 0.0050, 0.0066, 0.0050 |
| | | AUROC | 0.5016, 0.5224, 0.5568 | 0.4998, 0.5242, 0.5525 | 0.5137, 0.5168, 0.5041 |

| | | | New Prot. (seq identity) | New Prot. (k-mer identity) | New Compound (Tanimoto) |
|---|---|---|---|---|---|
| HRNN-GCN _cstr_sup | affinity | RMSE | 1.53, 1.75, 1.40 | 1.59,1.56,1.53 | 1.32, 1.44, 1.18 |
| | | Pearson's $r$ | 0.53, 0.41, 0.56 | 0.46,0.50,0.43 | 0.74, 0.69, 0.75 |
| | contact | AUPRC | 0.0340, 0.0662, 0.0654 | 0.0355, 0.0671, 0.0588 | 0.2001, 0.1972, 0.2101 |
| | | AUROC | 0.5627, 0.6388, 0.6695 | 0.5642, 0.6290, 0.6764 | 0.7361, 0.7435, 0.7539 |
| DeepRelations _cstr_sup | affinity | RMSE | 1.52, 1.71, 1.52 | 1.54, 1.65, 1.57 | 1.34, 1.39, 1.26 |
| | | Pearson's $r$ | 0.51, 0.39, 0.55 | 0.51, 0.42, 0.52 | 0.71, 0.70, 0.65 |
| | contact | AUPRC | 0.060, 0.030, 0.064 | 0.054, 0.046, 0.055 | 0.189, 0.196, 0.202 |
| | | AUROC | 0.688, 0.619, 0.69 | 0.675, 0.656, 0.673 | 0.763, 0.769, 0.757 |
| Gao et al. | affinity | RMSE | 1.67, 1.77, 1.62 | 1.73, 1.66, 1.59 | 1.72, 1.76, 1.44 |
| | | Pearson's $r$ | 0.40, 0.37, 0.33 | 0.39, 0.36, 0.37 | 0.60, 0.46, 0.58 |
| | contact | AUPRC | 0.0058, 0.0050, 0.0038 | 0.0059, 0.0048, 0.0038 | 0.0060, 0.0061, 0.0052 |
| | | AUROC | 0.5002, 0.5309, 0.5469 | 0.4987, 0.5356, 0.5431 | 0.5097, 0.5145, 0.5192 |

Table S9: Comparing DeepAffinity+, DeepRelations, and Gao's method in the generalizability of affinity prediction (RMSE and Pearson's $r$) and contact prediction (AUPRC and AUROC) to molecules unlike training data. Each cell includes three numbers for increasingly similar proteins or compounds to training data (global sequence or binding $k$-mer identity below 30%, between 30% and 60%, and above 60%; Tanimoto score below 0.5, between 0.5 and 0.8, and above 0.8).

The global and local sequence identities are defined as follows. We denote the $i$-th reference sequence in the training set as $S_i^{\mathrm{r}}$ and the test sequence in the new-protein or both-new set as $S^{\mathrm{t}}$. Then the global sequence identity for a test sequence $S^{\mathrm{t}}$ is defined as

$$\mathrm{ID}_{\mathrm{global}}(S_t) = \max_{i \in \mathrm{trainingSet}} \mathrm{SeqID}(S^{\mathrm{t}}, S_i^{\mathrm{r}}),$$

in which $\mathrm{SeqID}(S_t, S_{ri})$ denotes the sequence identity.
Similarly, the local identity means the sequence identity at the binding $k$-mer level. Here we only consider binding $k$-mers with at least two binding residues. Only around 4 binding-site residues (less than 10%) are present in an average $k$-mer ($k$=40) and around 3 binding $k$-mers are found in an average protein. The local, binding $k$-mer sequence identity for a given test sequence is simply the maximum binding $k$-mer level sequence identity averaged over all binding $k$-mers of the sequence.

#### 2.7 Distance patterns of top-10 predicted residue-atom contacts

| | Test | New Protein |
|---|---|---|
| HRNN-GCN_cstr_sup | 0.3986, 0.1862, 0.0729, 0.0556 | 0.1768, 0.2245, 0.0608, 0.0416 |
| HRNN-GIN_cstr_sup | 0.1170, 0.1049, 0.0776, 0.0783 | 0.0476, 0.0793 ,0.0420, 0.0534 |
| DeepRelations_cstr_sup | 0.3734, 0.1935, 0.0840, 0.0637 | 0.1746, 0.2367, 0.0958, 0.0794 |
| Gao et al. | 0.0055, 0.0179 , 0.0269, 0.0446 | 0.0050, 0.0134 ,0.0275 ,0.0671 |
| | New Compound | Both New |
| HRNN-GCN_cstr_sup | 0.4030 ,0.2099, 0.0823, 0.0612 | 0.1559, 0.1920 ,0.0539, 0.0539 |
| HRNN-GIN_cstr_sup | 0.1201, 0.1186, 0.0852, 0.0654 | 0.0534, 0.0782 ,0.0475 ,0.0594 |
| DeepRelations_cstr_sup | 0.3738, 0.2115, 0.0902, 0.0545 | 0.1551, 0.2131, 0.0770 ,0.0853 |
| Gao et al. | 0.0051 ,0.0149, 0.0239, 0.0395 | 0.0039, 0.0193 ,0.0282, 0.0504 |

Table S10: Distributions of top-10 binding sites predicted by DeepAffinity+, DeepRelations, and Gao et al.. Four fractions in each cell correspond to various distance ranges in the order of $\leq 4$, $[4, 6)$, $[6, 8)$, and $[8, 10)$ (unit: Å).

#### 2.8 Binding site prediction

| | | Train | Test | New Protein | New Compound | Both New |
|---|---|---|---|---|---|---|
| HRNN-GCN _cstr_sup | AUPRC | 0.5830 | 0.4216 | 0.1698 | 0.4314 | 0.1565 |
| | AUROC | 0.8576 | 0.7633 | 0.6493 | 0.7822 | 0.6518 |
| HRNN-GIN _cstr_sup | AUPRC | 0.1536 | 0.1252 | 0.0611 | 0.1296 | 0.0676 |
| | AUROC | 0.5728 | 0.5648 | 0.4865 | 0.5615 | 0.4930 |
| DeepRelations _cstr_sup | AUPRC | 0.594 | 0.431 | 0.208 | 0.440 | 0.195 |
| | AUROC | 0.865 | 0.769 | 0.726 | 0.789 | 0.720 |
| Gao et al. | AUPRC | 0.0545 | 0.0543 | 0.0495 | 0.0538 | 0.0496 |
| | AUROC | 0.5012 | 0.4979 | 0.4821 | 0.5051 | 0.4874 |

Table S11: Binding site prediction results for DeepAffinity+, DeepRelations, and Gao et al.'s method (adapted to affinity prediction).

#### 2.9 Case study: Affinity prediction for CA2 compounds against non-CA2 proteins

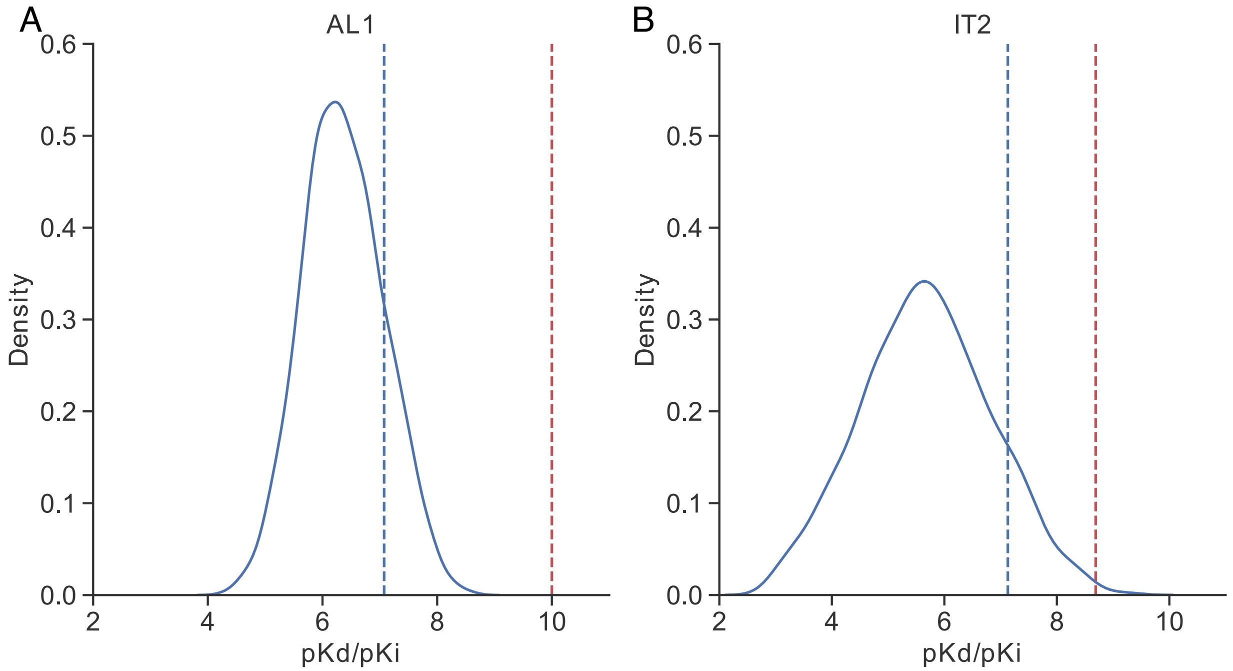


Figure S8: Probability distributions of DeepAffinity+ affinity predictions for AL1 and IT2 compounds against non-CA2 proteins. Red and blue dashed lines indicate actual and predicted affinity to the known target CA2.

#### 2.10 Affinity and contact prediction of DeepRelations with actual protein residue-residue contact maps

For the true residue-contact maps, we used 8Å-cutoff for $C_\beta$–$C_\beta$ distances in known structures (bound for now). The maps can be incomplete due to missing residues in PDB.

| | | | Training | Test | New Comp | New Prot | Both New |
|---|---|---|---|---|---|---|---|
| DeepRelation with predicted residue-contact maps | affinity | RMSE | 0.47 | **1.45** | **1.35** | **1.57** | **1.63** |
| | | Pearson's $r$ | 0.96 | 0.69 | **0.71** | **0.47** | **0.52** |
| | contact | AUPRC | 0.367 | 0.187 | 0.191 | **0.052** | **0.047** |
| | | AUROC | 0.845 | **0.760** | 0.764 | **0.669** | **0.659** |
| DeepRelation with true residue-contact maps | affinity | RMSE | 0.46 | 1.49 | 1.45 | 1.64 | 1.72 |
| | | Pearson's $r$ | 0.97 | **0.70** | 0.69 | 0.44 | 0.47 |
| | contact | AUPRC | 0.432 | **0.204** | **0.206** | 0.026 | 0.028 |
| | | AUROC | 0.858 | 0.754 | **0.766** | 0.652 | 0.63 |

Table S12: Comparing DeepRelations performances with predicted or actual protein residue-contact maps.

The performances with predicted and actual residue-contact maps are somewhat close. We conjecture that contact map prediction might be quite accurate for our dataset that contains proteins of known structures.

#### 2.11 SAR

| Method | Kendall's $\tau$ | Spearman's $\rho$ |
|---|---|---|
| 87mci | 0.71 | 0.86 |
| 7bi2k | 0.6 | 0.75 |
| yghq5 | 0.56 | 0.74 |
| 5rtw4 | 0.31 | 0.42 |
| a6kw3 | 0.16 | 0.25 |
| vshma | 0.13 | 0.32 |
| 5jfzy | 0.05 | 0.05 |
| 5ywmk | 0.02 | 0.00 |
| ngsyu | 0.02 | 0.07 |
| kqhj5 | 0.02 | 0.04 |
| tdvzq | -0.02 | 0.02 |
| ispmq | -0.09 | -0.08 |
| 8xf7u | -0.16 | -0.30 |
| ukjcf | -0.20 | -0.27 |
| a8eqr | -0.20 | -0.26 |
| yqoad | -0.31 | -0.50 |
| ha84u | -0.31 | -0.40 |
| gvzji | -0.56 | -0.7 |
| Gao et al. | -0.42 | -0.54 |
| HRNN-GCN (ours) | -0.36 | -0.47 |
| DeepRelations (ours) | 0.15 | 0.21 |

Table S13: Comparison of our structure-free methods and Gao et al. with structure-based models (marked in receipt IDs) that participated in D3R grand challenge 3 for JAK2.

| Method | Kendall's $\tau$ | Spearman's $\rho$ |
|---|---|---|
| uuihe | 0.57 | 0.76 |
| y7qxv | 0.57 | 0.74 |
| xpmn7 | 0.50 | 0.67 |
| h6qgu | 0.36 | 0.55 |
| 7d5vc | 0.29 | 0.38 |
| mey8v | 0.21 | 0.33 |
| hjy28 | 0.21 | 0.19 |
| jkqbh | 0.07 | 0.14 |
| 0dnju | 0.07 | 0.17 |
| bu5xc | 0 | 0.05 |
| pox3b | 0.00 | -0.05 |
| fn2qt | -0.07 | -0.05 |
| km744 | -0.07 | -0.17 |
| th0hn | -0.07 | 0.00 |
| 0fdt7 | -0.29 | -0.33 |
| 6685v | -0.43 | -0.60 |
| yfg86 | -0.5 | -0.67 |
| wg3ed | -0.57 | -0.69 |
| Gao et al. | 0.60 | 0.74 |
| HRNN-GCN (ours) | 0.65 | 0.79 |
| DeepRelations (ours) | 0.61 | 0.72 |

Table S14: Comparison of our structure-free methods and Gao et al. with structure-based models (marked in receipt IDs) that participated in D3R grand challenge 3 for TIE2.

| **Receipt ID** | Kendall's $\tau$ | Spearman's $\rho$ | Pearson's $r$ | $R^2$ |
|---|---|---|---|---|
| 7bi2k | 0.41 | 0.59 | 0.57 | 0.32 |
| 87mci | 0.36 | 0.52 | 0.49 | 0.24 |
| yghq5 | 0.28 | 0.41 | 0.36 | 0.13 |
| 5rtw4 | 0.21 | 0.32 | 0.27 | 0.07 |
| vshma | 0.1 | 0.18 | 0.16 | 0.03 |
| 5ywmk | 0.07 | 0.1 | 0.08 | 0.01 |
| kqhj5 | 0.07 | 0.09 | 0.09 | 0.01 |
| 5jfzy | 0.06 | 0.1 | 0.11 | 0.01 |
| a6kw3 | 0.02 | 0.04 | 0.04 | 0.0 |
| ngsyu | -0.02 | -0.03 | -0.1 | 0.01 |
| ukjcf | -0.04 | -0.06 | -0.03 | 0.0 |
| tdvzq | -0.05 | -0.05 | -0.09 | 0.01 |
| ispmq | -0.06 | -0.08 | -0.07 | 0.0 |
| a8eqr | -0.09 | -0.13 | -0.14 | 0.02 |
| ha84u | -0.13 | -0.19 | -0.18 | 0.03 |
| yqoad | -0.2 | -0.33 | -0.23 | 0.05 |
| 8xf7u | -0.21 | -0.32 | -0.31 | 0.09 |
| gvzji | -0.25 | -0.36 | -0.39 | 0.15 |
| Gao et al. | 0.05 | 0.08 | 0.1 | 0.01 |
| HRNN-GCN (ours) | -0.24 | -0.3 | -0.29 | 0.08 |
| DeepRelation (ours) $\Delta p\hat{K}_d$ | 0.16 | 0.26 | 0.22 | 0.05 |
| DeepRelation (ours) $\Delta p\hat{K}_d^R$ | 0.26 | 0.41 | 0.36 | 0.13 |

Table S15: Comparison of our models with structure-based models participated in D3R grand challenge 3 for JAK2 for $\Delta y$ prediction

| JAK2 (Subchallenge 3): $\Delta pK_d$ prediction | | | | | |
|---|---|---|---|---|---|
| Ranking[a] | Method(s) | $\tau$ | $\rho$ | $r$ | $R^2$ |
| 1–3 | 3 structure-based methods in D3R | $0.28 \sim 0.41$ | $0.41 \sim 0.59$ | $0.36 \sim 0.57$ | $0.13 \sim 0.32$ |
| 4 | Structure-free DeepRelations (ours) $\Delta p\hat{K}_d^R$ | 0.26 | 0.41 | 0.36 | 0.13 |
| 5 | 1 structure-based method in D3R | 0.21 | 0.32 | 0.27 | 0.07 |
| 6 | Structure-free DeepRelations (ours) $\Delta p\hat{K}_d$ | 0.16 | 0.26 | 0.22 | 0.05 |
| 7–10 | 4 structure-based methods in D3R | $0.06 \sim 0.1$ | $0.1 \sim 0.18$ | $0.11 \sim 0.16$ | $0.01 \sim 0.03$ |
| 11 | Structure-free Gao et al. | 0.05 | 0.08 | 0.1 | 0.01 |
| 12–20 | 9 structure-based methods in D3R | $-0.21 \sim 0.02$ | $-0.33 \sim 0.04$ | $-0.31 \sim 0.04$ | $0.00 \sim 0.09$ |
| 21 | Structure-free DeepAffinity+ | -0.24 | -0.3 | -0.29 | 0.08 |
| 22 | 1 structure-based method in D3R | -0.25 | -0.36 | -0.39 | 0.13 |

[a] The 18 structure-based methods participated in the D3R subchallenges and were assessed officially. The 4 structure-free methods were assessed *post hoc*.

Table S16: Summary of $\Delta pK_d$ scoring performances among three structure-free methods (including our DeepAffinity+ and two versions of DeepRelations and eighteen structure-based methods.

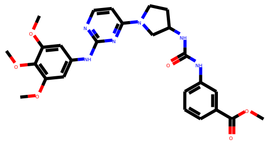

comp1 ,pKd: 6.47 ,pred: 8.02

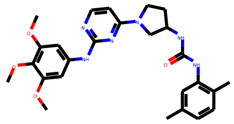

comp2 ,pKd: 5.0 ,pred: 7.6

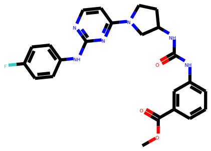

comp3 ,pKd: 5.0 ,pred: 7.01

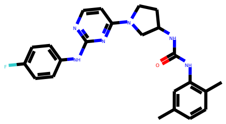

comp4 ,pKd: 5.0 ,pred: 6.78

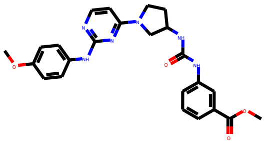

comp5 ,pKd: 5.0 ,pred: 7.62

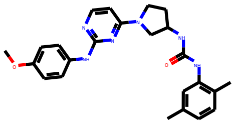

comp6 ,pKd: 5.0 ,pred: 7.25

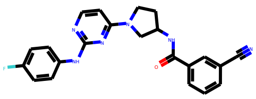

comp7 ,pKd: 5.24 ,pred: 6.79

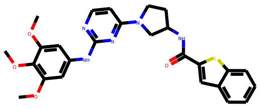

comp8 ,pKd: 6.15 ,pred: 7.38

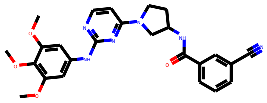

comp9 ,pKd: 6.74 ,pred: 7.42

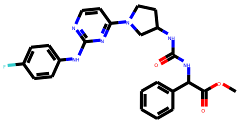

comp10 ,pKd: 5.11 ,pred: 6.89

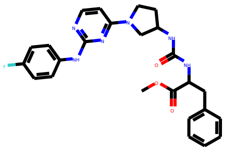

comp11 ,pKd: 5.72 ,pred: 6.74

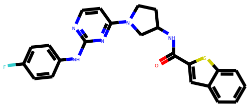

comp12 ,pKd: 5.0 ,pred: 6.52

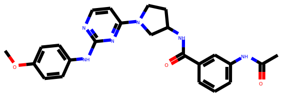

comp13 ,pKd: 5.62 ,pred: 6.65

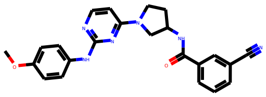

comp14 ,pKd: 5.33 ,pred: 7.09

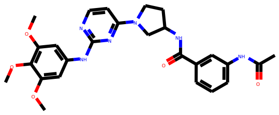

comp15 ,pKd: 7.27 ,pred: 7.02

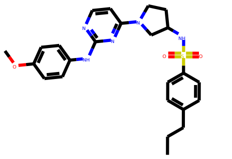

comp16 ,pKd: 6.17 ,pred: 7.43

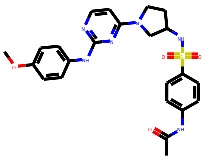

comp17 ,pKd: 7.28 ,pred: 7.12

Figure S9: Compounds in JAK2's SAR with actual and DeepRelations-predicted affinities.

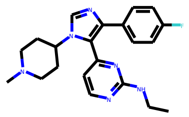

comp1 ,pKd: 5.0 ,pred: 8.2

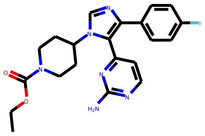

comp2 ,pKd: 5.0 ,pred: 7.91

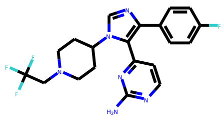

comp3 ,pKd: 5.0 ,pred: 7.81

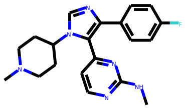

comp4 ,pKd: 5.0 ,pred: 7.64

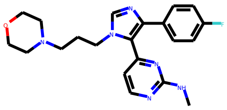

comp5 ,pKd: 5.0 ,pred: 7.58

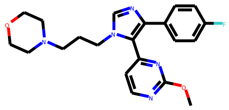

comp6 ,pKd: 5.0 ,pred: 7.57

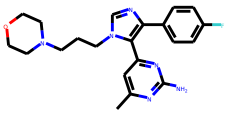

comp7 ,pKd: 5.0 ,pred: 7.5

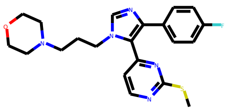

comp8 ,pKd: 5.0 ,pred: 7.6

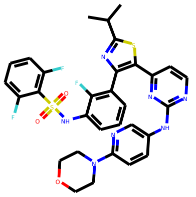

comp9 ,pKd: 8.47 ,pred: 10.25

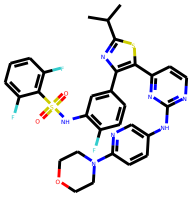

comp10 ,pKd: 7.62 ,pred: 10.41

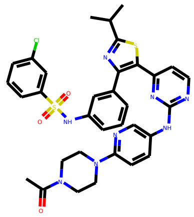

comp11 ,pKd: 7.03 ,pred: 9.45

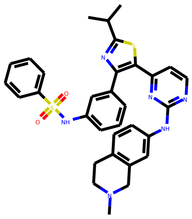

comp12 ,pKd: 7.04 ,pred: 8.35

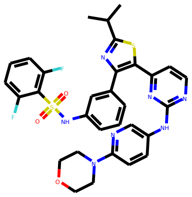

comp13 ,pKd: 7.57 ,pred: 10.0

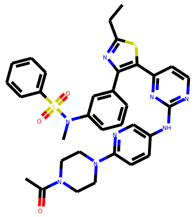

comp14 ,pKd: 5.49 ,pred: 8.62

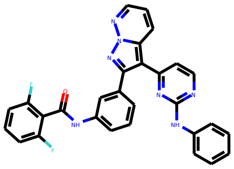

comp15 ,pKd: 5.6 ,pred: 8.65

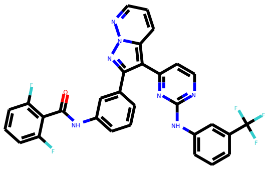

comp16 ,pKd: 5.0 ,pred: 9.54

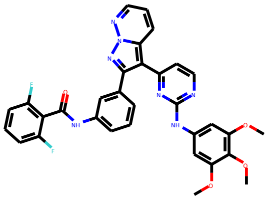

comp17 ,pKd: 6.7 ,pred: 9.21

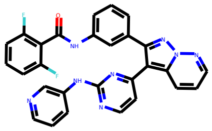

comp18 ,pKd: 5.0 ,pred: 9.51

Figure S10: Compounds in TIE2's SAR with actual and DeepRelations-predicted affinities.

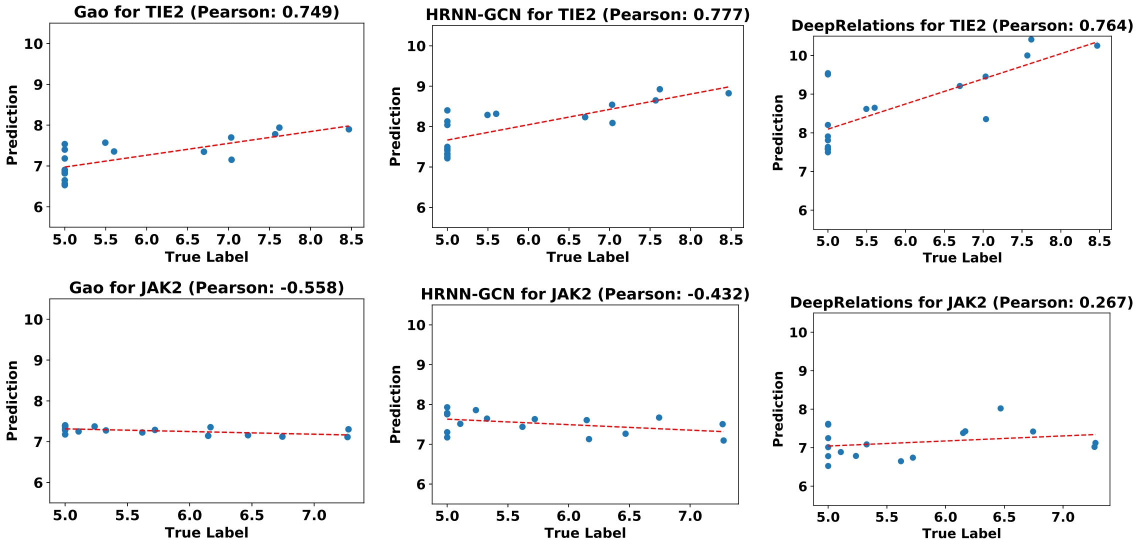


Figure S11: Comparison of predicted affinity versus true affinity for different methods for SAR cases.

#### 2.12 Combinatorial lead optimization

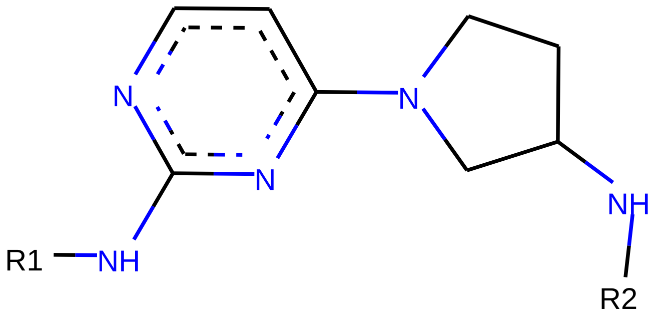


Figure S12: Diagram for the scaffold and two functional groups of the JAK2 compounds.

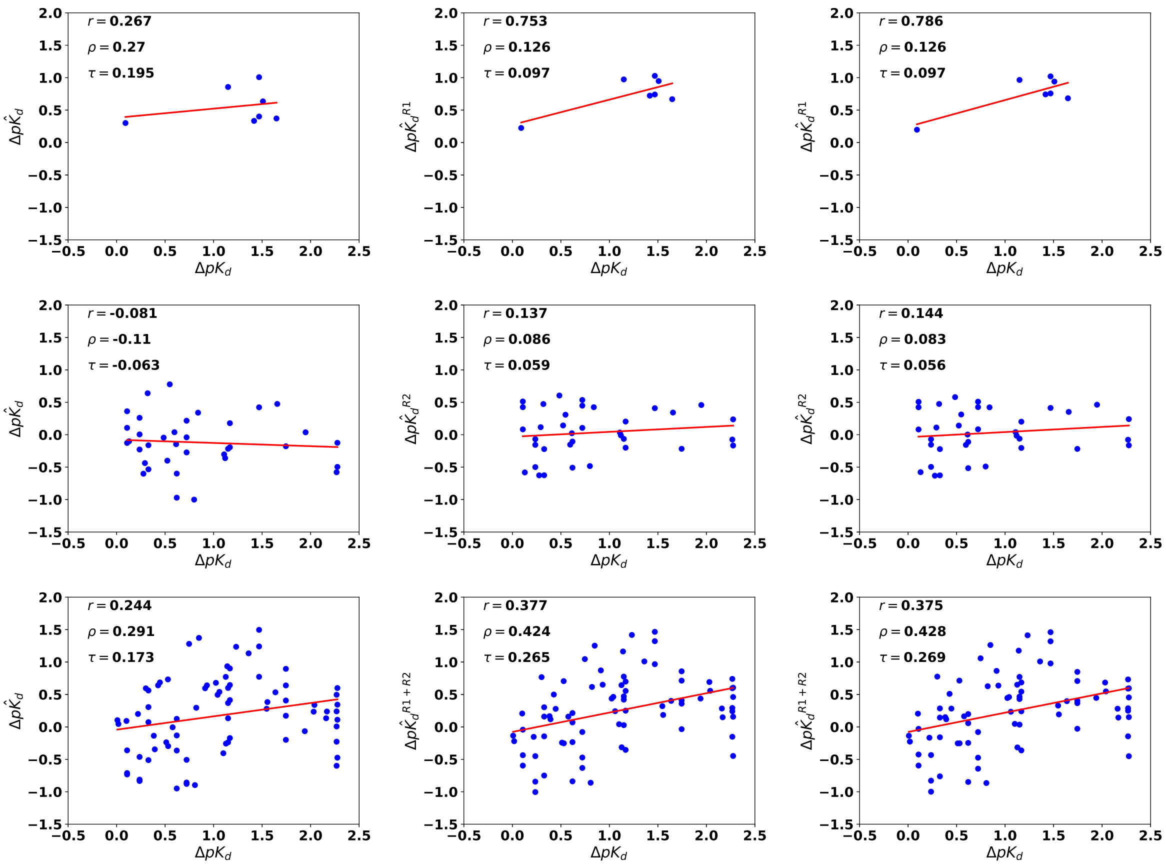


Figure S13: Comparison of true affinity changes versus various DeepRelations-based predictions, when JAK2 compounds are changed by substituting functional groups. The first row is for R1 substitution only, second for R2 substitution only and third for both substitutions. The first column is for prediction based on predicted affinity only, the second is based on decomposition (substituent group for the compound and all residues for the protein), and the third column is based on decomposition (substituent group for the compound and binding-site residues for the protein).

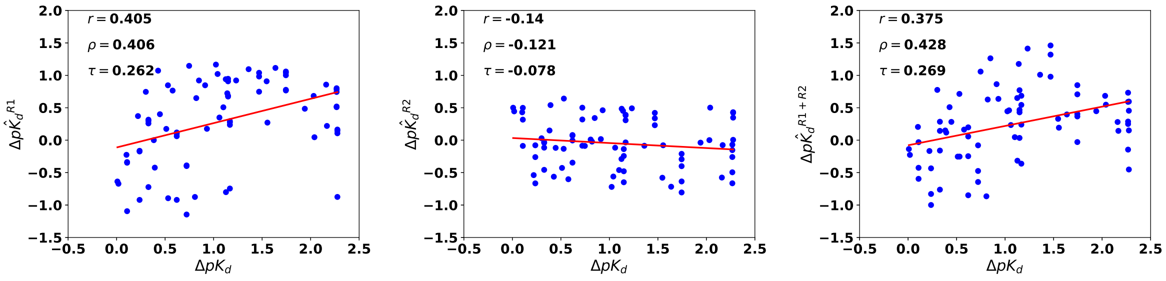


Figure S14: For both R1+R2 substitutions, we compare the contribution of R1 (left), R2 (middle) and R1+R2 (right).
